# Glioblastoma and other intracranial tumors elicit systemic sympathetic hyperactivity that limits immunotherapeutic responses

**DOI:** 10.1101/2023.11.02.565368

**Authors:** Selena J. Lorrey, Lucas P. Wachsmuth, John B. Finlay, Corey Neff, Katayoun Ayasoufi, Jessica Waibl Polania, Alexandra Hoyt-Miggelbrink, Mackenzie Price, Lindsay Rein, Samuel Dell, Rachael Reesman, Xiuyu Cui, Amelia Hawley, Emily Lerner, Daniel Wilkinson, Ethan Srinivasan, Kyle M. Walsh, Anoop Patel, Quinn T. Ostrom, Peter E. Fecci

**Author notes:** contributed equally (Selena J. Lorrey, Lucas P. Wachsmuth).

## Abstract

Intracranial tumors present unique challenges for immunotherapy. These can include both local and systemic modes of immune suppression whose mechanistic underpinnings are incompletely understood. Here, we reveal that tumors harbored intracranially elicit systemic increases to circulating catecholamine levels, with the resultant chronic sympathetic hyperactivity driving T cell dysfunction and limiting immunotherapeutic success. Conversely, treatment with β-adrenergic blockade increases NF-κB activity in immune cells, restores T cell polyfunctionality, modifies the tumor microenvironment, and licenses immune-based therapies in murine models of glioblastoma (GBM) to extend survival. Extended survival is also observed in GBM patients having received β-adrenergic blockade, as well as in patients with melanoma and lung cancer brain metastases who received β-blockade alongside concomitant immune checkpoint inhibition. While β-blockade also impacts outcomes in the setting of extracranial disease, the benefits are especially pronounced in patients harboring intracranial disease burdens. These data suggest that sympathetic hyperactivity facilitates systemic immune dysfunction in the setting of intracranial tumors, specifically and advance a role for β-adrenergic blockade in licensing immunotherapeutic responses within the intracranial compartment.

## INTRODUCTION

Glioblastoma (GBM) is the most common primary brain malignancy and remains nearly uniformly lethal. While modern immunotherapeutic approaches enjoy success against a variety of solid tumors, GBM persists as a notable exception^1–3^. Although GBM remains generally confined to the intracranial compartment, it inflicts a remarkably broad range of systemic immune impairments that contribute to treatment failures. Various modes of T cell dysfunction are prevalent and serve to hinder the antitumor response^4–27^. Additional peripheral immune derangements include lymphopenia, lymphoid organ atrophy, and sequestration of T cells in the bone marrow^4,20–22,26–28^. Many of these derangements are not particular to GBM, but instead accompany various tumors if and when these become situated within the intracranial compartment. Lung, melanoma, and breast cancers can all elicit comparable immune dysfunction, for instance, but only when harbored within the brain^20^.

Herein we report that intracranial tumors (but not their peripheral counterparts), elicit dramatic rises in systemic catecholamine levels in both patients and mice, which in turn drive loss of critical T cell functions. This remains true regardless of tumor histology Catecholamine infusions into naïve mice are sufficient to recapitulate the same deficiencies, as are specific β-adrenergic agonists (β2 > β1). β2-adrenergic receptors prove the predominant adrenergic receptor on lymphocytes, and their signaling has been shown to suppress immune cell function via inhibition of nuclear factor kappa B (NF-κB) activity^29–34^. Accordingly, we find that treatment with β-adrenergic receptor blockade increases NF-κB activity in immune cells, restores T cell polyfunctionality, modifies the tumor microenvironment (TME), and licenses immune-based therapies in murine models of GBM to newly elicit long-term survival.

Large scale analysis of Medicare data also reveals extended survival in GBM patients having received β-blockade, irrespective of indication. Similar survival benefits are seen in patients with melanoma and lung cancer brain metastases who received β-blockade alongside concomitant immune checkpoint inhibition (ICI), rather than ICI alone. While β-blockade impacts outcomes in the setting of extracranial disease as well, the benefits are especially pronounced in patients harboring intracranial disease burdens. These data suggest critical roles for increased adrenergic activity in facilitating systemic immune dysfunction in the setting of intracranial tumors and advance a particular role for β-adrenergic blockade in licensing immunotherapeutic responses within the intracranial compartment.

## RESULTS

### Tumors harbored intracranially elicit rises in circulating catecholamine levels

Investigating a role for the sympathetic nervous system (SNS) in mediating the systemic immune dysfunction observed in patients and mice with GBM, we collected sera and spleens from immunocompetent C57BL/6 mice bearing syngeneic intracranial CT2A glioma (CT2A). This model is widely used and is generally accepted as the syngeneic model that most closely recapitulates the immune microenvironment in human GBM^35^. Systemic catecholamine levels from tumor-bearing mice were compared to those in mice instead receiving sham tumor implantations **(Fig. 1A).** Catecholamine levels were elevated in both the serum **(Fig. 1B)** and spleens **(Fig. 1C)** of mice bearing intracranial CT2A, but corresponding increases were not observed in the tumor **(Extended Data Fig. 1 A,B).** Systemic catecholamine rises were observed once a particular tumor size threshold was reached, but further increases in tumor size did not correlate with similar increases in catecholamine levels (**Extended Data Fig. 1 C,D).** Additionally, we assessed corticosterone levels in murine serum and found mixed results (data not shown).

**Figure 1.**
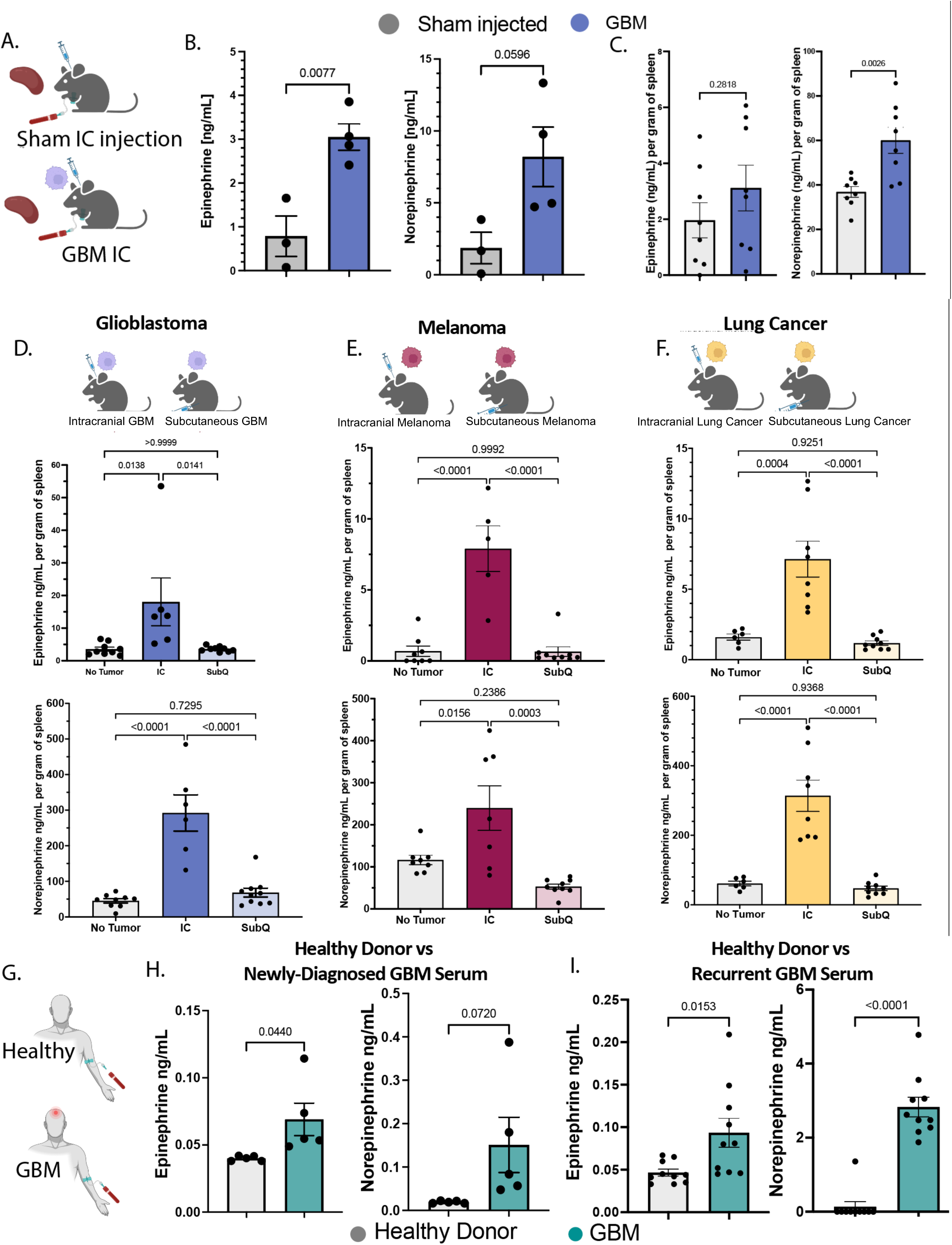
Patients and mice with intracranial tumors exhibit elevated systemic catecholamine levels. A. Schematic of serum collected from mice with intracranial glioblastoma or a sham intracranial injection (methylcellulose and PBS vehicle alone). B. Serum epinephrine and norepinephrine levels in mice with intracranial glioblastoma compared with a sham intracranial injection (n≥3 per group). C. Splenic epinephrine and norepinephrine levels, normalized to spleen weight, in mice with intracranial glioblastoma compared to those with a sham intracranial injection (n=8 per group). D. Splenic epinephrine and norepinephrine levels in mice with no tumor, intracranial glioblastoma or subcutaneous glioblastoma (n≥6 per group). E. Splenic epinephrine and norepinephrine levels in mice with no tumor, intracranial melanoma or subcutaneous melanoma (n≥7 per group). F. Splenic epinephrine and norepinephrine levels in mice with no tumor, intracranial lung cancer and subcutaneous lung cancer (n≥6 per group). G. Schematic of serum collection from patients with glioblastoma or healthy controls H. Serum epinephrine and norepinephrine levels in patients with newly-diagnosed glioblastoma compared with healthy controls (n=5 per group). I. Serum epinephrine and norepinephrine levels in patients with recurrent glioblastoma compared with healthy controls (n=10 per group). The p values for B, C, H and I were calculated using unpaired t-tests and the p values in D, E, and F were calculated using one-way ANOVAs and Tukey’s multiple comparisons tests. Error bars denote the standard error of the mean (SEM). Data are representative of individual experiments.

Intrigued, we examined whether rising systemic catecholamine levels might be: 1) particular to GBM; 2) reflective of tumors harbored within the intracranial compartment; or 3) a feature of cancer more broadly. CT2A tumors were injected into either the brains (intracranial, IC) or the flanks (subcutaneous, SubQ) of C57B/6 mice. Systemic elevations in epinephrine and norepinephrine were observed with IC CT2A only, suggesting a phenomenon not particular to GBM, but rather to the intracranial compartment **(Fig. 1D)**. Similar findings accompanied intracranial adaptations of Lewis Lung carcinoma (LLC) and B16 melanoma, but not their peripherally-introduced counterparts, further evidence of an intracranial tumor phenomenon **(Figs. 1E,F)**.

Having observed the above in murine models, we sought to determine whether similar findings might characterize patients. Thus, we examined serum catecholamine levels in patients with either newly diagnosed (n=5) or recurrent (n=10) GBM, as well as in sex-matched healthy controls **(Fig. 1G)**. GBM patients from each cohort exhibited significant elevations in circulating levels of both epinephrine and norepinephrine (**Fig. 1H, I)**. Similar rises were not observed in levels of stress hormones (data not shown).

### β2-adrenergic receptors predominate within the intracranial TME, as well as on circulating immune cells

Epinephrine and norepinephrine signal through α- and β-adrenergic receptors. Given the observed rise in catecholamine levels, we examined the relative prevalence of the various α- and β-adrenergic receptor subtypes within the intracranial TME, as well as on circulating immune cells. Publicly available single cell RNA sequencing (scRNAseq) datasets were interrogated and the expression of α-and β-adrenergic receptor subtypes analyzed on tumor cells, as well as on circulating and tumor-infiltrating immune populations. A dataset was generated by integrating scRNAseq data from human samples that included normal brain (n=5), newly-diagnosed GBM (n=24), recurrent GBM (n=17), low grade glioma (n=2), melanoma brain metastasis (n=8), breast cancer brain metastasis (n=3), lung cancer brain metastasis (n=3), healthy donor peripheral blood mononuclear cells (PBMC) (n=2), and PBMC from patients with GBM (n=5) **(Fig. 2A; Supplementary Fig. 1)**. Healthy brain tissue was also assessed in isolation **(Supplementary Fig. 2)**. After performing quality control, normalization, and integration, this dataset ultimately consisted of 552,442 cells. Clustering was performed and cells were visualized by Uniform Manifold Approximation and Projection (UMAP), yielding cell type-specific clusters that included tumor-, immune-, and brain-resident-cells **(Fig. 2B)**. Examination of clustering by patient **(Supplementary Fig. 1A)** and by tissue of origin **(Supplementary Fig. 1B)** demonstrated successful integration, with no evidence of batch effects. Clusters were identified and annotated according to their expression of signature population-defining genes.

**Figure 2.**
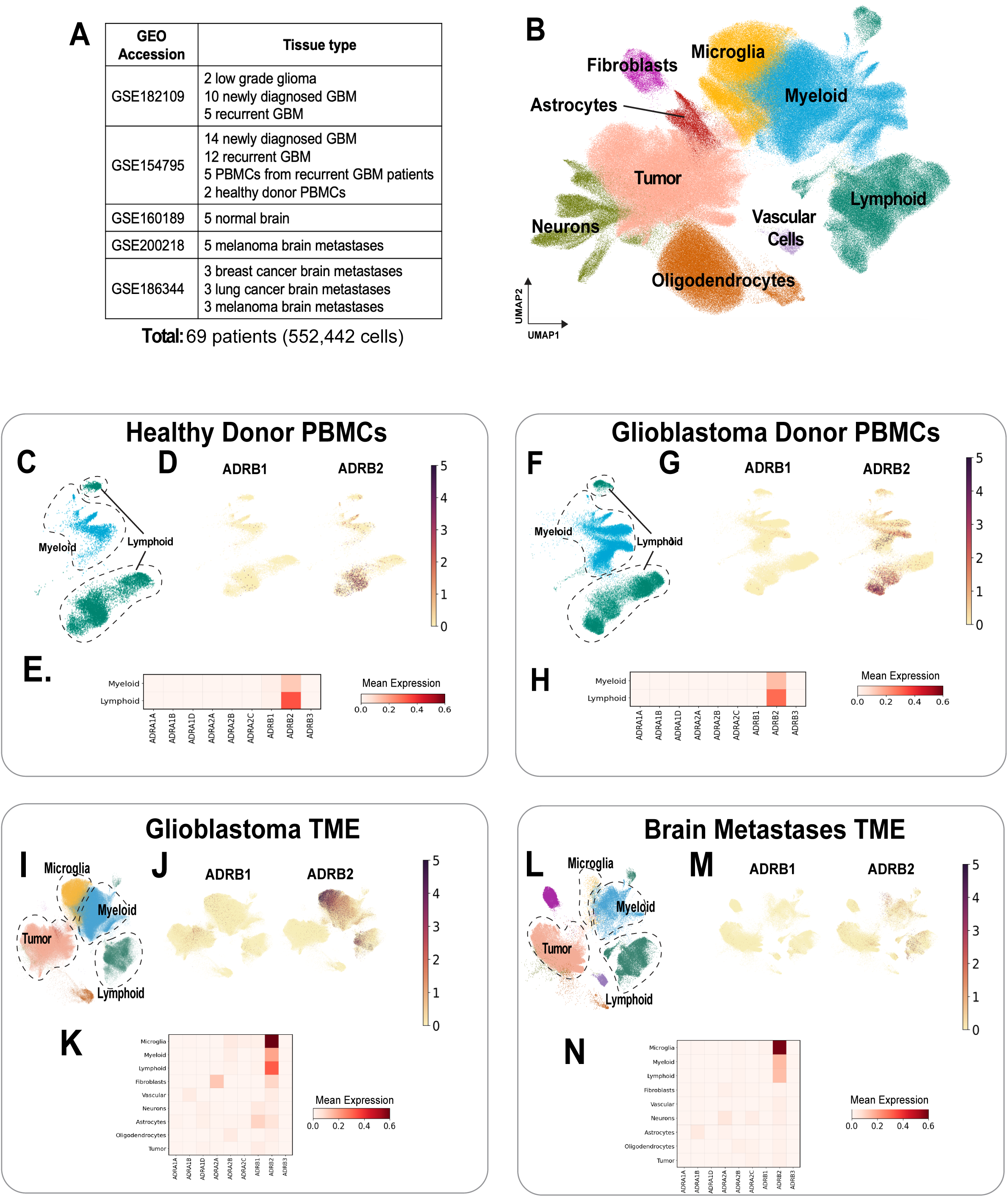
β-adrenergic receptors, not α-adrenergic receptors, are highly expressed on immune cells in the periphery and tumor microenvironment in humans. A. Integrated publicly available scRNAseq datasets from normal brain, glioblastoma, brain metastasis and PBMC samples. B. UMAP of clusters and cell types from integrated datasets. C. UMAP of clusters and cell types from healthy donor PBMCs. D. Feature plots of β1- and β2-adrenergic receptor expression in PBMCs from healthy donors. E. Heat map of β1- and β2-adrenergic receptor expression in healthy donor PBMC populations. F. UMAP of cluster and cell types from glioblastoma patient PBMCs. G. Feature plots of β1- and β2-adrenergic receptor expression in PBMCs from glioblastoma patients. H. Heat map of adrenergic receptor expression in glioblastoma PBMCs. I. UMAP of clusters and cell types glioblastoma tumor samples. J. Feature plots of β1- and β2-adrenergic receptor expression in the glioblastoma TME. K. Heat map of adrenergic receptor expression in various cell populations from the glioblastoma TME. L. UMAP of clusters and cell types from brain metastases samples. M. Feature plots of β1- and β2-adrenergic receptor expression in brain metastases TME. N. Heat map of adrenergic receptor expression in various cell types in the brain metastases TME.

Healthy donor PBMC profiles were found to consist of mainly myeloid and lymphoid clusters **(Fig. 2C),** which predominantly expressed the β2-adrenergic receptor (*ADRB2*). Low expression levels of β1-adrenergic receptors (*ADRB1*) were also seen, while expression of α- and β3-adrenergic receptors was negligible **(Fig. 2D)**. Lymphoid cells demonstrated significantly higher expression of *ADRB2* than myeloid cells **(Fig. 2E)**. PBMCs from patients with GBM yielded comparable findings **(Fig. 2F-H)**.

Repeating these analyses on tumor samples, clustering of newly diagnosed and recurrent GBM samples each revealed population subsets corresponding to tumor cells, microglia, myeloid cells, and lymphoid cells **(Fig. 2I)**. More in-depth analysis of immune compartments in PBMCs and tumor-infiltrating immune cells can be found in **Supplementary Figs. 4 and 5**. Feature plots demonstrated that the β2-adrenergic receptor was again the predominant adrenergic receptor present across all clusters, with minimal to negligible expression of the other receptor subtypes observed **(Fig. 2J and Supplementary Fig. 3)**. Likewise, lymphoid, myeloid, and microglial cells were the highest expressers of the β2-adrenergic receptor, while receptor expression was essentially absent on tumor cells **(Fig. 2K)**. Similar findings were uncovered in brain metastases **(Fig. 2L-N),** though the GBM TME appears to express higher levels of the β2-adrenergic receptor, particularly within the lymphoid compartment. β2-adrenergic receptor expression within the TME of murine CT2A gliomas was likewise confirmed by immunofluorescence **(Supplementary Fig. 6)**

Examination of feature plots of adrenergic receptor expression in healthy brain tissue demonstrated minimal expression of all receptors except for α1 (*ADRA1A* and *ADRA1B*), which was found to be concentrated in neurons and astrocytes (**Supplementary Fig. 2A**). While expression levels of β-adrenergic receptors in healthy brain were found to be quite low, comparison of the relative expression of β1- and β2-adrenergic receptors specifically indicated that astrocytes and microglia, respectively, were the cell populations with the highest expression of these two receptors (**Supplementary Fig. 2B)**.

### Systemic T cell function is impaired by **β**-adrenergic signaling but is restored by **β**-adrenergic blockade in the setting of intracranial tumors

We next sought to evaluate the contribution of elevated catecholamine levels to T cell dysfunction, which has been extensively characterized in the context of GBM. Historically, studies have reported diminished T cell reactivity, often demonstrating limited proliferative and cytokine responses ^4–6,8,9,14,16–19,24,26,27,36–42^. More current techniques assess multiple cytokines simultaneously to address T cell “polyfunctionality”,^43–54^ which has been linked to outcomes in various tumor types^55–57^.

We thus began by using a high-throughput, single-cell proteomics assay that detects cytokine secretion at the single-cell level. Splenic T cells from mice under each experimental condition were isolated and stimulated *ex vivo* with anti-CD3/anti-CD28 antibodies for 48 hours. Individual T cell production of Granzyme B, IFNγ, MIP-1α/CCL3, IP-10/CXCL10, RANTES/CCL5, IL-27, sCD137/4-1BB, IL-21, IL-17 and IL-6 was assessed and the proportion of T cells exhibiting polyfunctional cytokine production (2 or more cytokines) measured **(schematic in Fig. 3A).** In this single-cell proteomics assay, cytokine secretion was analyzed from individual cells. For validation, portions of the assay were performed twice, and representative data are shown. Flow cytometric validation of individual cytokines (Granzyme B, IFNγ) was also performed, and some of these data appear in subsequent figure panels. **Extended Figure 2A** tabulates the cytokines measured and catalogues the number of cells passing quality control (QC) for initial and repeat assays.

**Figure 3.**
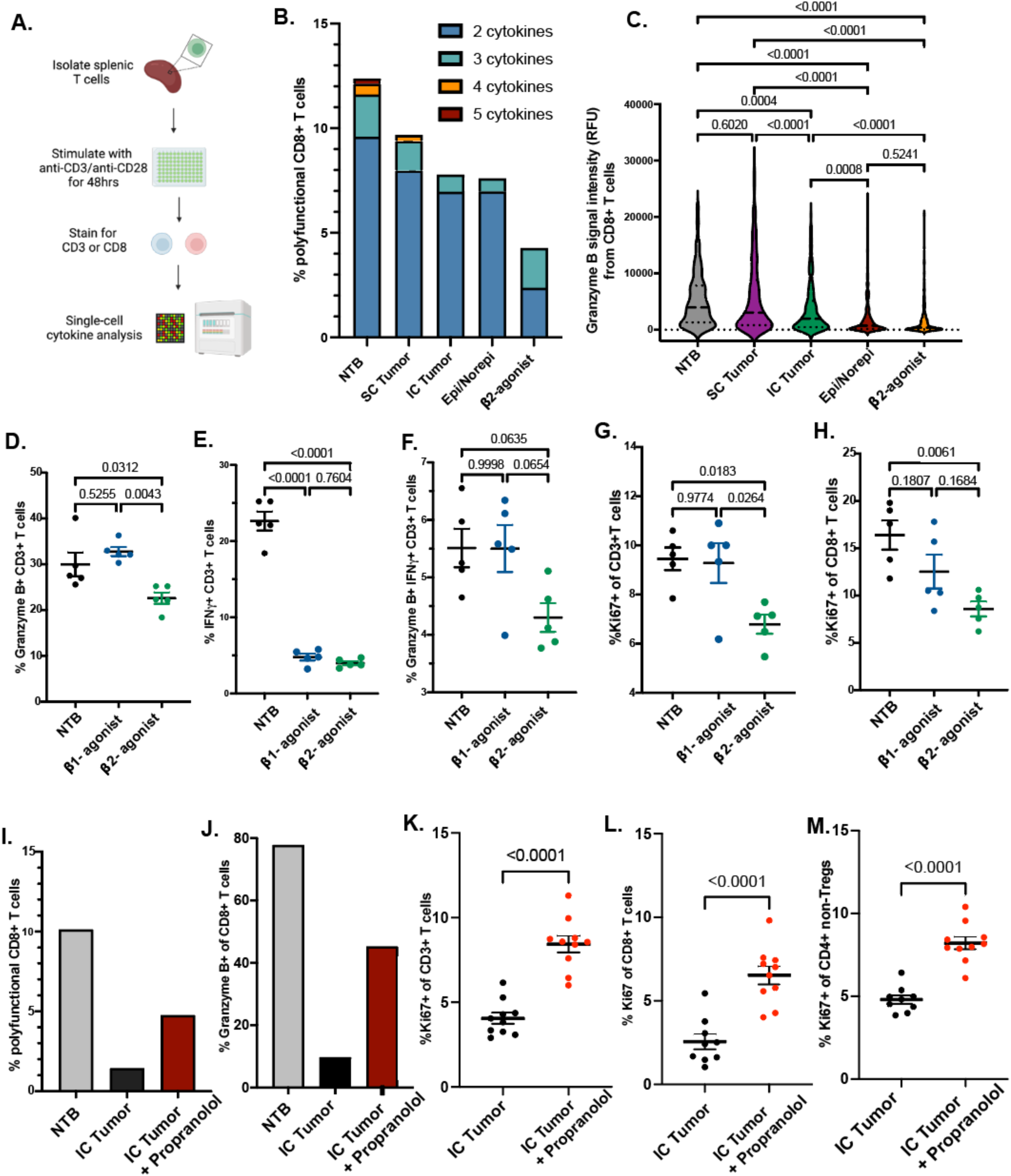
β-adrenergic signaling impairs systemic T cell function but β-adrenergic blockade restores function in the setting of intracranial tumors. A. Schematic of single-cell proteomics assay workflow (n=5 pooled mice per group for 30,000 total cells per chip). All intracranial tumor (IC tumor) mice were implanted with intracranial CT2A. All subcutaneous tumor (SC tumor) mice were implanted with subcutaneous CT2A. B. Proportion of CD8^+^ T cells from each group that are polyfunctional (individual cells that secrete two or more cytokines). C. Granzyme B signal intensity in relative fluorescence units (RFU) plotted from all available raw data for CD8^+^ T cells. Statistical significance was assessed using a one-way ANOVA with multiple comparisons. D,E. Flow cytometric analysis of non-tumor bearing (NTB) mice, or mice with subcutaneous pumps infusing dobutamine (β1-adrenergic agonist) or salbutamol (β2-adrenergic agonist). D,E. The percent of CD3^+^ T cells secreting Granzyme B (D.) or IFNγ (E.). F. Percentage of polyfunctional CD3^+^ T cells secreting both Granzyme B and IFNγ. G. Percentage of proliferating (Ki67^+^) CD3^+^ T cells. H. Percentage of proliferating (Ki67^+^) CD8^+^ T cells). Flow cytometric data reflect one experiment with n=5 mice. Statistical significance was assessed using a one-way ANOVA with multiple comparisons. I. Single-cell proteomics data demonstrating the percentage of CD8^+^ T cells that secrete two or more cytokines in mice with no tumor, intracranial CT2A or intracranial CT2A treated with the non-selective β-adrenergic antagonist propranolol. J. Single-cell proteomics data demonstrating the percentage of CD8^+^ T cells that secrete Granzyme B in mice with no tumor, intracranial CT2A or intracranial CT2A treated with propranolol. Flow cytometric analysis from mice with intracranial CT2A given no treatment or mice with intracranial CT2A treated with propranolol; percentage of proliferating K. CD3^+^ T cells L. CD8^+^ T cells or M. CD4^+^ non-Tregs. Panels K-M reflect at least two repeated experiments (n>9 mice per group). Statistical significance was assessed using unpaired t-tests.

The assay was employed to assess T cell polyfunctionality (the capacity of individual T cells to produce multiple pro-inflammatory cytokines) and to compare systemic (splenic) T cell function in the setting of either peripheral or intracranial tumor. Likewise, naïve mice were implanted with slow-release pumps and infused systemically with either a mixture of epinephrine/norepinephrine or the β2-agonist salbutamol in order to assess the sufficiency of adrenergic stimulation for eliciting compromises to T cell function (β2-agonism was chosen given the predominance of *ADRB2* expression detected within lymphoid populations above). CT2A gliomas elicited a greater decline in systemic CD8^+^ T cell function when situated intracranially, a decline that was likewise recapitulated by infusion of either catecholamines (epi/norepi) or salbutamol into naïve mice. In the context of intracranial CT2A, epi/norepi, or salbutamol, no T cells secreted more than 3 pro-inflammatory cytokines, a loss of function unique to these conditions **(Fig. 3B).** To validate at the single cytokine level and permit statistical comparisons, we examined Granzyme B production by CD8^+^ T cells. As the violin plot in **Fig. 3C** demonstrates, systemic Granzyme B production significantly declined in the setting of intracranial CT2A, epi/norepi, or salbutamol, but not subcutaneous CT2A.

A recent paper has suggested that the β1-adrenergic receptor, although not prevalent on T cells, may nonetheless be linked to T cell dysfunction^58^. Therefore, to better assess the relative contributions of both β1- and β2-adrenergic agonism to T cell dysfunction, we again implanted slow-release pumps into naïve, non-tumor-bearing (NTB) mice, this time infusing either the β1-selective agonist dobutamine or the β2-selective agonist salbutamol. Only salbutamol led to a decline in T cell Granzyme B production **(Fig. 3D)**, but both dobutamine and salbutamol severely impaired production of IFNγ **(Fig. 3E).** Salbutamol likewise elicited a reduction to the proportion of polyfunctional T cells secreting both Granzyme B and IFNγ **(Fig. 3F)**. T cell proliferation was also assessed by staining for Ki67. Only salbutamol led to a significant decline in Ki67^+^ T cells **(Fig. 3G)**, although there was a trend toward reduced proliferation in the CD8^+^ compartment of mice treated with dobutamine, as well **(Fig. 3H)**.

Given the observed impact of both β1- and β2-adrenergic agonism, we opted to evaluate the impact of treating tumor-bearing mice with the readily translatable nonselective β-adrenergic antagonist, propranolol. Treatment with propranolol partially restored polyfunctional T cell responses and Granzyme B production in mice with intracranial CT2A (**Fig. 3I,J).** The splenic CD3^+^, CD4^+^ non-Treg, and CD8^+^ T cell compartments all exhibited higher expression of the proliferation marker Ki67 (**Fig. 3 K-M)** and increases (or trending increases) in the proportion of cells with a central memory phenotype **(Extended Data Fig. 3B,C)**. Propranolol did not prevent systemic immune derangements such as lymphoid organ atrophy, T cell sequestration, or lymphopenia **(Extended Data Fig. 3)**, and there were no significant differences elicited to the overall immune composition of the various lymphoid compartments **(Supplementary Fig. 7).**

### β-adrenergic blockade proffers a pro-inflammatory intracranial TME

Having examined the impact of β-adrenergic blockade on systemic T cell function, we then turned to the local TME. Many of the proinflammatory cytokines impacted by adrenergic activity above (Granzyme B, IFNγ, MIP-1α/CCL3, sCD137/4-1BB, and RANTES/CCL5) are regulated by NF-κB. Interestingly, β2-adrenergic receptor signaling is known to suppress NF-κB activity and thus inhibit inflammatory responses in the setting of other intracranial pathologies, such as stroke^59^. We therefore sought to understand the impact of β-adrenergic blockade within the intracranial TME, as well as to investigate any influence on NF-κB-mediated immune activity. Mice with intracranial CT2A were administered either 0.5g/L propranolol or vehicle control in drinking water *ad libitum* beginning on the day of tumor implantation (Day 0). Upon clinical decline, intracranial tumors were harvested for analysis and CD45^+^ immune cells were positively selected for western blot analysis. In order to assess for differences in NF-κB at the protein level, we examined the expression of p65, the most abundant NF-κB subunit^60^. Higher levels of p65 protein expression were seen in CT2A-bearing mice treated with propranolol **(qualitative Fig. 4A; quantitative data Fig. 4B).**

**Figure 4.**
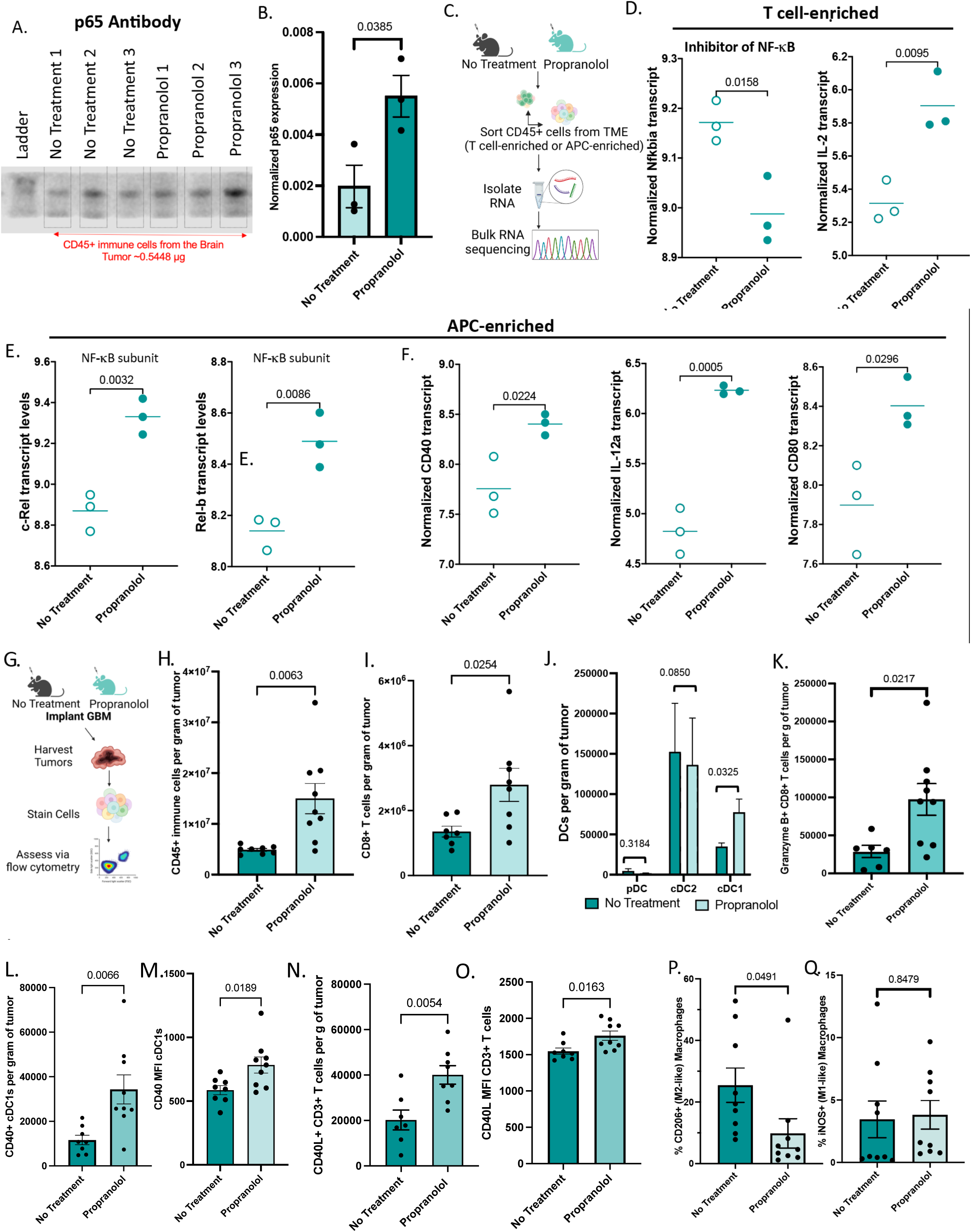
β-adrenergic receptor blockade favorably alters the local TME in GBM. A. Qualitative image of western blot denoting p65 protein expression in mice with intracranial CT2A given no treatment or propranolol. B. Quantification of p65 protein expression normalized to loading control. C. Schematic of bulk RNA-seq analysis of immune cells from the TME enriched for T cells or antigen-presenting cells (APCs). All mice were implanted with IC CT2A and tumors from 5 mice per sample were pooled (n=3 samples per group) for a total of 15 mice per group. D. RNA transcript levels of Nfkbia (IkBa) and IL-2in T cell-enriched immune cells from the TME. E. Transcript levels of NF-κB subunits c-Rel and Rel-B in APC-enriched immune cells from the TME. F. Transcript levels of genes canonically downstream of NF-kB (CD40, IL-12a and CD80) in APC-enriched immune cells from the TME. G. Schematic of flow cytometric analysis of immune phenotypes in the TME H. Number of CD45^+^ immune cells per gram of tumor (n≥8 mice per group) I. Number of CD8^+^ T cells per gram of tumor (n≥ 7 mice per group) J. Dendritic cell subsets per gram of tumor (n≥8 mice per group) K. Number of CD8^+^ T cells expressing Granzyme B per gram of tumor (n≥6 mice per group) L. Number of type 1 conventional dendritic cells (cDC1s) expressing CD40 per gram of tumor (n≥8 mice per group) M. MFI of CD40 on cDC1s (n≥8 mice per group).N. Number of CD40L^+^ CD3^+^ T cells per gram of tumor (n≥7 mice per group) O. MFI of CD40L on T cells (n≥7 mice per group) P. Percent of CD206+ immunosuppressive macrophages (n≥8 mice per group) Q. Percent of iNOS^+^ pro-inflammatory macrophages (n≥7 mice per group). Unpaired T-tests were used to calculate p values. Error bars represent standard error of the mean.

In order to further query for differences in NF-κB levels among tumor-infiltrating in immune cells, we sorted CD45^+^ cells from the TME, enriched for either T cells or APCs, and then performed bulk RNA-sequencing **(schematic in Fig. 4C).**

Indeed, T cells from tumors of mice treated with propranolol harbored significantly decreased *Nfkbia* (IkBα) transcript levels and, likewise, a striking increase in IL-2 transcript levels **(Fig. 4D).** IkBα is the “inhibitor of NF-κB” that is degraded upon activation of the canonical NF-κB signaling pathway^61^. IL-2 is a canonical downstream target of NF-κB. Together, these data support an increase in NF-kB activity in T cells within the intracranial TME. As NF-κB activation is required for T cell priming^62^, we also assessed downstream NF-κB target genes in the APC-enriched samples and observed increases in transcript levels of the NF-κB subunits c-Rel and Relb **(Fig. 4E),** as well as CD40, IL-12a and CD80 **(Fig. 4F),** all of which are expressed by dendritic cells (DCs).

Phenotypic changes to immune cells within the TME were also assessed via flow cytometry **(schematic in Fig. 4G; Gating strategy in Supplementary Fig. 8).** Treatment with propranolol increased the number of tumor-infiltrating immune cells overall **(Fig. 4H).** These increases were driven by increased numbers of CD8^+^ T cells **(Fig. 4I)** and conventional type 1 dendritic cells (cDC1s), which are critical for T cell priming and CD8^+^-mediated cytotoxicity^63,64^ **(Fig. 4J**). Accordingly, there were also more CD8^+^ T cells that produced Granzyme B in the TME of mice treated with β-adrenergic blockade (**Fig. 4K).** No differences in the proportion or number of migratory or resident cDC1s were observed (**Supplementary Fig. 10).**

Examining CD40 expression on cDC1s as a surrogate for T cell priming capacity, we found that propranolol increased both the frequency and intensity of CD40 expression on cDC1s in the TME **(Fig 4. L,M).** The same was true for CD40L on T cells, facilitating their interaction with CD40^+^ DCs (**Fig. 4N,O**). Upregulation of both CD40 and CD40L requires NF-κB activation^65,66^. The TME of mice treated with propranolol also harbored significantly fewer CD206^+^ immunosuppressive “M2-like” macrophages, with no concomitant in the percentage of iNOS^+^ inflammatory “M1-like” macrophages **(Fig. 4O-Q; Gating strategy in Supplementary Fig. 9).**

As T cell exhaustion is an established form of T cell dysfunction in the TME^67,68^ that may in turn be influenced by β1-adrenergic activity^58^, we evaluated the impact of β-adrenergic blockade on T cell differentiation states. In propranolol-treated mice, there was a slight trend among tumor-infiltrating CD8^+^ T cells towards a progenitor exhaustion phenotype (CD44^+^PD1^+^SLAMF6^+^TIM3^-^), but no difference in the prevalence of terminally exhausted cells (CD44^+^PD1^+^SLAMF6^-^TIM3^+^) **(Extended Data Fig. 4A,B)**. Likewise, while transcript levels of the classical exhaustion markers *Lag3*, *Havcr2* (Tim3) and *Pdcd1* (PD-1) trended lower in mice treated with propranolol, there was no significant difference between the groups **(Extended Data Fig. 4C)**.

### Treatment with **β**-adrenergic blockade enhances the response to immunotherapy in mice and patients with intracranial tumors

Having observed benefits to both systemic and local immune functions, we next examined whether β-adrenergic blockade might improve immunotherapeutic responses to intracranial tumors and meaningfully prolong survival. We first employed a CT2A glioma model engineered to overexpress the tumor-associated rejection antigen TRP2 (CT2A-TRP2). Mice with intracranial CT2A-TRP2 were administered either propranolol (initiated day 0); a 4-1BB agonist (on days 9, 12, 15, and 18); or the combination. The combination treatment provided a survival benefit when compared to no treatment, propranolol alone, or 4-1BB agonist alone (**Fig. 5A**). Similar patterns of survival were seen in a more stringent CT2A model that lacked a homogenous rejection antigen **(Fig. 5B).** Impact on tumor growth curves, specifically, was confirmed using a model of CT2A-luciferase and assessing tumor progression with bioluminescence imaging (BLI). Decreased rates of tumor growth were indeed observed in the combination group **(Extended Data Fig. 5).** Notably, no significant survival differences were observed between initiation of propranolol at day 0 compared to day 7, suggesting that β-adrenergic blockade can enhance immunotherapeutic responses in the context of established tumor **(Extended Data Fig. 6)**. For additional comparison, we also evaluated the impact of blocking the hypothalamic-pituitary-adrenal (HPA) axis with anti-corticotropin-releasing hormone (anti-CRH). No survival benefit was seen alone or in combination with immunotherapy (**Supplementary Fig. 11**).

**Figure 5.**
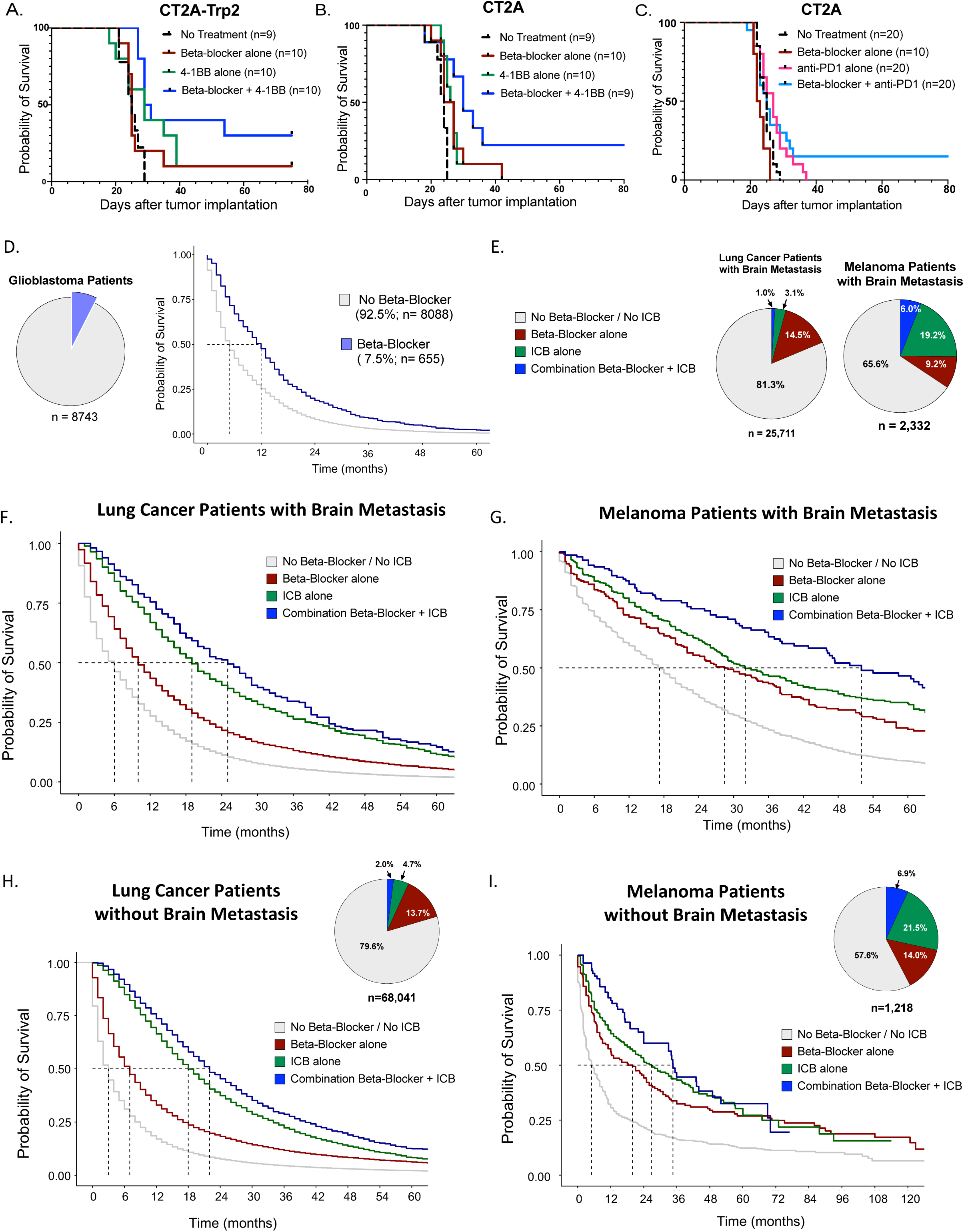
Repurposing β-adrenergic blockade enhances response to immunotherapy in primary and metastatic brain tumors. A. Kaplan-Meier survival curves of mice with IC CT2A-TRP2 receiving no treatment, β-blocker alone, 4-1BB immunotherapy alone or β-blocker in combination with 4-1BB. B. Kaplan-Meier survival curves of mice with IC CT2A (parental) receiving no treatment, β-blocker alone, 4-1BB alone or β-blocker in combination with 4-1BB. C. Kaplan-Meier survival curves of mice with IC CT2A (parental) receiving no treatment, β-blocker alone, anti-PD1 alone or β-blocker in combination with anti-PD1 D. Retrospective survival analysis of glioblastoma patients prescribed a β-blocker or not (n= 8,743; p= <0.0001). E. Patients from the SEER-Medicare database used for retrospective analysis; proportion of melanoma patients with brain metastasis or lung cancer patients with brain metastasis who received no β-blocker and no immune checkpoint blockade (ICB), β-blocker alone, immune checkpoint blockade alone or combination β-blocker and immune checkpoint blockade. F. Retrospective survival analysis of lung cancer patients with brain metastasis prescribed no immune checkpoint blockade and no β-blocker, β-blocker alone, immune checkpoint blockade alone or immune checkpoint blockade and β-blocker. G. Retrospective survival analysis of melanoma patients with brain metastasis prescribed no immune checkpoint blockade and no β-blocker, β-blocker alone, immune checkpoint blockade alone or immune checkpoint blockade and β-blocker. H. Kaplan-Meier survival curves of metastatic lung cancer patients with no brain metastasis who were prescribed no β-blocker and no immune checkpoint blockade, β-blocker alone, immune checkpoint blockade alone, or β-blocker and immune checkpoint blockade. I. Kaplan-Meier survival curves of metastatic melanoma patients with no brain metastasis who were prescribed no β-blocker and no immune checkpoint blockade, β-blocker alone, immune checkpoint blockade alone, or β-blocker and immune checkpoint blockade. Survival was assessed via log-rank (Mantel-Cox) tests.

The studies above were conducted with 4-1BB agonism, given the documented capacity to elicit a modest anti-tumor immune response against glioma^69^. We now inquired, however, whether β-adrenergic blockade might also be capable of licensing a response to anti-PD1 therapy, to which human GBM and murine CT2A have both been notoriously resistant^1,20,26,69,70^. The combination of propranolol and anti-PD1 indeed resulted in long-term survival, while neither therapy alone produced such therapeutic impact **(Fig. 5C)**.

To assess the significance of these findings to patients, we utilized the Surveillance, Epidemiology and End Results (SEER)-Medicare database to first retrospectively assess the impact of β-blockade therapy (prescribed for any indication) on outcome in patients with GBM (treatment codes outlined in **Supplementary Table 1**). As no immunotherapies are currently FDA-approved in GBM, they are not insurance-reimbursed and thus no data on the combination of β-blockade and immunotherapy are currently available through the SEER-Medicare database. We were, however, able to assess the impact of β-blockade monotherapy in patients with GBM. For the purposes of this retrospective study, a total of 8,743 patients met criteria, including the intrinsic criterion age > 65 (patients <65 are not recorded in Medicare data, unless they possess specific co-morbidities). Medicare patients enrolled due to end-stage renal disease or disability were excluded. Patients enrolled in a health maintenance organization (HMO) 6 months prior to and 12 months after primary diagnosis were also excluded (see demographics in **Table 1.1)**. Of these 8,743 patients, 655 had been prescribed β-adrenergic blockade at the time of their cancer diagnosis. Median overall survival (OS) of GBM patients receiving β-adrenergic blockade (12 months) was more than 200% that of those not receiving β-adrenergic blockade (5 months) (HR 0.63; 0.57, 0.69 95% CI; p<0.0001) **(Fig. 5D**; **Table 1.2, 1.3).**

**Table 1.** Demographics and Survival Data from Glioblastoma Patients in SEER-Medicare 2008-2017.

**1.1. Overall**
| Characteristic | N = 8,743<sup>1</sup> |
| --- | --- |
| **Patient Age** | 74 (69, 80) |
| **Sex** |  |
| M | 4,699 (54%) |
| F | 4,044 (46%) |
| **Race/ethnicity** |  |
| NH White | 7,572 (87%) |
| NH Black | 307 (3.5%) |
| NH Other | 315 (3.6%) |
| Hispanic | 549 (6.3%) |
| **Charlson score** |  |
| 0 | 4,654 (53%) |
| 1 | 1,854 (21%) |
| 2 | 1,049 (12%) |
| 3+ | 1,186 (14%) |
| **Received radiation** |  |
| None | 3,347 (38%) |
| Yes | 5,396 (62%) |
| **Received chemotherapy** |  |
| None/Unknown | 2,802 (44%) |
| Yes | 3,591 (56%) |
| Unknown | 2,350 |
| **Received surgery of primary** |  |
| None | 2,411 (28%) |
| Yes | 6,332 (72%) |
| **Any betablocker** |  |
| No betablocker | 8,088 (93%) |
| Betablocker | 655 (7.5%) |

<sup>1</sup>Median (IQR); n (%)

1.2. Median Survival Months
| Characteristic | Median Survival |
| --- | --- |
| **Any betablocker** |  |
| No betablocker | 5.0 (5.0, 5.0) |
| Betablocker | 12 (11, 13) |

1.3. Adjusted Survival Model
| Characteristic | HR<sup>1</sup> | 95% CI<sup>1</sup> | p-value |
| --- | --- | --- | --- |
| **Patient Age** | 1.02 | 1.02, 1.03 | **<0.001** |
| **Sex** |  |  |  |
| M | — | — |  |
| F | 1.00 | 0.95, 1.05 | 0.920 |
| **Race/ethnicity** |  |  |  |
| NH White | — | — |  |
| NH Black | 0.77 | 0.67, 0.88 | **<0.001** |
| NH Other | 0.74 | 0.66, 0.84 | **<0.001** |
| Hispanic | 0.95 | 0.86, 1.05 | 0.282 |
| **Charlson score** |  |  |  |
| 0 | — | — |  |
| 1 | 1.14 | 1.07, 1.22 | **<0.001** |
| 2 | 1.28 | 1.18, 1.39 | **<0.001** |
| 3+ | 1.41 | 1.30, 1.52 | **<0.001** |
| **Received radiation** |  |  |  |
| None | — | — |  |
| Yes | 0.56 | 0.52, 0.60 | **<0.001** |
| **Received chemotherapy** |  |  |  |
| None/Unknown | — | — |  |
| Yes | 0.56 | 0.52, 0.60 | **<0.001** |
| **Received surgery of primary** |  |  |  |
| None | — | — |  |
| Yes | 0.62 | 0.59, 0.66 | **<0.001** |
| **Any betablocker** |  |  |  |
| No betablocker | — | — |  |
| Betablocker | 0.63 | 0.57, 0.69 | **<0.001** |

<sup>1</sup>HR = Hazard Ratio, CI = Confidence Interval

In contrast to GBM, checkpoint blockade is FDA-approved in patients with brain metastases arising from melanoma and from certain lung cancers, permitting us to identify patients with these conditions who may have been treated with checkpoint blockade and β-adrenergic blockade either concomitantly or individually. We therefore returned to the SEER-Medicare database to retrospectively determine survival outcomes in patients with melanoma or lung cancer brain metastases who had received checkpoint blockade alone for their cancer versus those who had received checkpoint blockade concomitantly with β-adrenergic blockade (the latter again for any indication). Survival was also assessed in patients who received β-adrenergic blockade alone (for any indication), as well as in those who received neither β-adrenergic blockade nor checkpoint blockade.

A total of 25,711 lung cancer patients and 2,332 melanoma patients, all diagnosed with concomitant brain metastases, were assessed **(Fig. 5E).** Beginning with lung cancer, we evaluated outcomes in patients with brain metastasis who met the inclusion criteria described previously. Patients receiving no β-blocker and no immune checkpoint blockade (n=20,905) had a median OS of 6 months, while those receiving a β-blocker alone (n=3,733) had a longer median OS of 10 months (p<0.001). Patients receiving checkpoint blockade alone (n=806) exhibited a median OS of 19 months, while those prescribed both a β-blocker and checkpoint blockade (n=267) had a median OS of 25 months, which represented a > 30% increase (25 months vs. 19 months, p=0.013). Patients receiving both therapies also survived >150% longer than those that received β-blocker alone (25 months vs. 10 months, p<0.001) and >300% longer than those patients that received neither β-blocker nor checkpoint blockade (25 months vs. 6 months, p<0.001) **(Fig. 5F**; **Table 2). Table 2.4** demonstrates the hazard ratios (HR) for comparisons amongst groups.

**Table 2.** Demographics and Survival Data of Metastatic Lung Cancer Patients with Brain Metastasis in SEER-Medicare 2008-2017.

**2.1 Overall**
| Characteristic | N = 25,711<sup>1</sup> |
| --- | --- |
| **Patient Age** | 73 (68, 78) |
| **Sex** |  |
| M | 12,485 (49%) |
| F | 13,226 (51%) |
| **Race/ethnicity** |  |
| NH White | 20,813 (81%) |
| Hispanic | 1,224 (4.8%) |
| NH Black | 1,993 (7.8%) |
| NH Other | 1,681 (6.5%) |
| **Charlson score** |  |
| 0 | 11,288 (44%) |
| 1 | 7,148 (28%) |
| 2 | 3,542 (14%) |
| 3+ | 3,733 (15%) |
| **Received radiation** |  |
| None | 9,359 (36%) |
| Yes | 16,352 (64%) |
| **Received chemotherapy** |  |
| None | 15,137 (59%) |
| Yes | 10,574 (41%) |
| **Received surgery of primary** |  |
| None | 23,736 (92%) |
| Yes | 1,975 (7.7%) |
| **LC subtype** |  |
| NSCLC | 20,326 (79%) |
| SCLC | 5,385 (21%) |
| **Betablocker and checkpoint inhibitor use** |  |
| No betablocker/No ICB | 20,905 (81%) |
| Betablocker only | 3,733 (15%) |
| ICB only | 806 (3.1%) |
| Betablocker and ICB | 267 (1.0%) |

<sup>1</sup>Median (IQR); n (%)

2.2 Pairwise Log-rank P-values
| Treatment | No betablocker/No ICB | Betablocker only | ICB only |
| --- | --- | --- | --- |
| Betablocker only | <0.001 |  |  |
| ICB only | <0.001 | <0.001 |  |
| Betablocker and ICB | <0.001 | <0.001 | 0.013 |

2.3 Median Survival Months
| Characteristic | Median Survival |
| --- | --- |
| **Betablocker and checkpoint inhibitor use** |  |
| No betablocker/No ICB | 6.0 (5.0, 6.0) |
| Betablocker only | 10 (10, 11) |
| ICB only | 19 (18, 21) |
| Betablocker and ICB | 25 (22, 29) |

2.4 Adjusted Survival Model
| Characteristic | HR<sup>1</sup> | 95% CI<sup>1</sup> | p-value |
| --- | --- | --- | --- |
| **Patient Age** | 1.02 | 1.01, 1.02 | **<0.001** |
| **Sex** |  |  |  |
| M | — | — |  |
| F | 0.83 | 0.81, 0.85 | **<0.001** |
| **Race/ethnicity** |  |  |  |
| NH White | — | — |  |
| Hispanic | 0.97 | 0.92, 1.03 | 0.351 |
| NH Black | 1.02 | 0.97, 1.07 | 0.481 |
| NH Other | 0.79 | 0.75, 0.84 | **<0.001** |
| **Charlson score** |  |  |  |
| 0 | — | — |  |
| 1 | 1.04 | 1.01, 1.07 | **0.021** |
| 2 | 1.07 | 1.03, 1.11 | **<0.001** |
| 3+ | 1.17 | 1.13, 1.22 | **<0.001** |
| **Received radiation** |  |  |  |
| None | — | — |  |
| Yes | 0.76 | 0.74, 0.78 | **<0.001** |
| **Received chemotherapy** |  |  |  |
| None | — | — |  |
| Yes | 0.65 | 0.63, 0.67 | **<0.001** |
| **Received surgery of primary** |  |  |  |
| None | — | — |  |
| Yes | 0.32 | 0.31, 0.34 | **<0.001** |
| **LC subtype** |  |  |  |
| NSCLC | — | — |  |
| SCLC | 0.97 | 0.94, 1.00 | 0.075 |
| **Betablocker and checkpoint inhibitor use** |  |  |  |
| No betablocker/No ICB | — | — |  |
| Betablocker only | 0.65 | 0.63, 0.67 | **<0.001** |
| ICB only | 0.46 | 0.43, 0.50 | **<0.001** |
| Betablocker and ICB | 0.37 | 0.32, 0.43 | **<0.001** |

<sup>1</sup>HR = Hazard Ratio, CI = Confidence Interval

We next evaluated outcomes in melanoma patients with brain metastasis as above. Of the 2,332 patients included, 1,531 patients received neither β-blocker nor checkpoint blockade and exhibited a median OS of 17 months, while patients that received β-blocker only (n=215) exhibited a median OS of 28 months (28 months vs. 17 months, p<0.001). Patients who received checkpoint blockade alone exhibited a median OS of 32 months, while the 139 patients who were prescribed both a β-blocker and checkpoint blockade exhibited a median OS of 52 months (52 months vs. 32 months, p=0.006). This represents a >60% increase in median OS in patients who received both β-blockade and checkpoint blockade compared to checkpoint blockade alone and a >200% increase compared to those patients that received neither β-blocker nor checkpoint blockade (52 months vs. 17 months, p<0.001) **(Fig. 5G**; **Table 3)**. **Table 3.4** demonstrates the hazard ratios (HR) for comparisons amongst groups.

**Table 3.** Demographics and Survival Data from Metastatic Melanoma Patients with Brain Metastasis in SEER-Medicare 2008-2017.

**3.1. Overall**
| Characteristic | N = 2,332<sup>1</sup> |
| --- | --- |
| **Patient Age** | 74 (69, 81) |
| **Sex** |  |
| Male 1,606 (69%) Female 726 (31%) |  |
| **Race/ethnicity** |  |
| NH White | 2,216 (95%) |
| Hispanic 66 (2.8%) NH Black 22 (0.9%) |  |
| NH Other | 28 (1.2%) |
| **Charlson score** |  |
| 0 1,327 (57%) |  |
| 1 335 (14%) 2 362 (16%) |  |
| 3+ | 308 (13%) |
| **Received radiation** |  |
| None 1,473 (63%) Yes 859 (37%) |  |
| **Received chemotherapy** |  |
| None | 1,323 (57%) |
| Yes | 1,009 (43%) |
| **Received surgery of primary** |  |
| None | 819 (35%) |
| Yes | 1,513 (65%) |
| **Betablocker and checkpoint inhibitor use** |  |
| No betablocker/No ICB | 1,531 (66%) |
| Betablocker only | 215 (9.2%) |
| Characteristic | N = 2,332<sup>1</sup> |
| ICB only | 447 (19%) |
| Betablocker and ICB | 139 (6.0%) |

<sup>1</sup>Median (IQR); n (%)

3.2. Pairwise Log-rank P-values
| Treatment | No betablocker/No ICB | Betablocker only | ICB only |
| --- | --- | --- | --- |
| Betablocker only | <0.001 |  |  |
| ICB only | <0.001 | 0.066 |  |
| Betablocker and ICB | <0.001 | <0.001 | 0.006 |

3.3. Median Survival Months
| Characteristic | Median Survival |
| --- | --- |
| **Betablocker and checkpoint inhibitor use** |  |
| No betablocker/No ICB | 17 (16, 19) |
| Betablocker only | 28 (23, 37) |
| ICB only | 32 (29, 39) |
| Betablocker and ICB | 52 (46, 70) |

3.4. Adjusted Survival Model
| Characteristic | HR<sup>1</sup> | 95% CI<sup>1</sup> | p-value |
| --- | --- | --- | --- |
| **Patient Age** | 1.02 | 1.01, 1.02 | **<0.001** |
| **Sex** |  |  |  |
| Male | — | — |  |
| Female | 0.90 | 0.81, 0.99 | **0.029** |
| **Race/ethnicity** |  |  |  |
| NH White | — | — |  |
| Hispanic | 1.06 | 0.81, 1.39 | 0.675 |
| NH Black | 0.87 | 0.56, 1.36 | 0.547 |
| NH Other | 0.81 | 0.54, 1.22 | 0.323 |
| **Charlson score** |  |  |  |
| 0 | — | — |  |
| 1 | 1.08 | 0.95, 1.24 | 0.236 |
| 2 | 1.14 | 1.00, 1.30 | **0.043** |
| 3+ | 1.33 | 1.16, 1.53 | **<0.001** |
| **Received radiation** |  |  |  |
| None | — | — |  |
| Yes | 1.44 | 1.30, 1.58 | **<0.001** |
| **Received chemotherapy** |  |  |  |
| None | — | — |  |
| Yes | 0.72 | 0.65, 0.81 | **<0.001** |
| **Received surgery of primary** |  |  |  |
| None | — | — |  |
| Yes | 0.47 | 0.43, 0.52 | **<0.001** |
| **Betablocker and checkpoint inhibitor use** |  |  |  |
| No betablocker/No ICB | — | — |  |
| Betablocker only | 0.54 | 0.46, 0.63 | **<0.001** |
| ICB only | 0.56 | 0.49, 0.65 | **<0.001** |
| Betablocker and ICB | 0.40 | 0.31, 0.50 | **<0.001** |

<sup>1</sup> HR = Hazard Ratio, CI = Confidence Interval

β-adrenergic receptor expression has been seen on melanoma^71–74^ and other cancers,^74–77^ with a direct role in tumor progression reported. In melanoma, in particular, studies suggest that combination β-adrenergic and checkpoint blockade may promote extended recurrence-free survival in the absence of intracranial disease.^72,78^ Given our findings of increased adrenergic activity in the setting of intracranial tumors, specifically, we looked to establish the relative potency of β-adrenergic blockade for licensing immunotherapeutic responses when intracranial disease was present versus not. Finally, then, we examined the effects of concomitant β-blocker and checkpoint blockade in metastatic lung cancer and metastatic melanoma patients who had no brain metastases (metastatic extracranial disease only, at diagnosis). Median OS among lung cancer patients who received combination β-blocker and checkpoint blockade therapy in the setting of extracranial disease only was 22 months vs. 18 months among those who received checkpoint blockade only (p<0.001). This represented a modest ∼20% increase (respective increase in the setting of intracranial disease was 32%) **(Fig. 5H and Table 4.3)**. Median survival in patients with exclusively extracranial melanoma in the combination group was 35 months compared to 27 months in those who only received checkpoint blockade (∼30% increase), although this difference was not statistically significant **(Fig. 5I**; **Table 5.3)** (respective increase in the setting of intracranial disease was 63%, p=0.006). In all cohorts, the cause of death for the majority of patients was their cancer (**Supplementary Fig. 12).** Thus, while β-blockade also impacts outcomes in the setting of extracranial disease, the benefits are especially pronounced in patients harboring intracranial disease burdens.

**Table 4.**
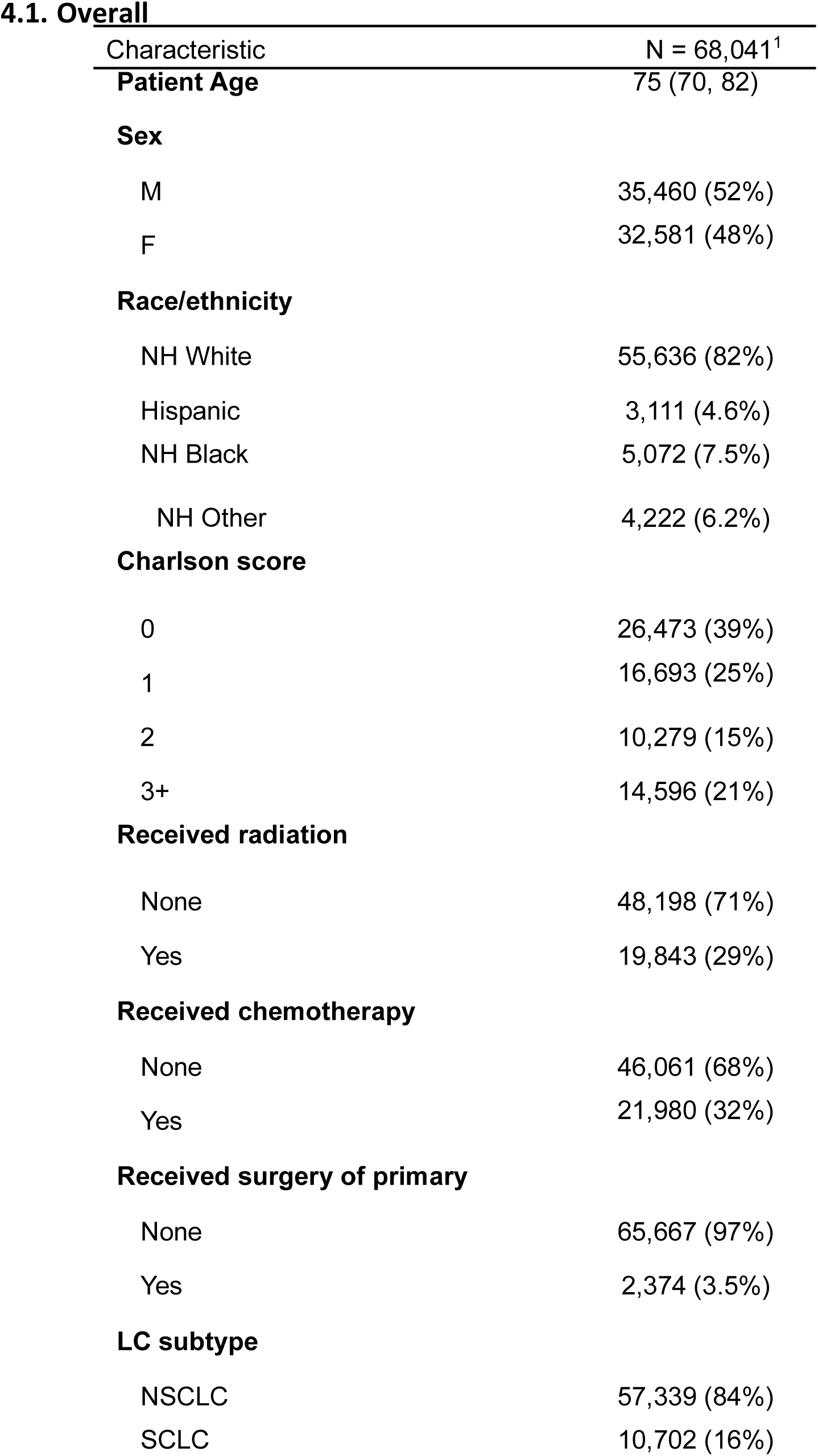
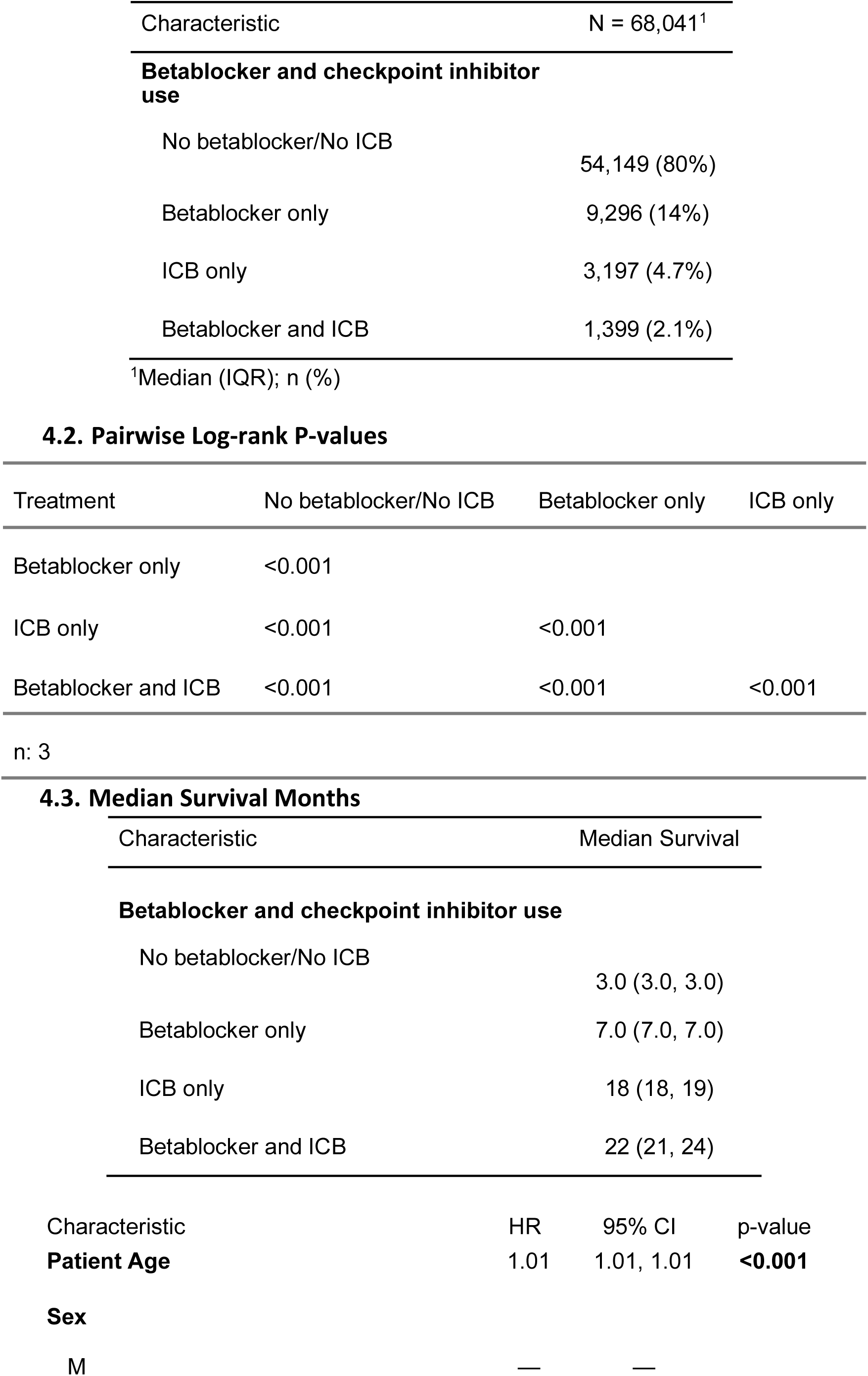
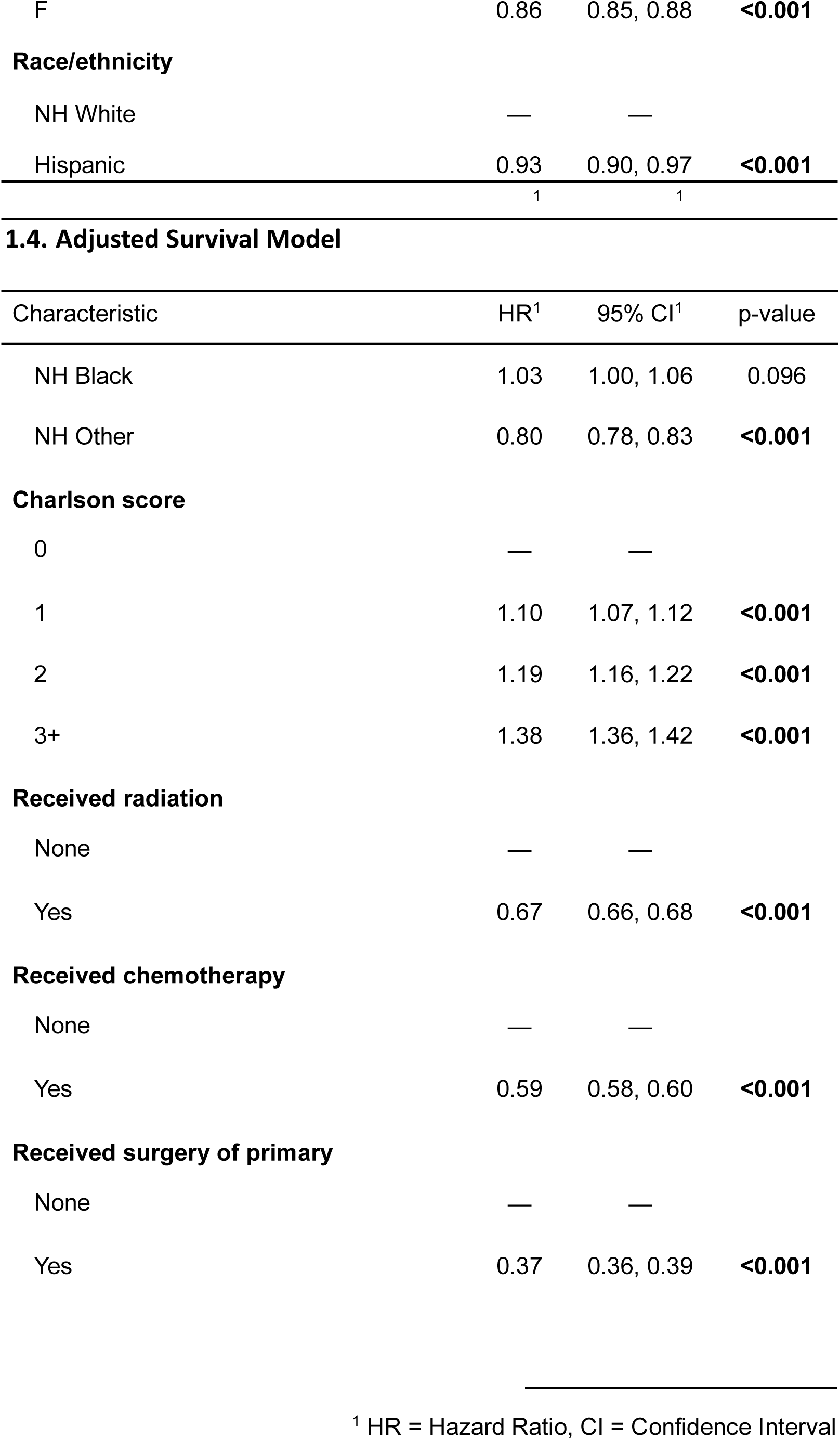
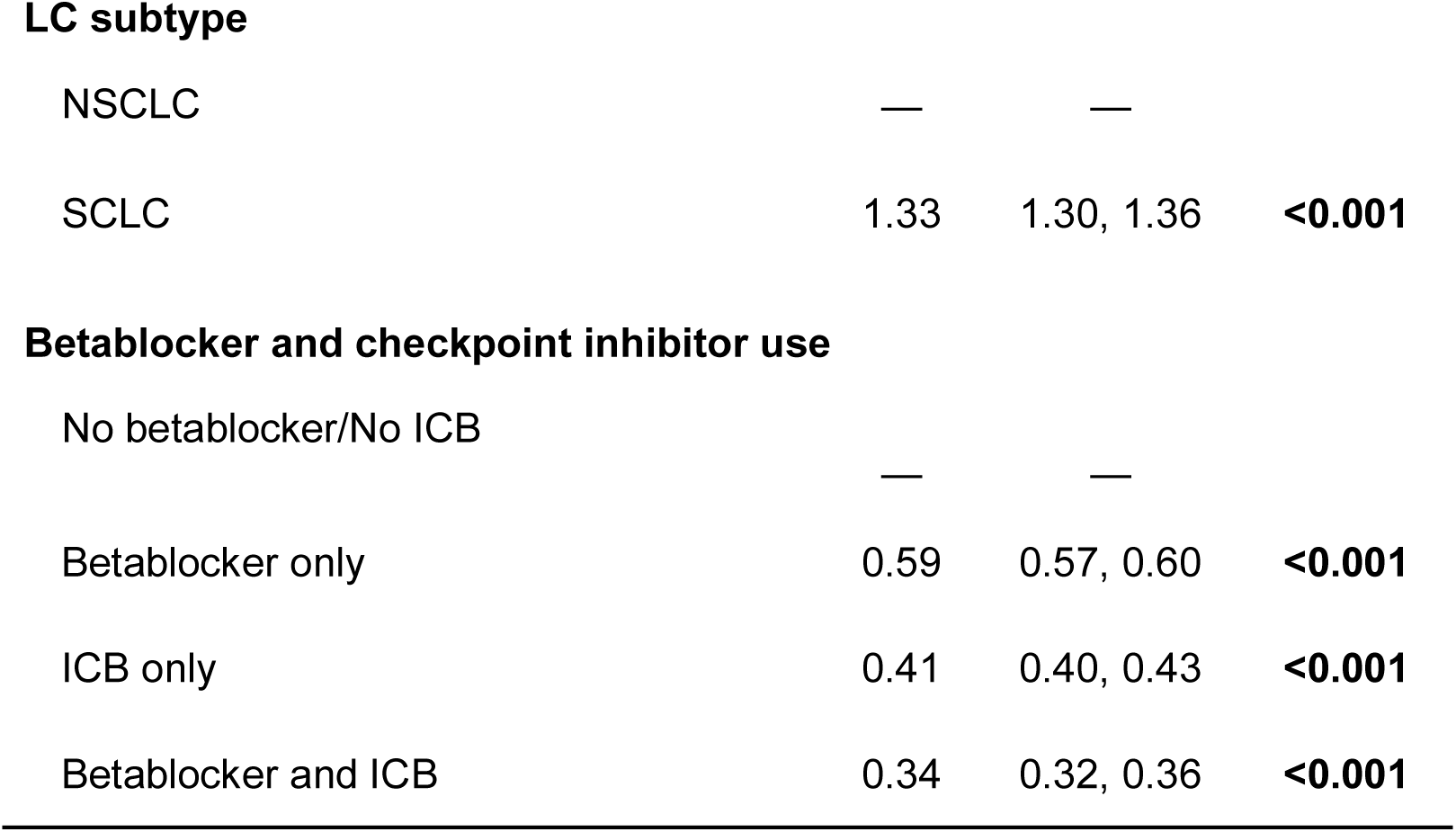
Demographics and Survival Data from Metastatic Lung Cancer Patients with no Brain Metastasis in SEER-Medicare 2008-2017.

**4.1. Overall**
| Characteristic | N = 68,041<sup>1</sup> |
| --- | --- |
| **Patient Age** | 75 (70, 82) |
| **Sex** |  |
| M | 35,460 (52%) |
| F | 32,581 (48%) |
| **Race/ethnicity** |  |
| NH White | 55,636 (82%) |
| Hispanic | 3,111 (4.6%) |
| NH Black | 5,072 (7.5%) |
| NH Other | 4,222 (6.2%) |
| **Charlson score** |  |
| 0 | 26,473 (39%) |
| 1 | 16,693 (25%) |
| 2 | 10,279 (15%) |
| 3+ | 14,596 (21%) |
| **Received radiation** |  |
| None | 48,198 (71%) |
| Yes | 19,843 (29%) |
| **Received chemotherapy** |  |
| None | 46,061 (68%) |
| Yes | 21,980 (32%) |
| **Received surgery of primary** |  |
| None | 65,667 (97%) |
| Yes | 2,374 (3.5%) |
| **LC subtype** |  |
| NSCLC | 57,339 (84%) |
| SCLC | 10,702 (16%) |
| **Betablocker and checkpoint inhibitor use** |  |
| No betablocker/No ICB | 54,149 (80%) |
| Betablocker only | 9,296 (14%) |
| ICB only | 3,197 (4.7%) |
| Betablocker and ICB | 1,399 (2.1%) |

<sup>1</sup>Median (IQR); n (%)

4.2. Pairwise Log-rank P-values
| Treatment | No betablocker/No ICB | Betablocker only | ICB only |
| --- | --- | --- | --- |
| Betablocker only | <0.001 |  |  |
| ICB only | <0.001 | <0.001 |  |
| Betablocker and ICB | <0.001 | <0.001 | <0.001 |

4.3. Median Survival Months
| Characteristic | Median Survival |
| --- | --- |
| **Betablocker and checkpoint inhibitor use** |  |
| No betablocker/No ICB | 3.0 (3.0, 3.0) |
| Betablocker only | 7.0 (7.0, 7.0) |
| ICB only | 18 (18, 19) |
| Betablocker and ICB | 22 (21, 24) |

| Characteristic | HR | 95% CI | p-value |
| --- | --- | --- | --- |
| **Patient Age** | 1.01 | 1.01, 1.01 | **<0.001** |
| **Sex** |  |  |  |
| M | — | — |  |
| F | 0.86 | 0.85, 0.88 | **<0.001** |
| **Race/ethnicity** |  |  |  |
| NH White | — | — |  |
| Hispanic | 0.93 | 0.90, 0.97 | **<0.001** |
|  | 1 | 1 |  |

1.4. Adjusted Survival Model
| Characteristic | HR<sup>1</sup> | 95% CI<sup>1</sup> | p-value |
| --- | --- | --- | --- |
| NH Black | 1.03 | 1.00, 1.06 | 0.096 |
| NH Other | 0.80 | 0.78, 0.83 | **<0.001** |
| **Charlson score** |  |  |  |
| 0 | — | — |  |
| 1 | 1.10 | 1.07, 1.12 | **<0.001** |
| 2 | 1.19 | 1.16, 1.22 | **<0.001** |
| 3+ | 1.38 | 1.36, 1.42 | **<0.001** |
| **Received radiation** |  |  |  |
| None | — | — |  |
| Yes | 0.67 | 0.66, 0.68 | **<0.001** |
| **Received chemotherapy** |  |  |  |
| None | — | — |  |
| Yes | 0.59 | 0.58, 0.60 | **<0.001** |
| **Received surgery of primary** |  |  |  |
| None | — | — |  |
| Yes | 0.37 | 0.36, 0.39 | **<0.001** |

<sup>1</sup> HR = Hazard Ratio, CI = Confidence Interval

**LC subtype**
|  |  |  |  |
| --- | --- | --- | --- |
| NSCLC | — | — |  |
| SCLC | 1.33 | 1.30, 1.36 | **<0.001** |

**Betablocker and checkpoint inhibitor use**
|  |  |  |  |
| --- | --- | --- | --- |
| No betablocker/No ICB | — | — |  |
| Betablocker only | 0.59 | 0.57, 0.60 | **<0.001** |
| ICB only | 0.41 | 0.40, 0.43 | **<0.001** |
| Betablocker and ICB | 0.34 | 0.32, 0.36 | **<0.001** |

**Table 5.** Demographics and Survival Data from Metastatic Melanoma Patients with no Brain Metastasis in SEER-Medicare 2008-2017.

**5.1. Overall**
| Characteristic |  | N = 1,218<sup>1</sup> |
| --- | --- | --- |
| **Patient Age** |  | 77 (70, 84) |
| **Sex** |  |  |
| Male |  | 797 (65%) |
| Female |  | 421 (35%) |
| **Race/ethnicity** |  |  |
| NH White |  | 1,139 (94%) |
| Hispanic | 42 (3.4%) | NH Black 18 (1.5%) |
| NH Other |  | 19 (1.6%) |
| **Charlson score** |  |  |
| 0 | 660 (54%) | 1 174 (14%) 2 171 (14%) |
| 3+ |  | 213 (17%) |
| **Received radiation** |  |  |
| None | 1,022 (84%) | Yes 196 (16%) |
| **Received chemotherapy** |  |  |
| None |  | 759 (62%) |
| Yes |  | 459 (38%) |
| **Received surgery of primary** |  |  |
| None |  | 751 (62%) |
| Yes |  | 467 (38%) |
| **Betablocker and checkpoint inhibitor use** |  |  |
| No betablocker/No ICB |  | 702 (58%) |
| Betablocker only |  | 170 (14%) |
| Characteristic |  | N = 1,218<sup>1</sup> |
| ICB only |  | 262 (22%) |
| Betablocker and ICB |  | 84 (6.9%) |

<sup>1</sup>Median (IQR); n (%)

5.2. Pairwise Log-rank P-values
| Treatment | No betablocker/No ICB | Betablocker only | ICB only |
| --- | --- | --- | --- |
| Betablocker only | <0.001 |  |  |
| ICB only | <0.001 | 0.165 |  |
| Betablocker and ICB | <0.001 | 0.019 | 0.158 |

5.3. Median Survival Months
| Characteristic | Median Survival |
| --- | --- |
| **Betablocker and checkpoint inhibitor use** |  |
| No betablocker/No ICB | 5.0 (4.0, 6.0) |
| Betablocker only | 20 (12, 26) |
| ICB only | 27 (21, 38) |
| Betablocker and ICB | 35 (24, 52) |

5.4. Adjusted Survival Model
| Characteristic | HR<sup>1</sup> | 95% CI<sup>1</sup> | p-value |
| --- | --- | --- | --- |
| **Patient Age** | 1.02 | 1.01, 1.03 | **<0.001** |
| **Sex** |  |  |  |
| Male | — | — |  |
| Female | 1.01 | 0.88, 1.16 | 0.878 |
| **Race/ethnicity** |  |  |  |
| NH White | — | — |  |
| Hispanic | 0.84 | 0.58, 1.21 | 0.358 |
| NH Black | 1.30 | 0.79, 2.15 | 0.299 |
| NH Other | 1.34 | 0.83, 2.14 | 0.229 |
| **Charlson score** |  |  |  |
| 0 | — | — |  |
| 1 | 1.20 | 0.98, 1.47 | 0.072 |
| 2 | 1.29 | 1.06, 1.58 | **0.013** |
| 3+ | 1.56 | 1.30, 1.86 | **<0.001** |
| **Received radiation** |  |  |  |
| None | — | — |  |
| Yes | 0.94 | 0.79, 1.12 | 0.470 |
| **Received chemotherapy** |  |  |  |
| None | — | — |  |
| Yes | 0.61 | 0.50, 0.75 | **<0.001** |
| **Received surgery of primary** |  |  |  |
| None | — | — |  |
| Yes | 0.59 | 0.52, 0.68 | **<0.001** |
| **Betablocker and checkpoint inhibitor use** |  |  |  |
| No betablocker/No ICB | — | — |  |
| Betablocker only | 0.50 | 0.41, 0.61 | **<0.001** |
| ICB only | 0.68 | 0.54, 0.86 | **0.002** |
| Betablocker and ICB | 0.53 | 0.37, 0.74 | **<0.001** |

<sup>1</sup> HR = Hazard Ratio, CI = Confidence Interval

## DISCUSSION

The intracranial compartment proffers a unique set of challenges that can serve to limit immunotherapeutic efficacy. Treatment failures have been especially prominent in the context of GBM. Historical impediments include the blood-brain barrier (BBB)^26,79–81^; challenges to T cell priming and function^82–85^ and a heavily immunosuppressive TME^22,26,37,82–88^. Furthermore, despite rarely leaving its intracranial confines, GBM is known to suppress systemic immune function. Much of this depressed systemic immune function is not unique to GBM, but instead accompanies pathologies of the intracranial compartment more broadly^20,89^. The mechanistic underpinnings of such systemic restrictions to immune function and their implications for immunotherapeutic success remain largely unexplored^90^. Herein we reveal that tumors of the intracranial compartment, uniquely, elicit steep rises in systemic catecholamine levels that are sufficient to restrict both local and systemic immune function. Conversely, treatment with nonselective β-adrenergic blockade increases NF-κB activity in immune cells, restores T cell polyfunctionality, modifies the TME, and licenses immune-based therapies in murine models of GBM.

The immunosuppressive effects of sympathetic activity are well-documented^91–94^. Likewise, elevated systemic catecholamine levels have been described in the setting of other intracranial pathologies, such as stroke, where they are an identified component of “stroke-induced immunodepression syndrome” (SIDS)^95^. Studies suggest that SIDS may be attributable to inhibition of NF-κB activity following β-adrenergic signaling^96^. Our data suggest that a similar immunosuppressive axis may be active in the context of intracranial tumors as a result of chronic sympathetic hyperactivity. Notably, we observe dramatic catecholamine rises in the spleen and in circulation, but not in the TME. These data suggest that the chronic sympathetic hyperactivity arises from triggering of a systemic sympathetic response, and not simply local production of catecholamines within the tumor. Additionally, there was a relative lack of adrenergic receptor expression found on human GBM cells, with higher expression found on both systemic and tumor-infiltrating immune cells. Consequently, β-adrenergic blockade likely acts predominantly on the systemic immune compartment. Further investigation into the immediate upstream determinants of sympathetic hyperactivity in the setting of intracranial tumors is warranted.

Our data demonstrate that both β1- and β2-adrenergic signaling likely play a role in contributing to catecholamine-induced immune dysfunction, although β2-adrenergic receptors are the more prevalent receptor subtype on systemic immune cells and within the TME. A recent publication has advanced a role for the β1-adrenergic receptor in promoting T cell exhaustion^58^, yet also found that the greatest improvements to immune responses in extracranial tumors were seen with nonselective (rather than β1-specific) adrenergic blockade^58^. Our results are in accordance with these findings and, together, serve to highlight contributions from both receptors.

Other studies investigating the effects of β-adrenergic signaling on immunity align with the data presented herein. One such study denoted that β-adrenergic signaling disrupts NF-κB pathways and CD40 signaling in dendritic cells in a manner that was similarly reversed by treatment with propranol^97^. Another group described the myeloid-directed anti-tumor activity of an α2 adrenergic agonist, although the authors examined extracranial tumors only^98^. Two additional studies highlighted the ability of β-adrenergic blockade to mitigate either the number^99^ or activity^100,101^ of immunosuppressive myeloid cells within the TME, much as we observed as well.

β-adrenergic receptor expression has been detected on melanoma^71–74^ and other cancers^74–77^, with a direct role in tumor progression suggested. As a result, β-adrenergic blockade has been previously investigated in peripheral cancers for its direct antitumor effects as a monotherapy^72,102,103^, with conflicting results^104–107^. Further, several studies have examined the direct inhibitive effects of propranolol on glioma cell lines *in vitro* (predominantly rat)^108–111^, although our data reveal relatively low expression of β-adrenergic receptors on glioma cells compared to the immune compartment. Studies in peripheral cancers have examined the local immune impact of adrenergic signaling within the TME^102,103^, generally positing a role promoting T cell exhaustion^58,101,112–115^. We, however, report a larger array of β-adrenergic-mediated immune deficits amidst intracranial cancers, which are likely attributable to the chronic systemic sympathetic hyperactivity that we also uncover in the context of these tumors. Such chronic sympathetic hyperactivity also quite likely explains the added benefit to β-adrenergic blockade seen in the intracranial tumor setting.

Our study reports extended survival in GBM patients having received β-adrenergic blockade, as well as in patients with melanoma and lung cancer brain metastases who received β-blockade alongside concomitant ICI. While β-blockade also impacts outcomes in settings of exclusively extracranial disease, the benefits are especially pronounced in patients harboring an intracranial disease burden. Our studies examining patients with brain metastases are particularly relevant, as brain metastases are now 10 times more prevalent than GBM, with lung cancer and melanoma exhibiting two of the highest rates of intracranial metastasis^116,117^. In melanoma, studies have previously examined the combination of β-adrenergic and checkpoint blockade^72,78^, but these have frequently excluded patients with stage IV disease, and thus, brain metastases^101,118–120^. One study has previously investigated the impact of β-adrenergic blockade in the setting of GBM, assessing outcomes retrospectively in patients with recurrent GBM (n=243) and revealing no association between β-blockade and overall survival^121^. In contrast, our analysis of the SEER-Medicare database investigated a cohort of 8743 GBM patients (35x larger) and included those who were newly diagnosed.

In our analysis of the SEER-Medicare data, we attempted to list all reasonable exclusions. The nature of retrospective data, however, is such that it is not possible to account for all potential confounders and caveats remain. Underlying differences in a patient’s past medical and diagnostic history could account for a component of the survival benefit associated with β-blocker usage, as could the impact of β-blockers on those conditions. We would expect some of this variability to be captured in the patient comorbidity score that is included in all multivariate models shown. Additionally, care must be taken when interpreting the use of claims data for metastasis research.^122^ Use of ICD-9 or ICD-10 to identify patients with asynchronous brain metastases will be imperfect due the nature of diagnostic claims data as, while SEER-Medicare contains nearly all claims billed through Medicare, previous research has shown that the information contained in the claims themselves is not always reliable or valid.^123–126^ Threats to the validity of claims may include under-/over-diagnosing, patient compliance with billed treatments, and inconsistent use of billing codes, among others. Prior evaluations have demonstrated that use of tumor-specific algorithms have ∼55% sensitivity and 85% specificity for lung cancer, suggesting that individuals with metastasis will be missed but those assigned as having metastatic disease are likely correctly identified. These algorithms will result in the underestimation of total asynchronous brain metastases.^127^ It was also not possible to determine further details of β-blocker usage, beyond their being billed to Medicare. Together, this makes our data, as with all retrospective data, subject to the bias of indication. While our data examining survival in patients with GBM and intracranial metastases are therefore limited in scope by this bias, the prospective complement proffered by our murine survival and immune data strongly suggest a specific benefit to combining β-adrenergic blockade and immunotherapy in the setting of intracranial tumors.

The data presented herein support the initiation of prospective randomized controlled trials to examine the addition of β-adrenergic blockade to immunotherapeutic regimens in patients with intracranial cancers, whether primary or metastatic. Such trials, we anticipate, will be viewed favorably, as β-blockers are FDA-approved, relatively inexpensive, and have exhaustively established safety profiles^128^. Repurposing β-blockers as an immunotherapeutic adjunct may ultimately represent a novel intervention for licensing long sought-after effective immune responses within the intracranial compartment.

## METHODS

### Mice

The Institutional Animal Care and Use committee (IACUC) at Duke University Medical Center approved all experimental procedures. Female C57BL/6 mice purchased from Charles River Laboratories were purchased at 5-6 weeks of age. Animals were maintained under specific pathogen-free conditions at the Cancer Center Isolation Facility of Duke University Medical Center. All experimental procedures were approved by the IACUC.

### Cell lines

Murine CT-2A malignant glioma was kindly provided by Robert L. Martuza (Massachusetts General Hospital, MA, USA). LLC1/2 lung adenocarcinoma were purchased from the American Tissue Culture Collection (ATCC). Murine B16F0 melanoma was obtained from the Duke Cell Culture Facility but is also available from ATCC. All cell lines are syngeneic on the C57BL/6 background. CT-2A and LLC1/2 were grown *in vitro* in Dulbecco’s Modified Eagle’s Medium (DMEM) with 2mM L-glutamine and 4.5mg ml^-1^ glucose (Gibco) containing 10% fetal bovine serum (FBS) (Sigma). B16F0 cells were grown *in vitro* in DMEM with 4mM L-glutamine, 4.5mg ml^-1^ glucose, 1mM sodium pyruvate, 1500 mg/L sodium bicarbonate containing 10% FBS (Sigma). Cells were collected in the logarithmic growth phase. All cell lines were authenticated and tested negative for mycoplasma and interspecies contamination by IDEXX Laboratories.

### Tumor Implantation

For intracranial implantation, tumor cells were resuspended at a given concentration in PBS, then mixed 1:1 with 3% methylcellulose and loaded into a 250mL syringe (Hamilton). Prior to injection, mice were anesthetized with isoflurane and subject to an anteroposterior incision to reveal the skull. The needle was positioned 2mm to the right of the bregma and 4mm below the surface of the skull at the coronal suture using a stereotactic frame. 1e^4^ CT-2A, 1e^3^ B16F0 or 1e^3^ LLC1/2 cells were delivered in a total volume of 5mL per mouse. For subcutaneous implantation, 5e^5^ CT2A, 2.5e^5^ B16F0, or 5e^5^ LLC1/2, were injected in a total of 200mL of PBS into the subcutaneous tissues of the left flank.

### Catecholamine and Corticosterone Analysis

De-identified human samples were used in this study with exempt Institutional Review Board (IRB) status granted from the Duke University Health System IRIB (protocol ID: Pro00112766). De-identified serum samples from 5 female and 5 male GBM patients were collected at time of recurrence, processed immediately, and stored in vapor phase liquid nitrogen. De-identified serum samples from 5 newly diagnosed GBM patients (3 males, 2 females) were collected at time of diagnosis, likewise processed immediately, and stored similarly. Serum from patients with both newly-diagnosed and recurrent GBM was collected during a pre-operative clinic visit. Patient serum samples were obtained from patients at Duke University (protocol # Pro00007434, BTBR). De-identified healthy control human serum was obtained from BioIVT (5 males and 5 females, HUMANSRM-0001254 and HUMANSRM-0001253, respectively). Healthy control human serum was obtained according to the BioIVT Standard Operating Procedures. Briefly, after whole blood was drawn into a dry collection bag, it was sun at 4,500-5,000 x g for 10-15min in a refrigerated centrifuge. Supernatant was then transferred into another bag and allowed to clot at room temperature. The resulting material was spun at 3,500-5,000 x g for 15-20min in a refrigerated centrifuge and subsequently aliquoted.

Glioblastoma patient serum was obtained according to the standard operating procedure for clinical samples at Duke University. Briefly, whole blood was allowed to clot for up to 60min then centrifuged at 2200 RPM for 10min at room temperature. Serum was aliquoted and stored at - 80C and subsequently transferred to liquid nitrogen for long-term storage.

Blood and spleen were collected from non-tumor bearing mice or mice with intracranial tumors or subcutaneous tumors at respective humane endpoints. To mitigate the differential impact of stress on mice, all mice were handled as minimally as possible and were immediately anesthetized using isoflurane, with only one cage (n=5 mice) handled at any given time. They were then subject to retro-orbital bleed and subsequent euthanasia and tissue harvest. All samples were stored on ice and processed immediately, with total processing time not exceeding 1hr. Serum samples and spleen supernatant from all groups were processed simultaneously. Blood samples were centrifuged at 2000g for 15min at 4°C to isolate plasma, which was isolated in 58.8mL aliquots with 1.2mL 50mM EDTA 200mM sodium metabisulfite per sample to protect against catecholamine degradation. Spleens were collected into 500mL of a PBS/ 50mM EDTA 200mM sodium metabisulfite solution, weighed, homogenized through 70mm filters, washed with an addition 500mL of PBS/EDTA/sodium metabisulfite solution for a total of 1mL, then centrifuged at 500g for 5min at 4°C. Supernatants were collected by combining 58.8mL supernatant and 1.2mL 50mM EDTA 200mM sodium metabisulfite into Eppendorf tubes.

All samples, including human serum samples were stored at −80°C. Epinephrine and norepinephrine levels in plasma, serum, and spleen supernatant were analyzed using the 2-CAT (A-N) Research ELISA (Rocky Mount Diagnostics).

### Murine tissue harvest

Tumor, spleen, thymus, inguinal or deep cervical lymph node, and tibia were collected at defined and/or humane endpoints, in accordance with protocol. For intracranial tumor-bearing animals, humane endpoints include inability to ambulate two steps forward with prompting. For subcutaneous tumor-bearing animals, humane endpoints include tumor size greater than 20 mm in one dimension, 2000mm^3^ in total volume, or tumor ulceration or necrosis. Spleens, lymph nodes and thymi were weighed. Briefly, tissues were processed in RPMI or PBS, minced into single cell suspensions, cell-strained, counted, stained with antibodies, and analyzed via flow cytometry.

Tumors were mechanically homogenized using a Dounce Tissue Homogenizer and enzymatically digested using 0.05mg ml^-1^ Liberase TL (Roche), 0.2mg ml^-1^ Dnase I (Roche) in HBSS with calcium and magnesium. Tumors in digestion buffer were then incubated at 37°C for 15min, PBS was used to quench the enzymatic reaction, and tumors were filtered (70μm). After centrifugation, cells were resuspended in 1x RBC Lysis Buffer (eBioscience) for 3min and quenched with PBS. Samples were subsequently centrifuged then resuspended in 30% Percoll (Sigma Aldrich) and centrifuged at 500g for 20min at 18°C with a low brake. The myelin-containing supernatant was then aspirated, and the cell pellet resuspended in PBS for staining.

Blood was collected into EDTA tubes and mixed thoroughly to prevent coagulation. After collection, 75μL of each blood sample combined with 1mL 1x RBC Lysis Buffer (eBiosciences) and incubated for 3min then quenched with PBS. RBC lysis was performed twice before proceeding to staining.

### Single-cell RNA sequencing analysis

Integration of a single cell atlas and downstream analyses were performed in Python (v3.9.13) using Scanpy (v1.9.1) and scvi-tools (v0.17.4). Filtered feature gene matrices from the following datasets were queried from the gene expression omnibus or provided by the original authors: GSE182109 (2 low grade glioma, 10 newly diagnosed GBM, 5 recurrent GBM)^129^, GSE154795 (14 newly diagnosed GBM, 12 recurrent GBM, 5 PBMCs from recurrent GBM, 2 healthy donor PBMCs)^130^, GSE160189 (5 normal brain)^131^, GSE200218 (5 melanoma brain metastases)^132^, and GSE186344 (3 breast cancer brain metastases, 3 lung cancer brain metastases, and 3 melanoma brain metastases)^133^.

Matrices were read into Python with sc.read_10x_mtx(), and relevant metadata information was added (patient of origin, study of origin, tumor type). Anndata objects from each dataset were then concatenated with adata.concatenate() with join=outer. QC metrics were calculated and cells with >8000 genes by counts, <500 total counts, and >25 percent mitochondrial DNA were removed. A normalized counts layer for downstream analysis was created with sc.pp.normalize_total(), with a target sum of 10,000. 5000 highly variable genes were selected using scvi.data.poisson_gene_selection() with batch_key=patient of origin, and using the raw counts matrix, as is required for scvi. The model was then constructed using the filtered anndata object containing only highly variable genes with scvi.model.SCVI.setup_anndata(). Again, the raw counts layer was used, with the batch_key set to patient of origin, and percent mitochondrial counts used as continuous categorical covariate. Because we were interested in primarily distinguishing between healthy cells, tumor cells, and immune cells, and not specific tumor types, tumor type was used as a categorical covariate in the model. The model was run with model.train() with an early stopping patience of 20 and 700 maximum epochs. The latent representation of the model was obtained with model.get_latent_representation() and added to .obsm of the original anndata object (containing all genes). For clustering, nearest neighbors were calculated using the .obsm latent representation from the model and n_neighbors=15, followed by sc.tl.umap() with min_dist=0.5 and sc.tl.leiden() with resolution=2.0. Clusters were analyzed individually with log1p total counts and percent mitochondrial counts. The model tends to cluster low quality cells (high mitochondrial content, high total counts, or low total counts) together, allowing for further quality control, rather than using a strict initial cutoff by QC metrics prior to visualizing data and cell types^134^. Expression of known marker genes from each tumor type, normal cell type, and immune cell components were evaluated by plotting gene expression onto the UMAP, using the normalized counts layer. We also assessed marker genes per leiden cluster with sc.tl.rank_genes_groups(), method=Wilcoxon, using the normalized counts layer. If a leiden cluster had exceptionally high or low total counts, or high mitochondrial counts, and did not align with known tumor, normal cell, or immune marker combinations (by UMAP gene expression and Wilcoxon Rank-Sum cluster markers), it was filtered out as low-quality cells or likely doublets. Each time leiden clusters were removed, another model was retrained with the same parameters, starting at the highly variable gene selection step and progressing through cluster quality control, until there were no clear low quality cell clusters remaining. Notably, in order to minimize bias in downstream analysis, expression of adrenergic receptors was not considered in any of the quality control and filtering steps.

All UMAP plots were generated with sc.pl.umap(), and clusters were annotated manually based on combinations of known marker genes. Matrix plots and Dot plots were generated with sc.pl.matrixplot() and sc.pl.dotplot(), respectively, with the normalized counts layer in both cases. For generating disease specific UMAPs, the global anndata object was subset based on patient metadata, and the primary UMAP embedding was retained to preserve graph architecture, allowing for easier comparison between subset and data objects.

### *In vivo* Treatment

For immunotherapy administration, 200μg 4-1BB agonist or vehicle control was administered intraperitoneally in 200μL PBS every 3 days, beginning at day 9, through day 18 post-tumor implantation for a total of 4 treatments.

For propranolol administration, mice were given 0.5g/L propranolol or vehicle control in drinking water *ad libitum* beginning on the day of tumor implantation.

For epinephrine/norepinephrine, dobutamine (β1-adrenergic receptor agonist) and salbutamol (β2-adrenergic receptor agonist) administration, naïve mice were implanted with subcutaneous osmotic pumps (Alzet) for seven days. Mice received a combination of 10mg/kg/day of epinephrine and 7mg/kg/day of norepinephrine, or 2mg/kg/day of dobutamine or 8mg/kg/day of salbutamol.

### Western Blot Analysis

CD45^+^ immune cells were isolated from CT2A tumor bearing mouse brains. Single cell suspensions were made, and cells were counted on a hemocytometer. CD45^+^ immune cells were then isolated using CD45 positive selection kit (Stem Cell, cat# 100-0350) according to the manufacturer’s instructions. Isolated immune cells were washed with PBS and spun down at 400g for 5 minutes. Cell pellet was placed on ice and resuspended in 200μL of lysis buffer. Lysis buffer was made as following: 10x lysis buffer (Cell Signaling, cat# 9803S) was diluted in ddH2O supplemented with 1mM PMSF (200x stock solution was purchased and reconstituted with 1ml of isopropanol-Cell Signaling, cat# 8553S), and 1x protease Protease/Phosphatase Inhibitor Cocktail (100X stock solution cat# 5872S). Samples were incubated on ice for 5min and then rapidly sonicated in an ice bath sonicator. Samples were then spun at 14000g at 4C and the protein content of the supernatant was calculated using a BCA assay (Pierce, Cat # 23227). Samples were frozen at −20 C until the day assay was performed. Samples were diluted in 2x Laemmli Sample Buffer (Biorad, cat #1610737) supplemented with 2-Mercaptoethanol (Biorad, cat# 1610710 2-Mercaptoethanol). 950μL of 2x Laemmli buffer was mixed with 50μL of 2-Mercaptoethanol to obtain 2x sample buffer. Samples were loaded based on protein concentration and adjusted with lysis buffer supplemented with PMFS and protease/phosphatase inhibitor. Samples were then mixed with the 2x sample buffer in a 1:1 ratio. Samples were boiled at 95 C for 10min, and immediately cooled on ice for 5 mins before being loaded into a 4-20% gradient gel (Invitrogen, cat #XP04205BOX). Gel electrophoresis was performed at 100 constant volts until samples reached the bottom. Precision Plus Protein™ Kaleidoscope™ Prestained Protein Standards was used as a ladder (Biorad #1610375). PVDF membrane (Biorad, cat# 1620177) was activated in 100% methanol for 1min, then placed in ddH2O for 1min, and rested in 1x transfer buffer with 20% methanol (10x transfer buffer Cell Signaling, 12539S). 1x transfer buffer contained 100ml of 10x transfer buffer, 700ml of ddH2O, and 200ml of methanol. An overnight wet tank transfer was performed using 1x transfer buffer with 20% methanol using Mini Trans-Blot® Cell wet tank transfer system (Biorad, cat# 1703930) at 25 constant volts. Membrane was blocked with SuperBlock (Thermofisher, cat# 37515) for 90-120 minutes at 4°C on a shaker. Membranes were then cut and incubated with primary antibody and incubated overnight in 10mL of a mixture of SuperBlock and 1XTBST plus 10mL of primary antibody. The p65 antibody was from the NF-κB Pathway Antibody Sampler Kit (Cell Signaling, cat #9936). Antibodies against Histone H3 were used as loading control (Cell Signaling, cat # 4499S). Membranes were washed 5 times with 1x TBST (Cell Signaling, cat # 9997S) with gentle agitation on a shaker at room temperature. Membranes were then incubated with a mix of anti-rabbit and anti-mouse HRP conjugated secondary antibodies according to the instructions provided with NF-κB Pathway Antibody Sampler Kit in a mix of Superblock and 1x TBST for 60-90min on a shaker at room temperature. Membranes were washed 4 times in 1xTBST and once in 1x TBS (Cell Signaling, cat# 12498S) on a shaker at room temp. Membranes were developed using SuperSignal™ West Pico PLUS Chemiluminescent Substrate (Thermofihser, cat# 34580). Images were acquired and quantified on a Licor Odyssey FC running Image Studios software.

### *In vivo* Imaging for Tumor Growth

Mice were implanted with 50K CT2A-Luciferase, ear-tagged such that individual mice could be followed, and imaged twice weekly, starting at day 10 after tumor implantation. Luciferin was resuspended at 15mg/mL. Mice were weighed then injected with a 150mg/kg luciferin, in groups of 2-3 mice at a time. Five minutes after luciferin injection, all mice were anesthetized in an isoflurane chamber. After an additional 2min, mice were transferred into the In Vivo Imaging System (IVIS) for imaging. Images were analyzed upon termination of the experiment where total flux (radiance) was measured for all mice across timepoints after images were adjusted to reflect the same scale on every image.

### Flow Cytometry

Cells were first washed with PBS then resuspended in Zombie Aqua Viability Dye (1:500 in PBS, Biolegend). For extracellular staining, samples were first resuspended in 50mL TruStain FcX Plus (1:500, Biolegend) in FACS buffer (2% FBS in PBS) for 15min at room temperature to block Fc receptors. After blocking, 50mL of a 2x antibody mastermix was added to samples and samples were incubated for 30min at 4°C. Samples were then washed in FACs buffer and fixed with 2% formaldehyde in PBS for 20min at 4°C or fix/permeabilized for subsequent intracellular staining.

For intracellular staining, samples were fixed and permeabilized with a 1X FoxP3 Fixation/Permeabilization Buffer (eBioscience FoxP3/Transcription Factor Staining Buffer Set, Thermo Fisher Scientific) for 30min at 4°C.

Prior to sample acquisition, 10mL of Count Bright counting beads (Thermo Fisher Scientific) in 40mL of FACs buffer were added to each sample. Samples were acquired on a Fortessa (BD Biosciences) using FACS Diva software v.9 and analyzed using FlowJo v.10 (Tree Star).

Information regarding antibodies can be found below:

| Marker | Color | Company | Catalog # | Clone |
| --- | --- | --- | --- | --- |
| Live/Dead | BV510 | Biolegend | 423102 | NA (Dye) |
| CD45 | APC-Cy7 | Biolegend | 103116 | 30-F11 |
| CD3 | PE-Cy5 | Biolegend | 100310 | 145-2C11 |
| CD19 | PE-Cy5 | Biolegend | 115509 | 6D5 |
| NK1.1 | PE-Cy5 | Biolegend | 108715 | PK136 |
| Ly6G | PE-Cy5 | Invitrogen | 15-9668-82 | 1A8-Ly6g |
| CD64 | BV711 | Biolegend | 139311 | X54-5/7.1 |
| CD11c | BV786 | Biolegend | 117335 | N418 |
| CD11b | PE | Biolegend | 101207 | M1/70 |
| Siglec-H | PE-Cy7 | Invitrogen | 25-0333-80 | eBio440c |
| MHCII | AF488 | Biolegend | 107615 | M5/114.15.2 |
| CD8 | BV650 | Biolegend | 100742 | 53-6.7 |
| CD103 | PE-Dazzle | Biolegend | 121429 | 2E7 |
| XCR1 | BV421 | Biolegend | 148216 | ZET |
| CD40 | BV605 | BD Biosciences | 745218 | 23-Mar |
| CD86 | APC | Biolegend | 105011 | GL-1 |
| CD3 | PE | BD Biosciences | 553064 | 145-2C11 |
| CD4 | BV605 | Biolegend | 100548 | RM4-5 |
| CD8 | APCR700 | BD Biosciences | 564983 | 53-6.7 |
| CD11b | PE Dazzle | BD Biosciences | 562287 | M1/70 |
| Ly6C | BV650 | Biolegend | 128049 | HK1.4 |
| Ly6G | BV421 | Biolegend | 127627 | 1A8 |
| F4/80 | PE-Cy5 | Biolegend | 123112 | BM8 |
| MHCII | AF488 | Biolegend | 107615 | M5/114.15.2 |
| FoxP3 | PE-Cy5.5 | Biolegend | 320012 | 150D |
| CD44 | BV786 | Biolegend | 103059 | N418 |
| PD1 | PE-Dazzle | Biolegend | 109116 | RMP1-30 |
| CD25 | BV421 | Biolegend | 102043 | PC61 |
| CD40L | PE-Cy7 | Biolegend | 106512 | MR1 |
| CD62L | PE-Cy5 | Biolegend | 104410 | MEL-14 |
| KLRG1 | APC | Biolegend | 138412 | 2F1/KLRG1 |
| CD69 | BV711 | Biolegend | 104537 | H1.2F3 |
| FoxP3 | AF488 | Biolegend | 320012 | 150D |
| CD8 | BUV737 | BD<br>Biosciences | 612759 | 53-6.7 |
| iNOS | PE-Cy5.5 | Invitrogen | 35-5920-80 | CXNFT |
| CD68 | PE-Cy7 | Biolegend | 137015 | FA-11 |
| F4/80 | PE-Cy5 | Invitrogen | 15-4801-82 | BM8 |
| CD206 | PE-Dazzle<br>594 | Biolegend | 141731 | C068C2 |
| Arg1 | PE | Invitrogen | 12-3697-80 | A1exF5 |
| CD4 | APC-Cy7 | Biolegend | 100526 | RM4-5 |
| CD11c | APCR700 | BD<br>Biosciences | 565872 | N418 |
| TOX | EF600 | Invitrogen | 50-6502-82 | TXRX10 |
| MHCII | AF488 | Biolegend | 107616 | M5/114.15.2 |
| PDL1 | BV785 | Biolegend | 124331 | 10F.9G2 |
| CD19 | BV750 | Biolegend | 115561 | 6D5 |
| PD1 | BV711 | Biolegend | 135231 | 29F.1A12 |
| TIM3 | BV421 | Biolegend | 119723 | RMT3-23 |
| SLAMF6 | BV650 | BD<br>Biosciences | 740628 | 13G3 |
| Ly6G | BV605 | BD<br>Biosciences | 563005 | 1A8 |
| Ki67 | BUV805 | Invitrogen | 368-5698-82 | SolA15 |
| CD11b | BV480 | Invitrogen | 414-0112-82 | M1/70 |
| CD3 | BUV395 | BD<br>Biosciences | 563565 | 14-2C11 |
| CD45 | BUV496 | Invitrogen | 364-0451-82 | 30-F11 |
| Ly6C | BUV563 | BD<br>Biosciences | 755198 | HK1.4.rMab |
| CD44 | BUV615 | Invitrogen | 366-0441-82 | IM7 |

### Single-Cell Cytokine Analysis

Spleens were harvested and T cells were isolated via untouched CD8^+^ T cell isolation or Pan T cell isolation kits (Miltenyi) and stimulated ex vivo with plate-coated anti-CD3, soluble anti-CD28 antibodies (Invitrogen) for 16hours. T cells were then harvested and loaded into single-cell adaptive immune chips according to manufacturer instructions. Chips were run on the Isolight machine (Isoplexis) overnight and data were analyzed using IsoSpeak software (Isoplexis).

### Immunofluorescence Imaging

Paraffin-embedded sample blocks of murine CT2A were utilized to prepare 5μm tissue sections on glass slides. One slide per sample was stained with a b2-adrenergic receptor (Abcam) primary antibody at 1:500 concentration. Slides were also stained with DAPI (1:20). One slide per sample was stained using non-immune rabbit and rat IgG (Invitrogen). Immunofluorescence was measured using a Leica SP5 inverted confocal microscope. Adrenergic receptor and nucleus expression were quantified using Qupath computer software. β2-adrenergic receptor expression was normalized to IgG control and DAPI expression.

### Gene Expression Analysis

C57BL/6 mice were implanted with intracranial CT2A glioma and treated with propranolol or vehicle control as described above. An additional non-tumor bearing group was included. Tumors were harvested and combined (n=5 tumors per group) with n=3 biological replicates per sample for a total of 15 tumors per sample. Immune cells were FACS-sorted for APC-enriched samples or T cell-enriched samples and RNA was subsequently extracted (RNeasy Mini Kit, Qiagen). RNA was analyzed on an nCounter MAX Analysis System (Nanostring) with the PanCancer Immune Profiling panel (Nanostring) according to manufacturer instructions. Expression data were analyzed using Rosalind, nSolver and nSolver Advanced Analysis software.

### Human Retrospective Data

SEER is a population-based cancer registry that collects data on newly diagnosed cancers and provides cancer statistics for these cancers. SEER data are aggregated from cancer registries and contain patient demographic and tumor characteristics as well as the first course of treatment. SEER-Medicare^135,136^ is the result of a linkage between existing SEER case data and detailed information about Medicare beneficiary’s covered healthcare services from the time of Medicare eligibility until death. Importantly, this database only includes individuals who are eligible for Medicare, which, in the U.S., generally occurs at age 65. Individuals may be eligible for Medicare at younger ages due to reasons such as disability or certain medical conditions, but these individuals were excluded from this analysis due to the potentially confounding effects of their qualifying conditions (ie. kidney failure necessitating dialysis). The linked dataset includes 15 SEER registries covering ∼24% of the US population. Medicare included approximately 95% of patients age 65 and older in the SEER files^136^. Additionally, among all Medicare enrollees, the proportion of those enrolled in Medicare Advantage plans has increased from 21% in 2008 to 34% in 2018.^137^ As enrollees in Advantage plans will lack claims for the time period they are enrolled, this restriction represents a significant limitation to the population that is able to be used. Furthermore, as some patients may have private health insurance or not be continuously enrolled in Medicare Part B in addition to Part A, there is a possibility that some included individuals may have an incomplete claims history. Individuals were identified in the SEER cancer files for diagnoses made from January 1, 2008 – December 1, 2017, excluding patients younger than 65 years and those enrolled due to end-stage renal disease or disability **(ICD codes in Supplementary Table 1**). Patients enrolled in a Health Maintenance Organization six months prior and twelve months after primary diagnosis were excluded (34.1% of lung cancer, 31.3% of melanoma, 24.3% of GBM). Individuals with missing age, race, ethnicity, sex, and survival or follow-up time were excluded from the analysis. These exclusions were made to assure completeness of claims data in the analytic dataset.

Cases of primary GBM (International Classification of Diseases for Oncology 3^rd^ Edition (ICD-O-3)^138^ morphology codes 9440/3, 9441/3, 9442/3, 9445/3; topographic codes C71.0-C71.9), melanoma (ICD-O-3 morphology codes 8720/3-8790/3, ICD-O-3 topographic codes C44.0-C44.9) and lung cancer (ICD-O-3 morphology codes 8000/3-8579/3, ICD-O-3 topographic codes C34.0-C34.9) were identified in the SEER cancer file. Those with a prior cancer diagnosis, which was identified using a SEER-provided variable (*Sequence_number)*, were also excluded (36.9% of lung cancer, 53.9% of melanoma, 32.8% of GBM).

Intracranial metastases were identified using the SEER synchronous brain metastasis variable and the Medicare Provider Analysis and Review (MEDPAR), National Claims History (NCH), and Outpatient (OUTPAT) claims files as previously described in Ascha, et al.^127^ Briefly, patients. Patients with intracranial metastases were defined as those with a diagnosis code for an intracranial metastasis (International Classification of Diseases 9^th^ and 10^th^ Ed. (ICD-9 and ICD-10) codes 198.3 and C79.3, respectively) and at least one procedure code for a brain or head diagnostic test within 60 days of the intracranial metastasis diagnosis (Current Procedural Terminology^139^ codes 70450-70470, 70551-70553, 78607-78608). Prior comorbidity score was estimated using the rule-out option of the SEER-Medicare SAS comorbidity macro (2021 version)^140^ using both ICD-9 and ICD-10 diagnosis codes. The comorbidity scores were calculated with Charlson weights and grouped as a score of 0, 1, 2, and 3+. Cancer treatment patterns – including radiation, chemotherapy, β-blocker, and checkpoint blockade usage (including Pembrolizumab, Necitumumab, Nivolumab, Ipilimumab, Atezolizumab, and Ramucirumab) – were identified using both SEER data and the MEDPAR, NCH, OUTPAT, Part D Event, and Durable Medical Equipment files. Surgery was identified using SEER -provided variable (*RX_Summ_Surg_prim-Site_1998)* which includes biopsies with excision and claims with CPT and ICD codes specific to the type of primary cancer (**Supplementary Table 1**). Radiation therapy and chemotherapy were defined by ≥2 claims for the same code separated by at least one week, and prior use of β-blockers was defined as ≥2 dispensing for the same β-blocker prior to diagnosis (**Supplementary Table 1**). Patients that started checkpoint blockade before detection of intracranial metastasis were excluded from the analysis.

Cases of lung and melanoma that have been determined to have brain metastases include those that are synchronous (found at initial diagnosis) and asynchronous (those found more than 1 month after diagnosis). Synchronous brain metastases were identified via a combination of the SEER-provided brain metastasis variables and a previously validated algorithm to detect brain metastases in claims data, while asynchronous brain metastases will be identified solely through the latter. Patients with asynchronous brain metastases may not be identified as stage IV at the initial diagnosis of their primary due to the follow-up information available through Medicare claims. Cases with extracranial metastases at diagnosis were identified using the SEER-provided stage at diagnosis variable only and did not include cases identified as having brain metastasis.

Sensitivity analyses showing the effects of concomitant β-blocker and checkpoint blockade in patients with lung cancer and melanoma that were identified as having continuous enrollment in Medicare Part A and B, as well as equivalent months of enrollment in Medicare Part D, showed no substantial differences in effect size from the unrestricted cohort.

### Statistical Analyses

Experimental group assignment was determined by random designation. All measurements were taken from distinct samples. The statistical tests employed for each data presentation and number of samples (n) are designated in the respective figure legends. Individual data points are represented as dots in graphs. Statistical analyses were performed using GraphPad Prism software. Error bars represent the standard error of the mean (SEM). No data were excluded from analysis. Mice were randomized into treatment groups after tumor injection. P-values <0.05 were considered statistically significant. Investigators were not blinded to group assignment during experimental procedures or analysis, but survival studies were monitored both by investigators and additionally monitored by veterinary staff from the Duke animal facility who were blinded to the studies and reported humane endpoints accordingly.

Analyses of clinical data were performed stratified by cancer type within selected treatment groups. Kaplan-Meier estimation was used to assess the effect of a combination β-blocker and checkpoint blockade versus checkpoint blockade alone on overall survival. In GBM, where no checkpoint inhibitors are currently approved for use, the effect of β-blockers on overall survival was examined. Multivariable Cox proportional hazards models were used to assess the relationship between combination β-blocker and checkpoint blockade versus checkpoint blockade alone among the cohorts, adjusting for SEER-provided demographics and potential confounders, including age; sex (male or female); and race/ethnicity (non-Hispanic White, non-Hispanic Black, non-Hispanic Other, or Hispanic [all races], or Unknown); treatment pattern (yes/no surgery of primary tumor, radiation and chemotherapy); and prior comorbidity score. Sensitivity analyses were performed in the subset of individuals identified as having continuous enrollment in Medicare Part A and B, as well as equivalent months of enrollment in Medicare Part D as Medicare Part A and B. P-values <0.05 were considered statistically significant. All analyses of clinical data were conducted using R version 4.1.3 using the following packages: gtsummary, patchwork, survival, survminer, and tidyverse suite.^141–145^

## Ethics statement

All studies performed in research animals and on clinical samples are compliant with all relevant ethical regulations. All animal experiments were approved by the IACUC at Duke University Medical Center (protocol A163-21-08). De-identified human samples were used in this study with exempt Institutional Review Board (IRB) status granted from the Duke University Health System Institutional Review Board (protocol ID: Pro00112766).

## DATA AVAILABILITY

All scRNAseq datasets used for analysis are publicly available: GSE182109^129^, GSE154795^130^, GSE160189^131^, GSE200218^132^, and GSE186344^133^. SEER-Medicare data are publicly available, but these data are released on a project-specific basis that requires an approved application. Applications are reviewed by NCI-led content experts and application revisions may be required. Initial reviewer comments are typically provided within 2-3 weeks. Fees are incurred to cover the cost of creating the requested data files. Instructions for the application can be found here: https://healthcaredelivery.cancer.gov/seermedicare/obtain/requests.html#docs.

Gene expression data that support the findings of this study have been deposited in the Gene Expression Omnibus under the accession number GSE241720.

## CODE AVAILABILITY

Integration and analysis of publicly available scRNAseq datasets were conducted based on publicly available workflows and can be accessed at https://github.com/jbfinlay/Lorrey_Wachsmuth_et_al. Analysis of relevant SEER-Medicare data can be accessed at https://zenodo.org/record/8289570.

## Author Contributions

SJL and LPW conceived and designed experiments. SJL, LPW, KA, JWP, AHM, XC, AH, LR, RR, acquired data. SJL, LPW, JBF, KA, SD, MP, CN, QTO analyzed data. SJL, LPW and PEF interpreted data. SJL, LPW, CN, QTO, and PEF wrote the manuscript. EL, DW, ES, KMW, AP provided feedback.

## Acknowledgements

Dr. Fecci is the recipient of a CRI Arash Ferdowsi Lloyd J. Old STAR grant, which helped to fund this work. We thank Gary Archer from the Duke University Brain Tumor Center for providing human serum samples. We further thank Amanda van Swearingen from the Duke University Center for Brain and Spine Metastases for IRB protocol assistance. We thank the Duke University School of Medicine for the use of the Microbiome Core Facility that provided the Nanostring Gene Expression service. This study used the linked SEER-Medicare database, and we thank Zhen Liu for her assistance with data analysis. The interpretation and reporting of these data are the sole responsibility of the authors. The authors acknowledge the efforts of the National Cancer Institute; Information Management Services (IMS), Inc.; and the Surveillance, Epidemiology, and End Results (SEER) Program tumor registries in the creation of the SEER-Medicare database. The collection of cancer incidence data used in this study was supported by the California Department of Public Health pursuant to California Health and Safety Code Section 103885; Centers for Disease Control and Prevention’s (CDC) National Program of Cancer Registries, under cooperative agreement 1NU58DP007156; the National Cancer Institute’s Surveillance, Epidemiology and End Results Program under contract HHSN261201800032I awarded to the University of California, San Francisco, contract HHSN261201800015I awarded to the University of Southern California, and contract HHSN261201800009I awarded to the Public Health Institute. The ideas and opinions expressed herein are those of the author(s) and do not necessarily reflect the opinions of the State of California, Department of Public Health, the National Cancer Institute, and the Centers for Disease Control and Prevention or their Contractors and Subcontractors. Schematics in Fig.1, 3 and 4 were created with BioRender.com.

