## Supplementary Figures for "Glioblastoma and other intracranial tumors elicit systemic sympathetic hyperactivity that limits immunotherapeutic responses"

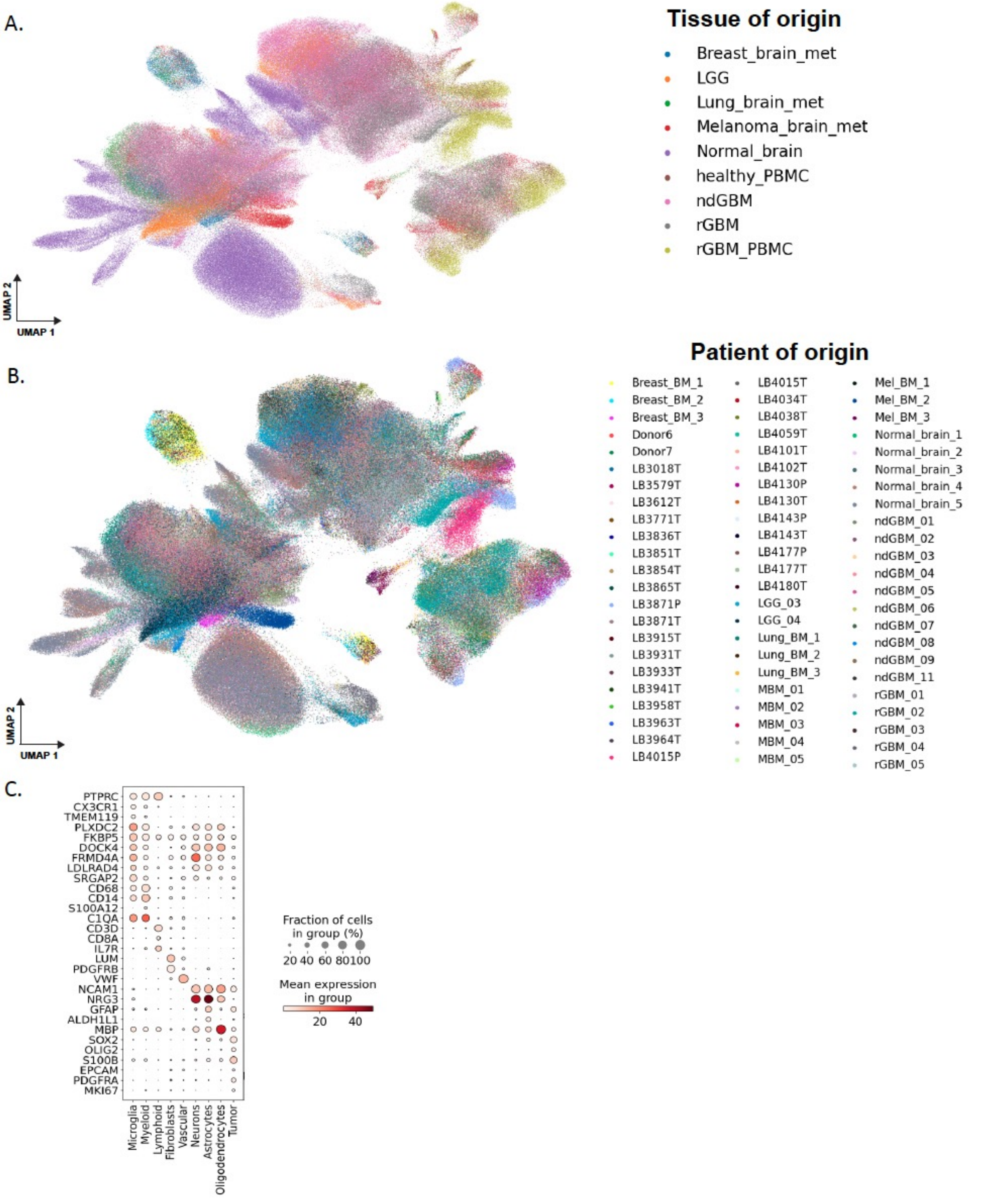


**Supplementary Figure 1. Integration and cell type annotation of publicly-available human scRNAseq datasets.** A. UMAP of integrated scRNAseq datasets from normal brain, GBM, brain metastasis and PBMC samples labelled by tissue type. B. UMAP of integrated scRNAseq datasets from normal brain, GBM, brain metastasis and PBMC samples labelled by patient sample. C. Expression of signature population-defining genes in annotated clusters**.**

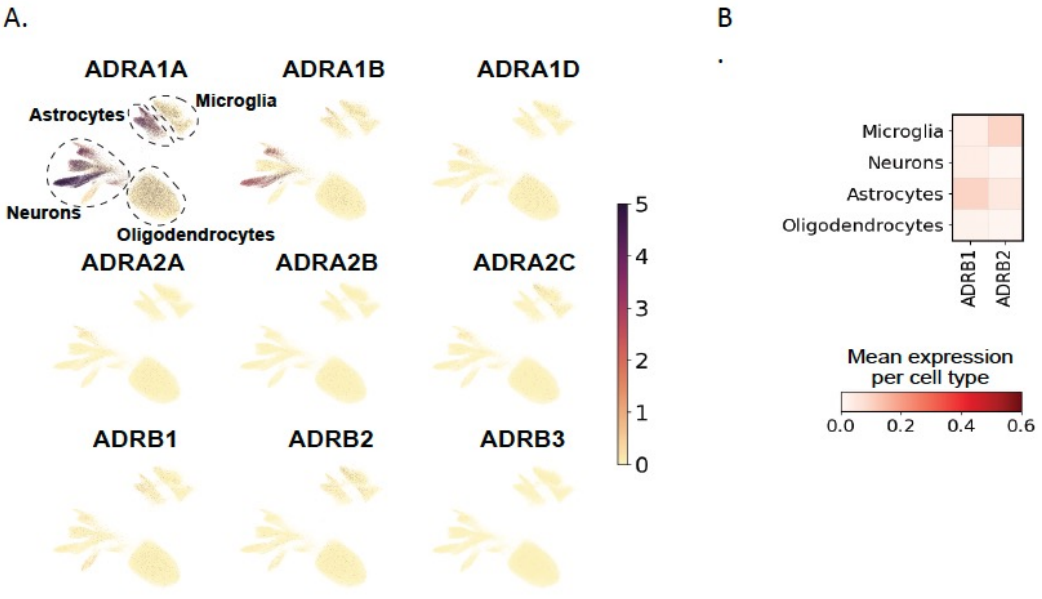


**Supplementary Figure 2. Human healthy brain tissue exhibits low expression of adrenergic receptors.** A. Feature plots of adrenergic receptor expression in healthy brain tissue. B. Heat map of β-adrenergic receptor expression in healthy brain tissue cell populations. ADRA1A= α-1A adrenergic receptor, ADRA1B= α-1B adrenergic receptor, ADRA1D= α-1D adrenergic receptor, ADRA2A= α-2A adrenergic receptor, ADRA2B= α-2B adrenergic receptor, ADRA2C= α-2C adrenergic receptor, ADRB1= β-1 adrenergic receptor, ADRB2= β-2 adrenergic receptor, ADRB3= β-3 adrenergic receptor.

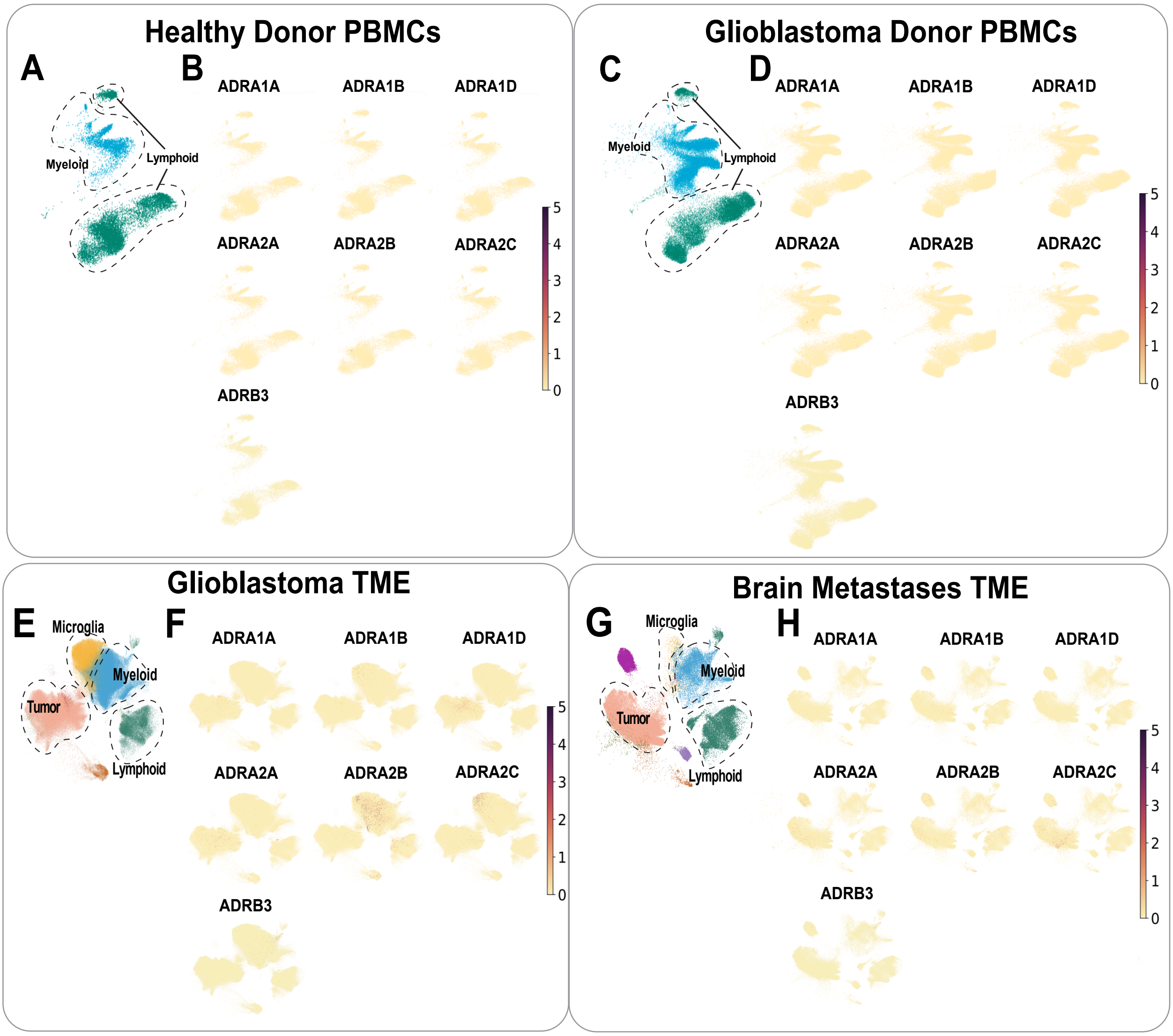


**Supplementary Figure 3. Human β3-adrenergic receptors and α-adrenergic receptors, are not highly expressed on immune cells in the periphery or tumor microenvironment.** These data were integrated from publicly available scRNAseq datasets from normal brain, glioblastoma, brain metastasis and PBMC samples. A. UMAP of clusters and cell types from healthy donor PBMCs. B. Feature plots of β3- adrenergic and α-adrenergic receptor expression in PBMCs from healthy donors. C. UMAP of cluster and cell types from glioblastoma patient PBMCs. D. Feature plots of β3–adrenergic and α-adrenergic receptor expression in PBMCs from glioblastoma patients. E. UMAP of clusters and cell types from glioblastoma tumor samples. F. Feature plots of β3- adrenergic and α-adrenergic expression in glioblastoma TME. G. UMAP of clusters and cell types from brain metastases samples. H. Feature plots β3- adrenergic and α-adrenergic receptor expression in brain metastases TME.

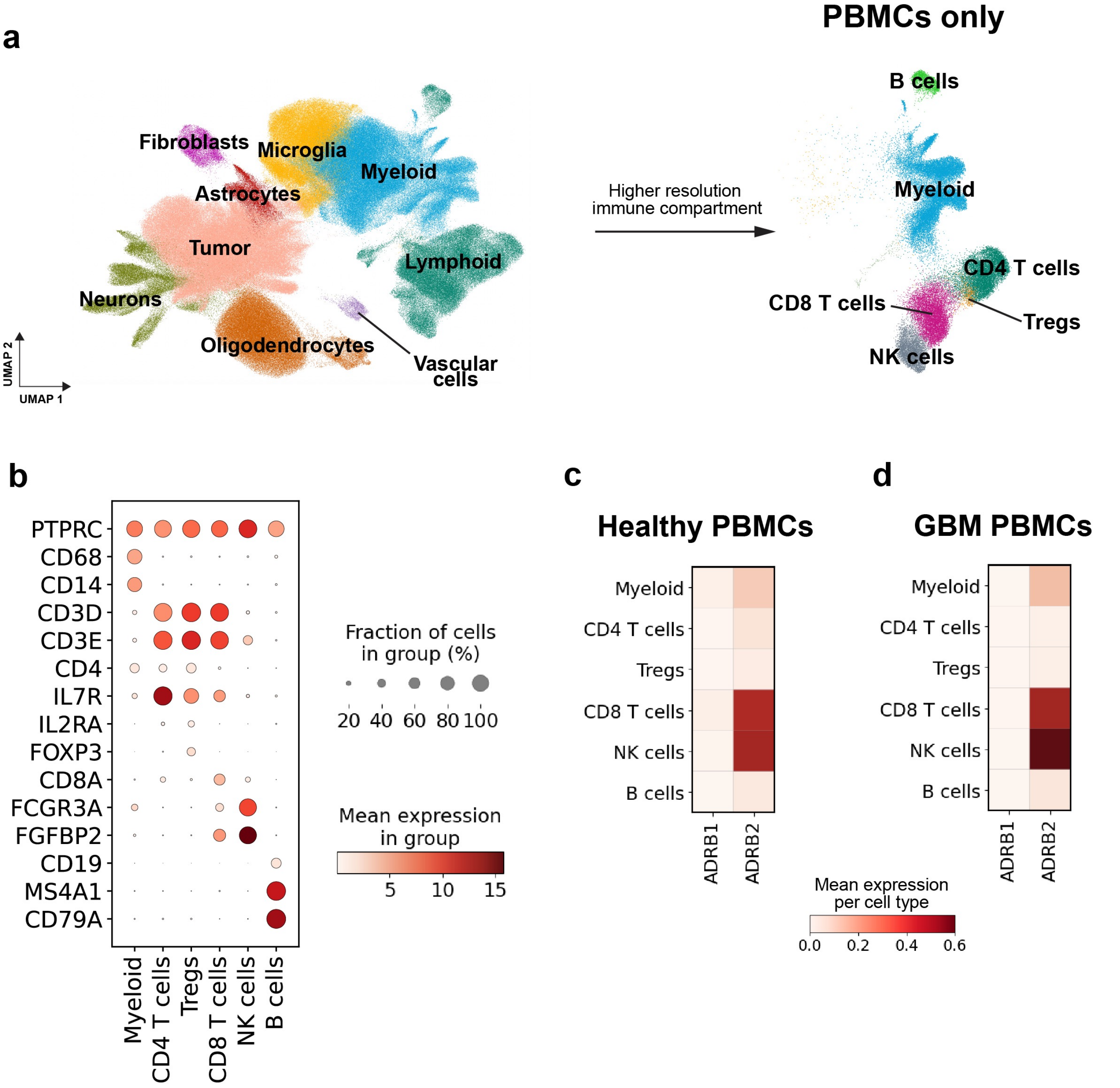


**Supplementary Figure 4. In-depth resolution of PBMC compartment from human healthy donors vs GBM patients** A. UMAP of clusters and cell types from integrated dataset showing higher resolution of the immune compartment. B. Annotation dot plot with cell cluster annotations and canonical marker genes C. $\beta 1-$ adrenergic receptor and $\beta 2-$ adrenergic receptor expression in PBMCs from healthy donors. D. $\beta 1-$ adrenergic receptor and $\beta 2-$ adrenergic receptor expression in PBMCs from GBM patients.

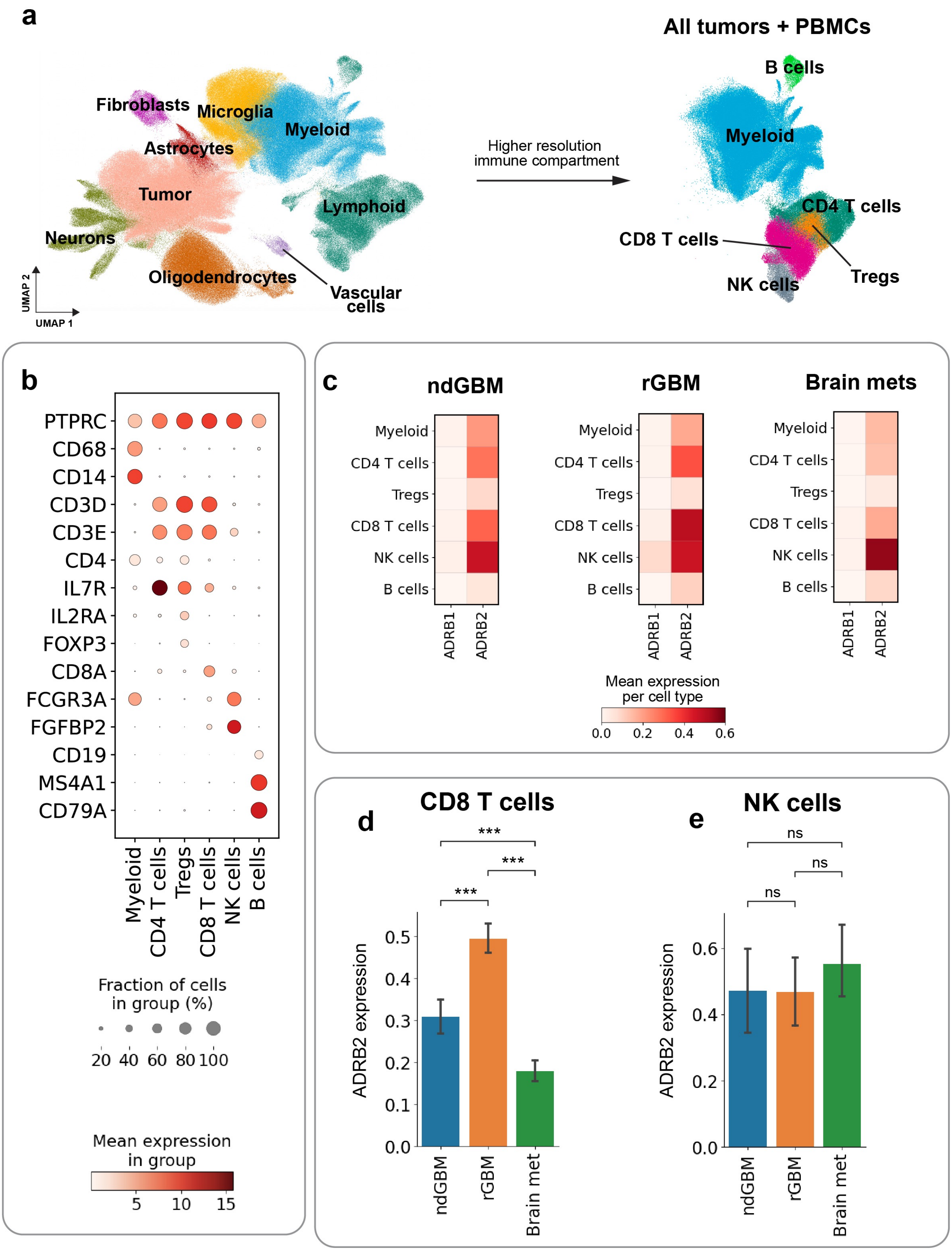


**Supplementary Figure 5. In-depth resolution of PBMC compartment from healthy healthy donors vs GBM patients** A. UMAP of clusters and cell types from integrated dataset showing higher resolution of the immune compartment. B. Annotation dot plot with cell cluster annotations and canonical marker genes C. β2– adrenergic receptor and β1– adrenergic receptor expression in immune cells from the TME of newly-diagnosed GBM patients, recurrent GBM patients or patients with brain metastasis. Comparison of β2– adrenergic receptor expression in D. CD8+ T cells and E. NK cells from newly diagnosed GBM patients, recurrent GBM patients or patients with brain metastasis. Significance was assessed using a Mann-Whitney test with Bonferonni correction; error bars represent 95% confidence intervals. *** represents p<0.001

A. B.

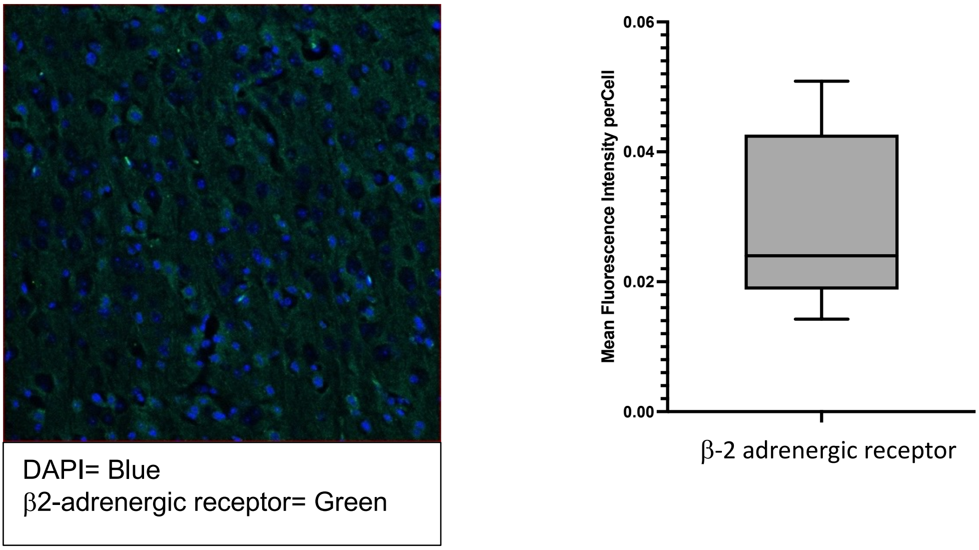


**Supplementary Figure 6. β2-adrenergic receptor is expressed within the tumor microenvironment of murine CT2A.**
A. Representative image of immunofluorescence staining for DAPI and β2-adrenergic receptor B. Quantification of the mean fluorescence intensity of β2-adrenergic receptor expression across 10 images.

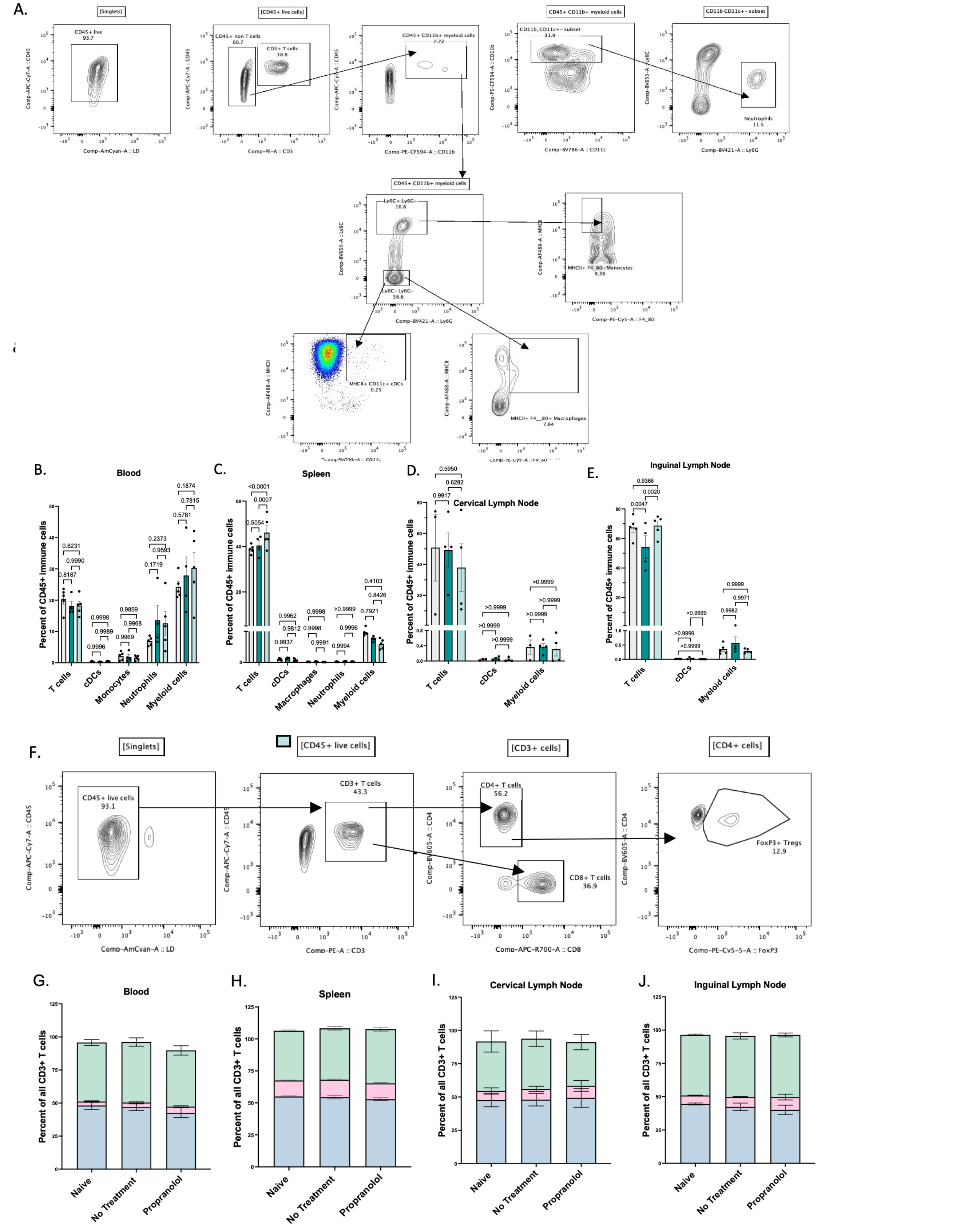

**Supplementary Figure 7. No difference in immune compartments in peripheral organs of glioma-bearing mice treated with β–blockade.** Mice harbored either no tumor or were implanted with 10K CT2A IC and given no treatment or propranolol. A. Flow cytometry gating strategy for immune profiling. Immune cell populations as a percentage of total immune cells in the B. blood C. spleen D. cervical lymph node and E. inguinal lymph node. F. Flow cytometry gating strategy for T cell subsetting. T cell subsets as a percentage of total T cells in the G. blood H. spleen I. cervical lymph node J. inguinal lymph node. Experiments were repeated independently at least 2 times. Error bars represent the standard error of the mean N=5 mice per group for all experiments. Statistical analysis was assessed using two-way ANOVAs with Tukey's multiple comparison tests.

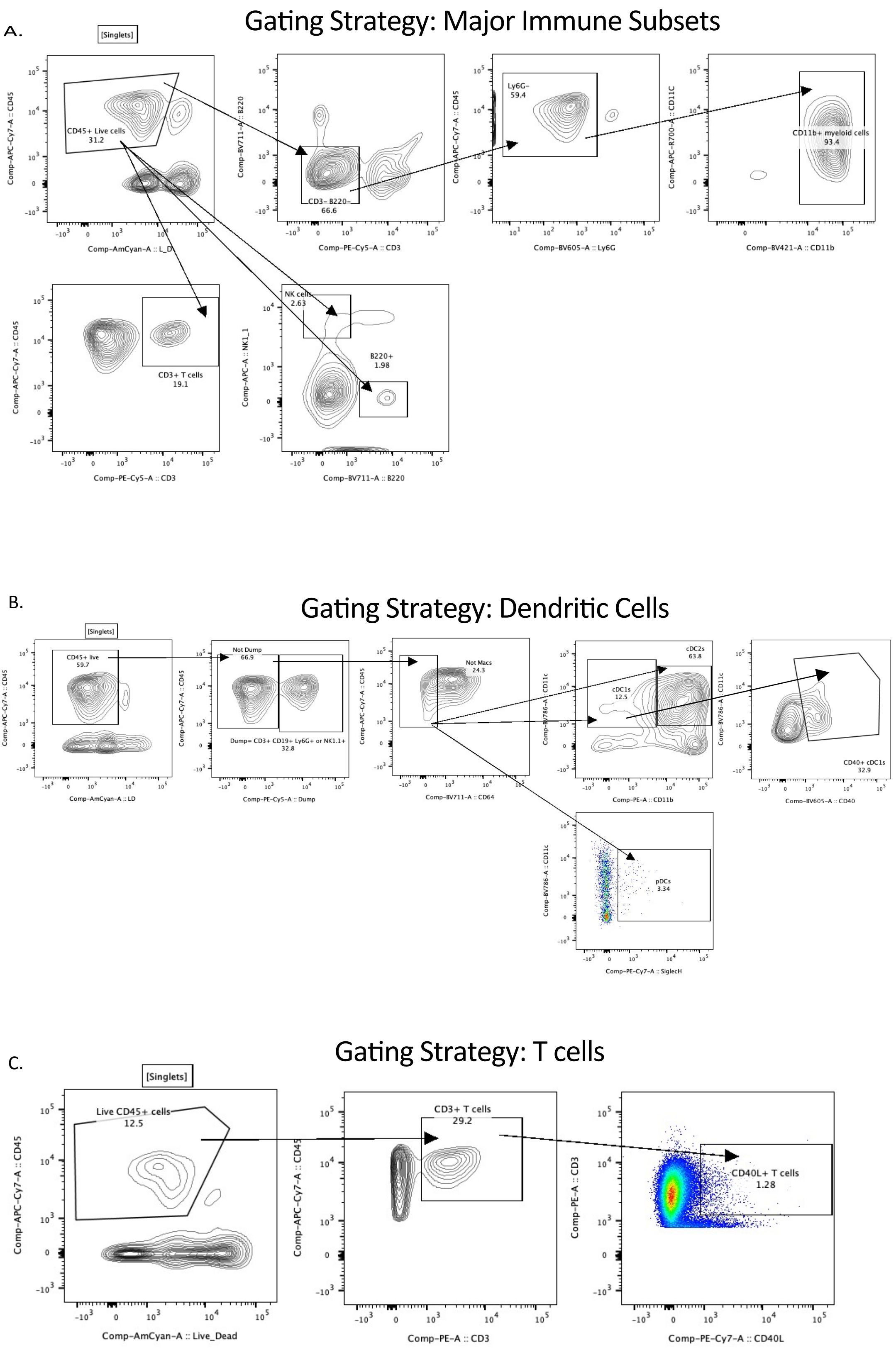


**Supplementary Figure 8. Flow cytometry gating strategies within the tumor microenvironment.** C57BL/6 mice were implanted with 10K CT2A IC and given either no treatment or propranolol. All samples were taken from the tumor and gating strategy begins after singlet gate. Gating strategy for: A. immune profiling of major subsets B. dendritic cells and C. T cells in the tumor microenvironment

A.

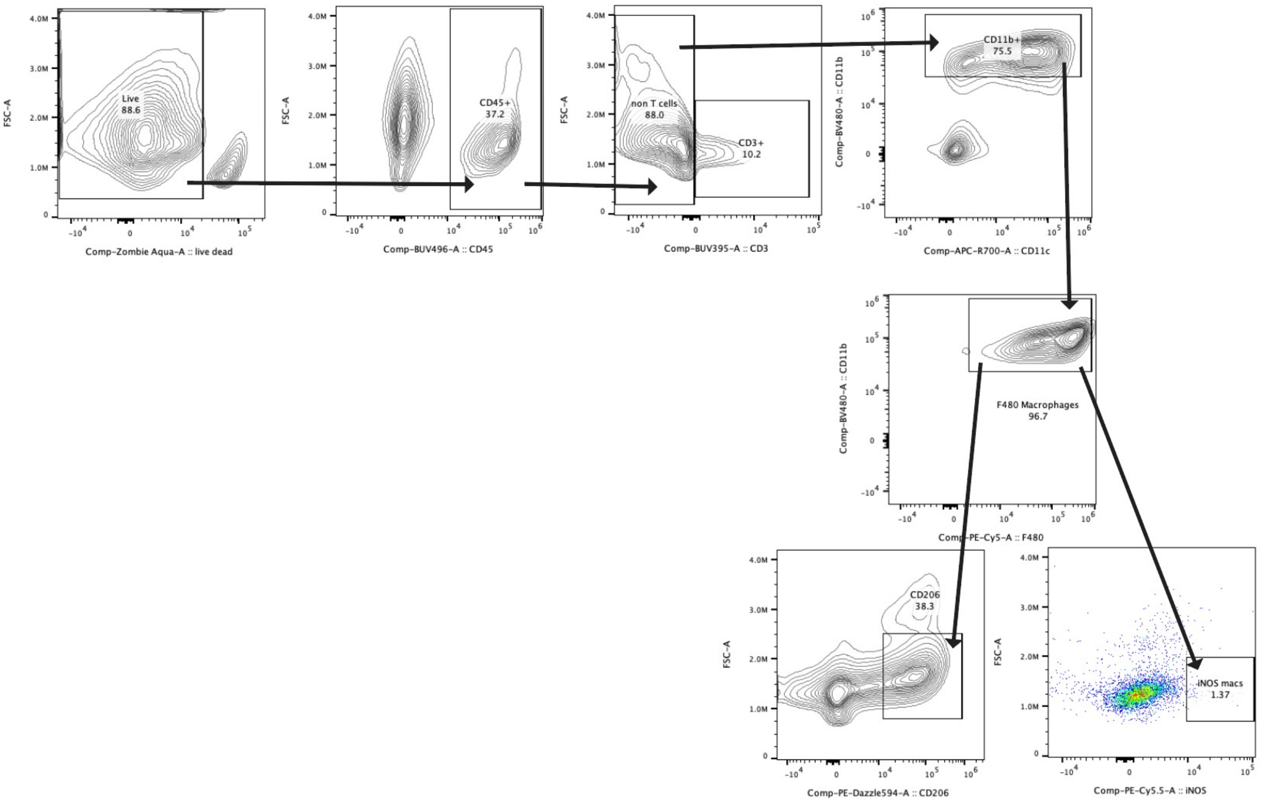


**Supplementary Figure 9. Gating strategy for macrophage phenotyping in the tumor microenvironment.** A. Flow cytometric gating strategy for macrophage phenotyping. Macrophages were defined as live, CD45+ CD3- CD11b+ F4/80+ cells and further subsetted as CD206+ “immunosuppressive” or iNOS “inflammatory” from this population.

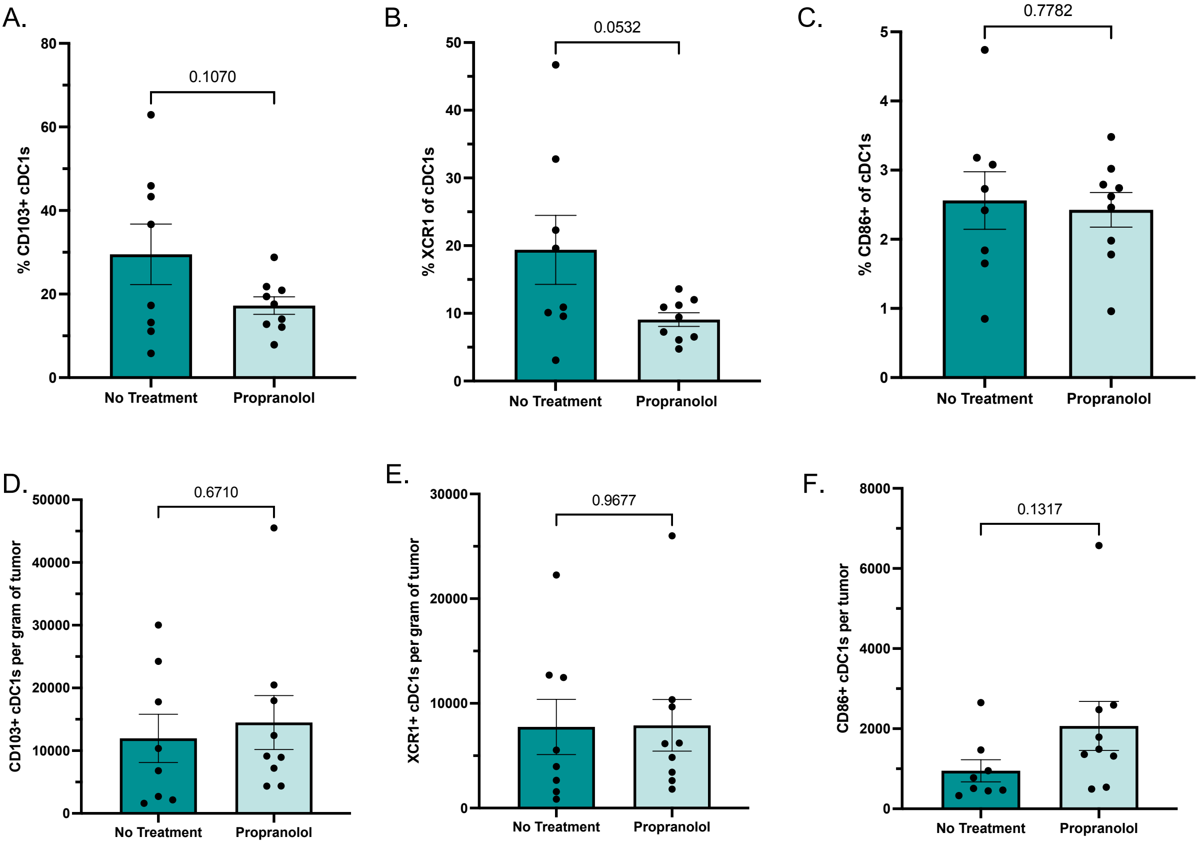


**Supplementary Figure 10. Phenotyping of conventional type 1 dendritic cells in the tumor microenvironment.** Mice were implanted intracranially with 10K CT2A and given no treatment or propranolol (n≥8 per group). A. Percent of CD103+ migratory cDC1s. B. Percent of XCR1+ cross-presenting cDC1s. C. Percent of CD86+ cDC1s. D. Number of CD103+ cDC1s per gram of tumor. E. Number of XCR1+ cDC1s per gram of tumor. Number of CD86+ cDC1s per gram of tumor. Error bars represent the standard error of the mean. Student's T. test was used for statistical comparison. Experiments were performed three times.

A.

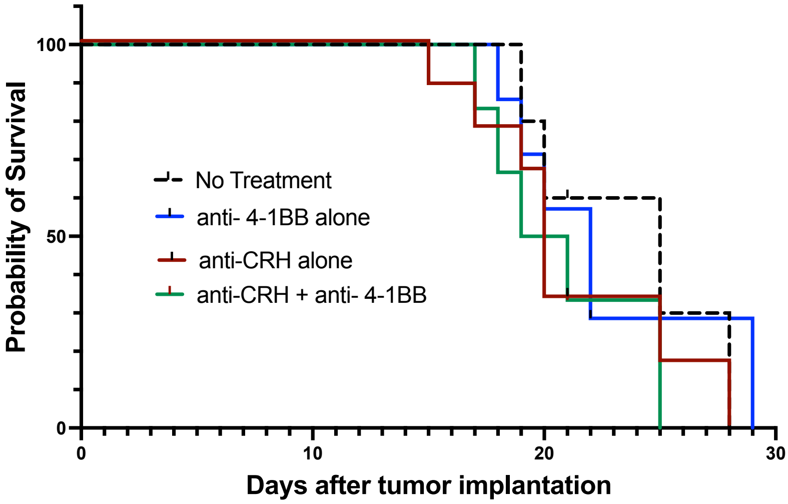


**Supplementary Figure 11. Blocking the HPA axis provides no survival benefit in combination with 4-1BB agonism**. A. Kaplan-Meier survival curve of mice with intracranial CT2A treated with: no treatment (n=5), 4-1BB agonist (n=7), anti-CRH (n=9) or anti-CRH and 4-1BB agonist (n=6). Anti-CRH is a functional inhibitor of the HPA axis ("anti-corticotropin-releasing hormone").

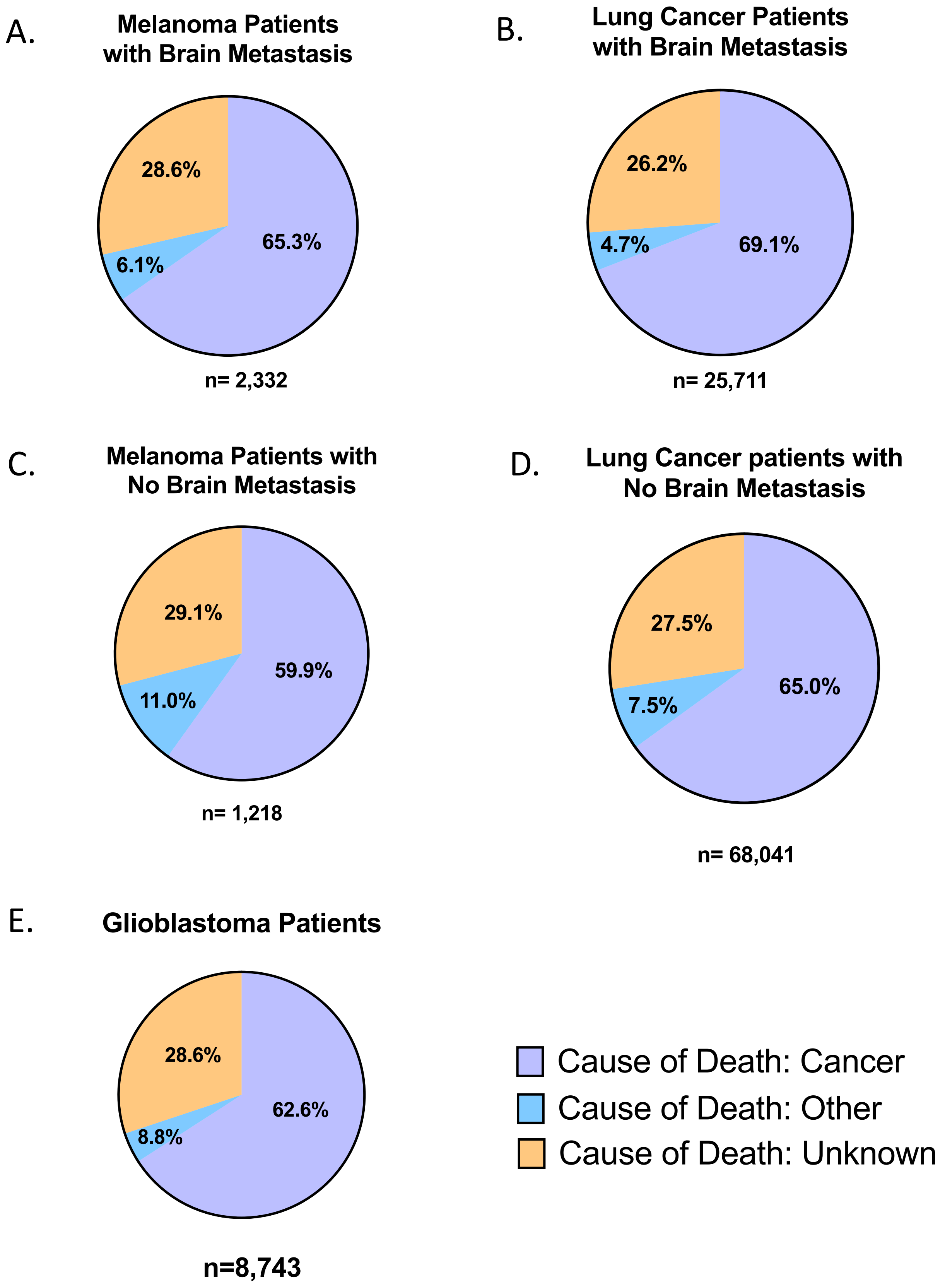


**Supplementary Figure 12. Causes of death in retrospective SEER-Medicare data.** Cause of death (cancer, other, unknown,) in: A. Metastatic melanoma patients with brain metastasis, B. Metastatic lung cancer patients with brain metastasis, C. Metastatic melanoma patients with no brain metastasis, D. Metastatic lung cancer patients with no brain metastasis, or E**.** Glioblastoma patients
