## Extended data for "Glioblastoma and other intracranial tumors elicit systemic sympathetic hyperactivity that limits immunotherapeutic responses"

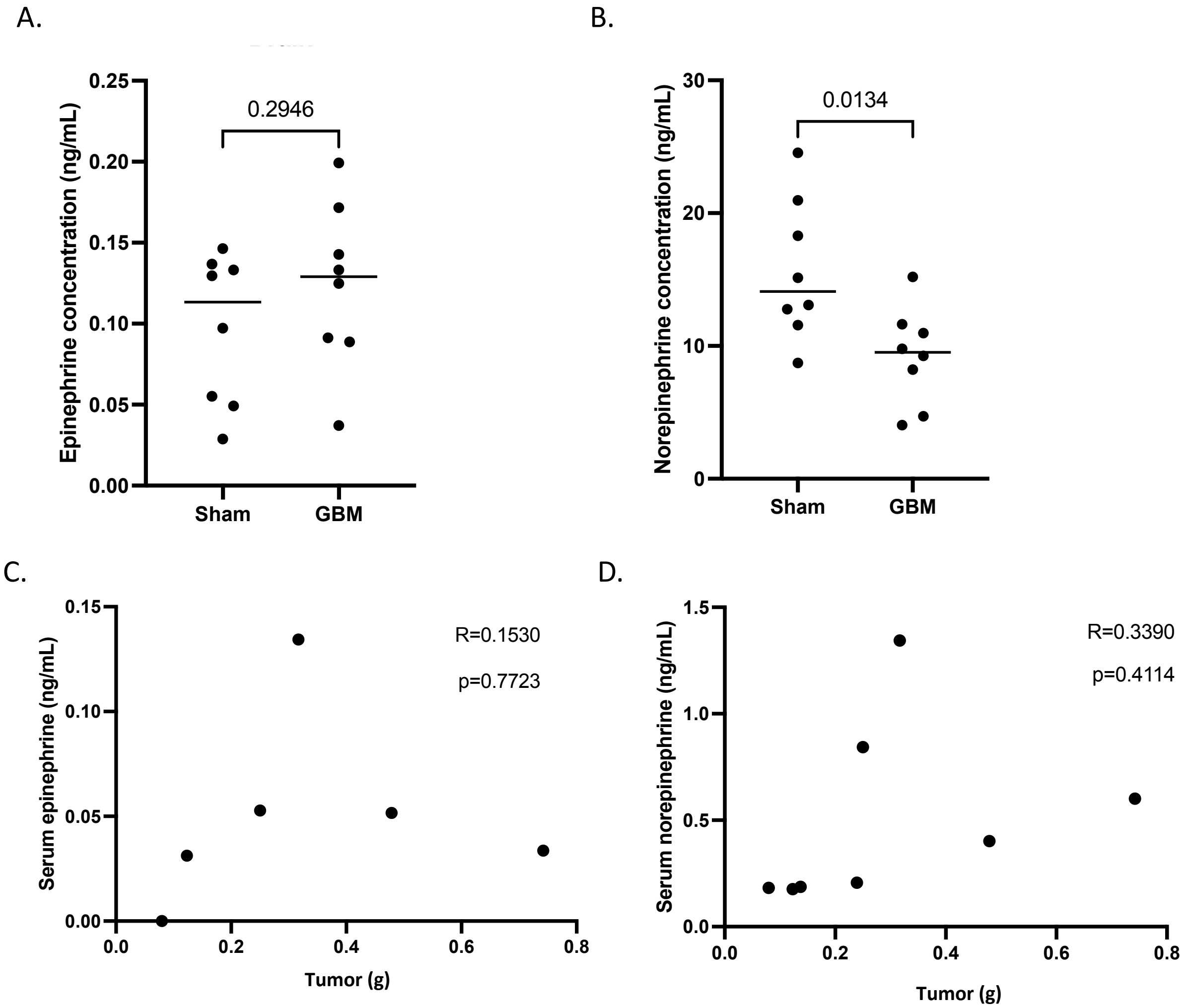


**Extended Data Figure 1. Catecholamine levels in brains of mice with or without intracranial tumors**
Mice (n=8 per group) received either intracranial injections of PBS in methylcellulose ("sham") or intracranial implantation of CT2A in methylcellulose ("GBM"). Tumor-bearing or sham-injected hemispheres were harvested from each mice upon humane endpoint and catecholamine concentrations were determined for A. epinephrine and B. norepinephrine. Statistical comparisons were assessed using unpaired T tests. C. Correlation of epinephrine concentration and weight of tumor-bearing hemisphere. D. Correlation of norepinephrine and tumor-bearing hemisphere. Epinephrine and norepinephrine concentrations were measured in tumor-bearing hemisphere.

A.

| Sample | Cells that Passed QC | Cytokines Secreted | Repeated Y/N | Cells that passed QC in repeat |
|---|---|---|---|---|
| NTB | 180 | GrzmB, IFNg, IL-17A, IL-21, IL-6, IP-10, MIP1a, RANTES, sCD137 | Yes | 649 |
| IC Tumor | 139 | GrzmB, IFNg, MIP1a IP-10, RANTES, sCD137 | Yes | 761 |
| SC Tumor | 593 | GrzmB, IFNg, IL-17A, IL-21, IL-27, IP-10, MIP1a, RANTES, sCD137 | No | NA |
| Epi/Norepi | 482 | GrzmB, IFNg, IP-10, MIP1a, RANTES, sCD137 | No | NA |
| β2-agonist | 341 | GrzmB, IFNg, IP-10, MIP1a, RANTES, sCD137 | No | NA |
| β-antagonist | 42 | GrzmB, IFNg, MIP1a IP-10, RANTES, sCD137 | Yes | 194 |

B.

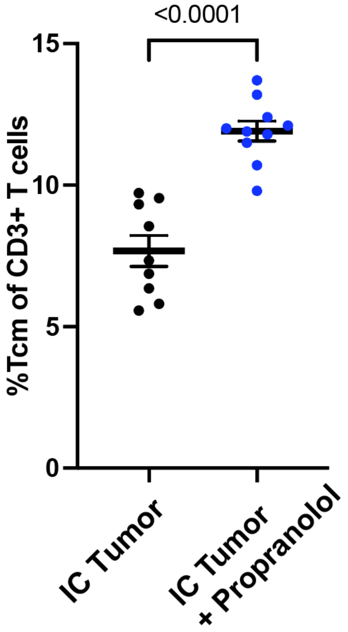


C.

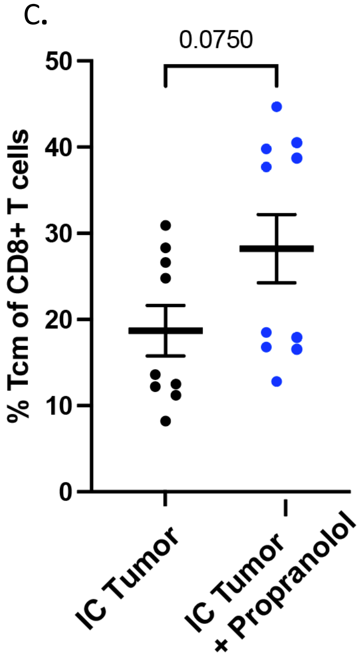


**Extended Data Figure 2. Splenic T cells have decreased polyfunctionality and proportion of central memory T cells in mice with intracranial tumors.** A. Table delineating number of cells that passed QC, which cytokines contributed to polyfunctionality and number of cells that passed QC if repeated. C. Proportion of $CD3^+$ T cells with a central memory phenotype in mice with intracranial CT2A treated with or without propranolol. C. Proportion of $CD8^+$ T cells with central memory phenotype in mice with intracranial CT2A treated with or without propranolol.

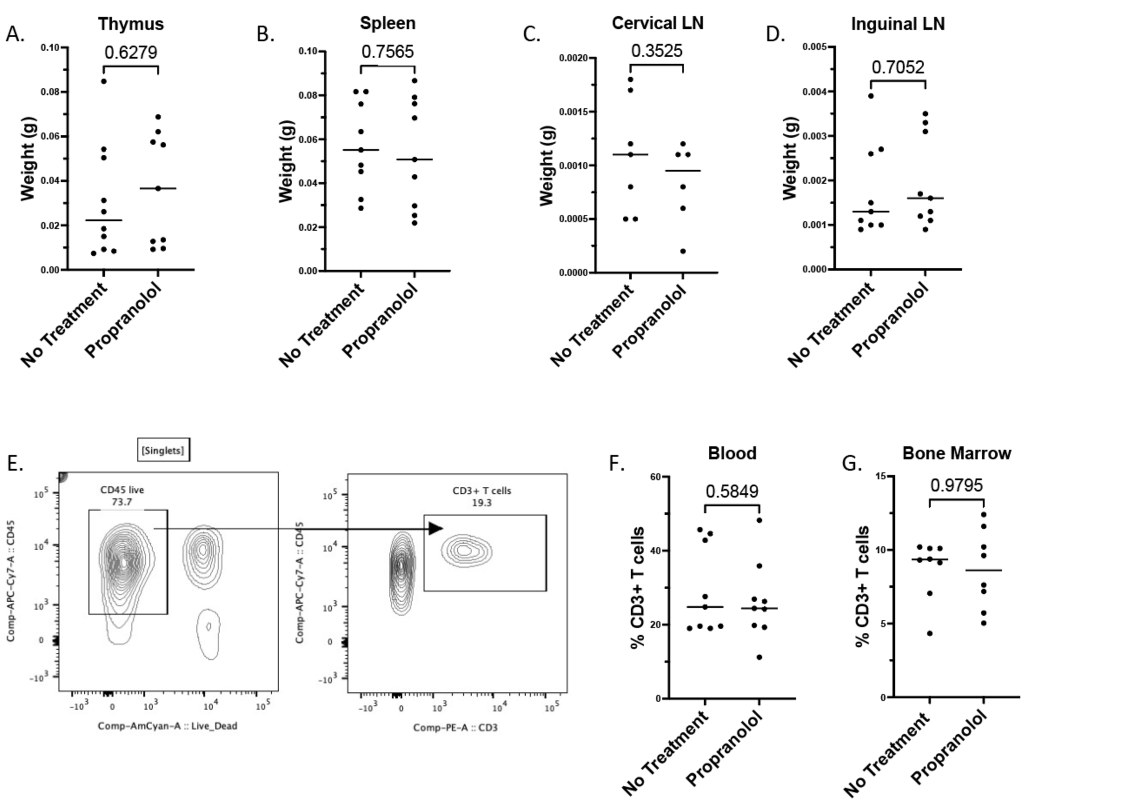


**Extended Data Figure 3. Treatment with β-blockade does not prevent systemic immune derangements in the setting of glioblastoma.** C57BL/6 mice were implanted with 10K CT2A IC given no treatment or propranolol. Organ weights of A. thymus, B. spleen, C. cervical lymph node, D. inguinal lymph node. E. Gating strategy for $CD3^+$ T cells in blood and bone marrow. F. Percent of $CD3^+$ T cells in the bone marrow. G. Percent of $CD3^+$ T cells in the blood. Statistical significance was assessed using unpaired T-tests. Experiments were repeated independently at least 2 times. N≥9 mice per group for all experiments.

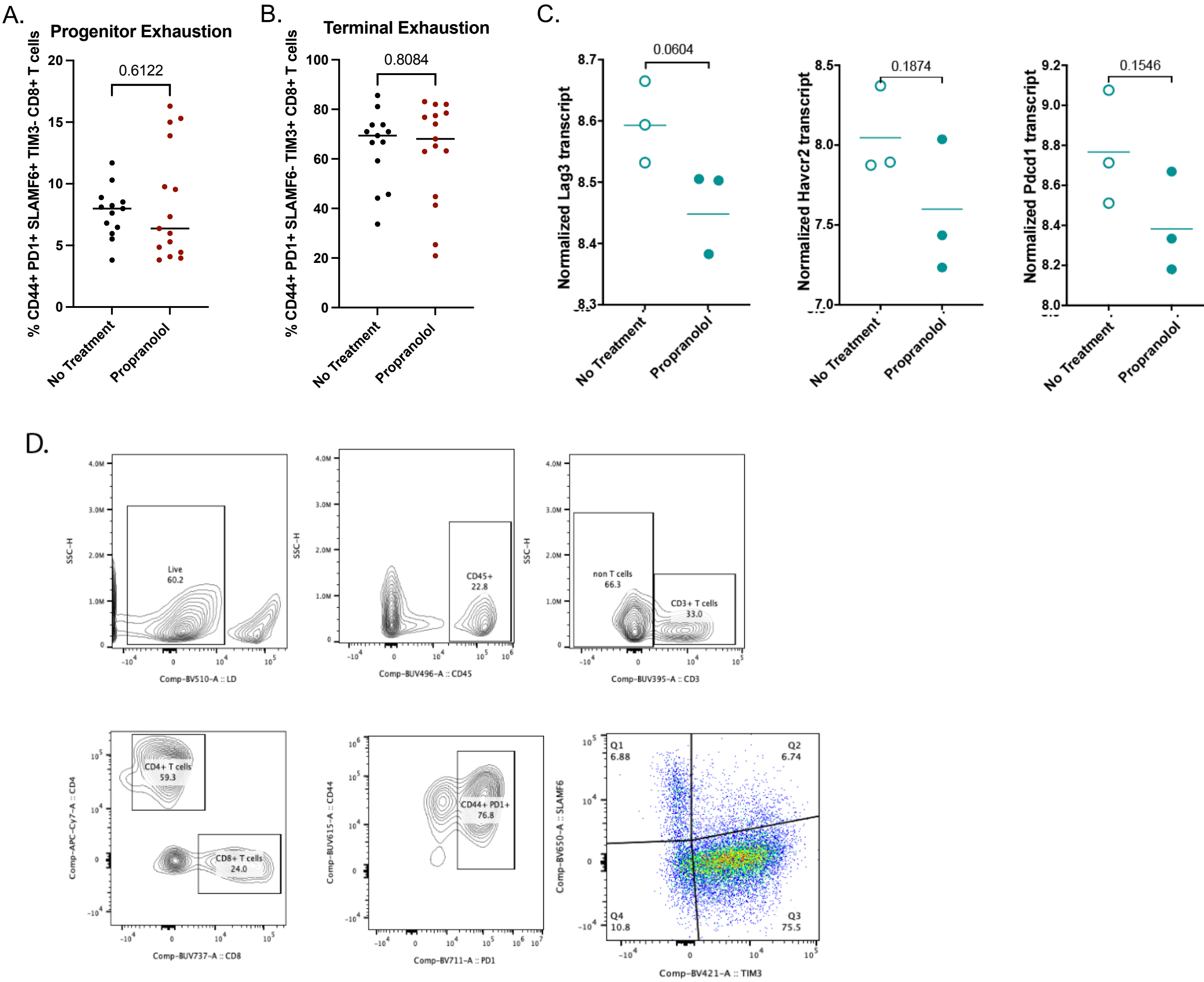


**Extended Data Figure 4. Treatment with β-adrenergic blockade does not significantly change the percentage of progenitor or terminal exhaustion.** 10K CT2A was implanted into all mice and tumors were harvested at humane endpoint. A. Percent of progenitor exhausted (SLAMF6+ TIM3-) of CD44+ PD1+ CD8+ T cells. B. Percent of terminally exhausted (SLAMF6- TIM3+) of CD44+ PD1+ CD8+ T cells. C. Transcript levels of LAG3, TIM3 and PD1. D. Gating strategy used to determine progenitor and terminal exhaustion in panels A and B. RNAseq experiments in A were performed once. Flow cytometry experiments in B and C were performed three times and represent pooled data. Significance was determined using unpaired t-tests.

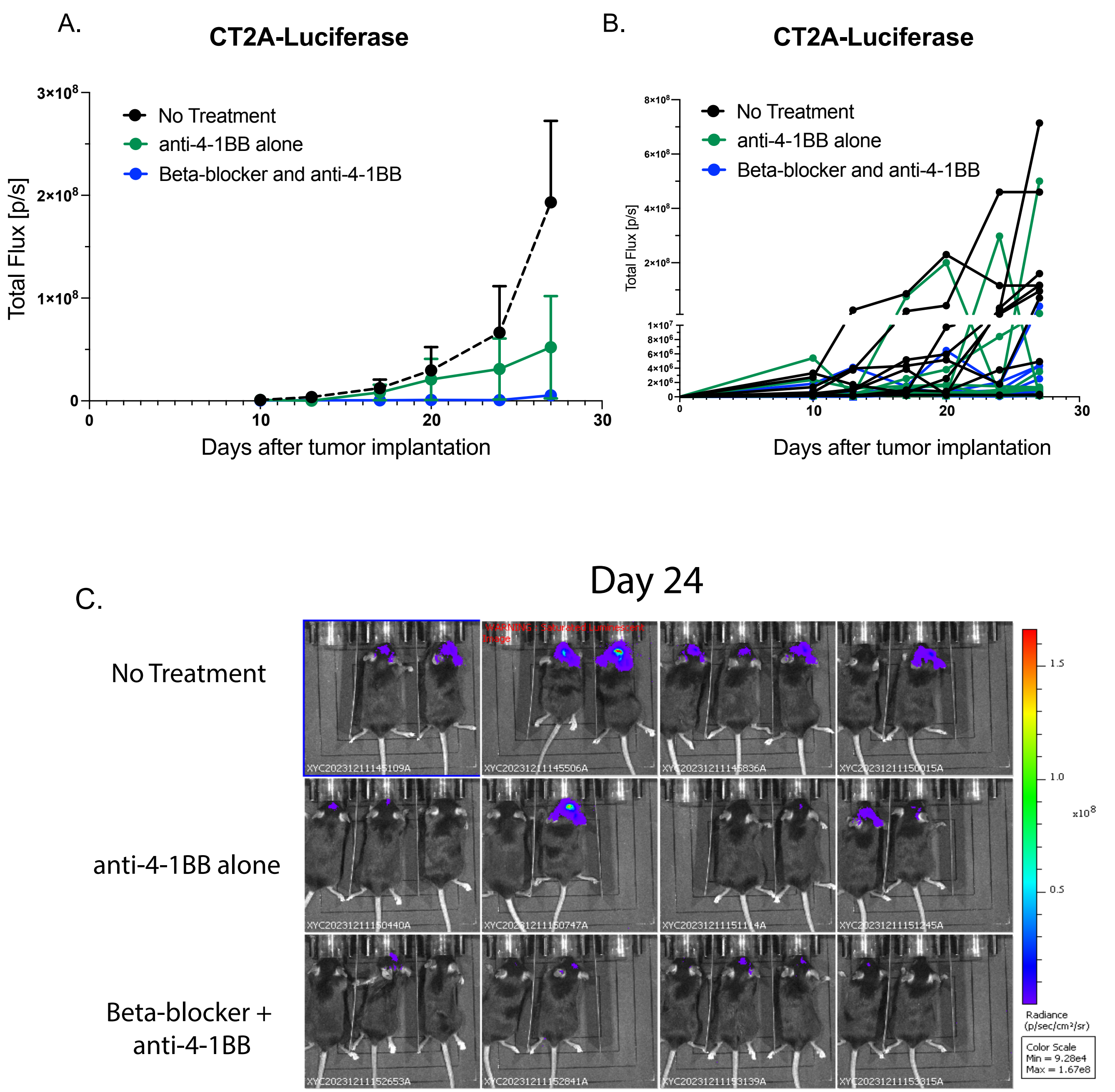


**Extended Data Figure 5. Individual plots of tumor growth from mice with intracranial CT2A-Luciferase**
All mice were implanted intracranially with 50K CT2A-luciferase and given no treatment, anti-4-1BB alone or a combination of β-blocker and 4-1BB (n=10 mice per group). A. Grouped tumor growth plot. B. Tumor growth plot of individual. C. Imaging of individual mice at Day 24 after tumor implantation .

A.

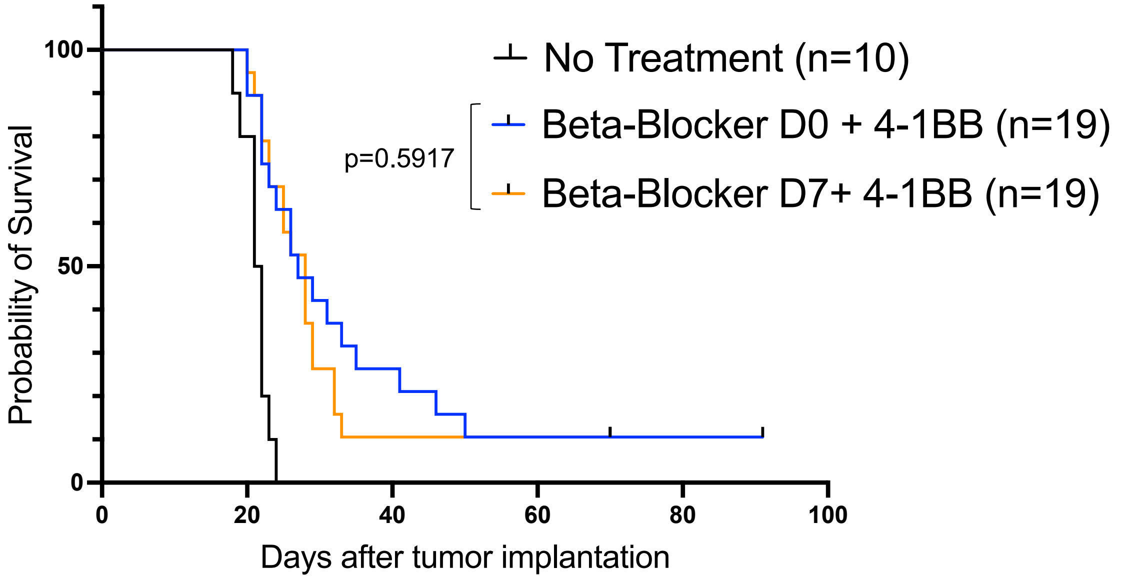


**Extended Data Figure 6. Treatment with β-blocker at day 0 or day 7 in combination with 4-1BB agonist**
All mice were implanted intracranially with 10K CT2A and were given either given no treatment, or 4-1BB agonist and β-blocker beginning at either day 0 or day 7 after tumor implantation. A log-rank test was used to determine statistical significance.
