## Supplementary Table 1 for "Glioblastoma and other intracranial tumors elicit systemic sympathetic hyperactivity that limits immunotherapeutic responses"

### Supplementary Table 1. Codes used to identify treatment patterns

| Treatment Type | Code Classification | Code |
|---|---|---|
| **Biopsy** | CPT | Neurosurgical: 61140, 61750, 61751 |
| **Diagnostic Imaging (cranial only)** | CPT | 70450-70470, 70551-70553, 78607-78608 |
| **Radiation Therapy** | ICD-9-CM | 92.23–92.24, 92.30–92.33, and 92.39. |
| | CPT | 77261-77263, 77280, 77285, 77290, 77295, 77299–77301, 77305, 77310, 77315, 77321, 77332-77334, 77336-77337, 77370-77372, 77399, 77402-77414, 77416, 77418-77420, 77425, 77427, 77430, 77432, and 0073T. |
| | HCPCS | G0173-G0174, G0242-G0243, G0251, and G0338-G0340. |
| **Stereotactic radiosurgery** | ICD-9-CM | Cranial: 92.30–92.33 and 92.39. |
| | CPT | Cranial: 61793, 61796-61800, 77371-77372, and 77432. |
| | HCPCS | Cranial: G0173, G0242, G0243, G0251, G0338, G0339, and G0340. |
| **Neurosurgical resection** | ICD-9-CM | 01.21–01.25, 01.31, 01.51, and 01.59. |
| | CPT | 61304–61305, 61312–61315, 61320–61321, 61330, 61332–61334, 61340, 61343, 61345, 61440, 61450, 61458, 61460, 61470, 61500–61501, 61510, 61512, 61514, 61516, 61518, 61519- 61522, 61524, 61526, 61530–61531, 61533–61536, 61538- 61539, 61541–61546, 61550, 61552, 61556–61559, 61563- 61564, 61570–61571, 61575–61576, 61580–61586, 61590–61592, 61596–61598, 61600–61601, 61605–61613, and 61615–61616. |
| **Chemotherapy administration services** | CPT (Intravenous) | 90782-90788, 96408-96419, 96520, 96530, 96545, 96549 |
| | CPT (Not intravenous) | 61517, 96400-96406, 96420-96425, 96440-96446, 96450, 96520-96522, 96542, 99601-99602 |
| | HCPCS Level II | Q0083-Q0085, J9000-J9999 |
| | ICD-9 CM (Diagnostic) | V58.1, V58.11, V58.12 |
| **Melanoma Specific Treatment Codes** | | |
| **Biopsy** | CPT | 11100, 11101, 11104-11107, 11300-11303, 11305-11308, 11310-11313 |
| **Excision, <3.1cm** | CPT | 11400-11403, 11420-11423, 11440-11443, 11600-11603, 11620-11623, 11640-11643 |
| **Excision, >3.0 cm** | CPT | 11404, 11406, 11424, 11426, 11444, 11446, 11604, 11606, 11624, 11626, 11644, 11646 |
| **Destruction** | CPT | 17000, 17003, 17004, 17106-17108, 17110, 17111, 17260-17264, 17266, 17270-17274, 17276, 17280-17284, 17286 |
| **Mohs** | CPT | 17311-17315 |
| **Lung Cancer Specific Treatment Codes** | | |
| **Resection <1 lobe** | CPT | 32505-32608 |
| | ICD-9-CM | 320.X, 322.X, 323.X |
| | ICD-10 | 0BBXXXX |
| **Lobectomy/bilobectomy** | CPT | 32480, 32482, 32485, 32486 |
| | ICD-9-CM | 324.X |
| | ICD-10 | 0BTCXXX, 0BTDXXX, 0BTFXXX, 0BTGXXX, 0BTHXXX, 0BTJXXX |
| **Pneumonectomy** | CPT | 32440, 32442, 32445, 32488 |
| | ICD-9-CM | 325.X |
| | ICD-10 | 0BTKXXX, 0BTLXXX, 0BTMXXX |
| **Immune checkpoint inhibitors** | | |
| **Pembrolizumab** | HCPCS | C9027, J9271 |
| | NDC | 0006-3026-01, 0006-3026-02, 0006-3026-04, 0006-3029-02, 0006-3029-01 |
| **Nivolumab** | HCPCS | C9453, J9299 |
| | NDC | 00003-3734-13, 00003-3756-14, 00003-3772-11, 00003-3774-12 |
| **Ipilimumab** | HCPCS | C9284, J9228 |
| | NDC | 00003-2327-11, 00003-2328-22 |
| **Necitumumab** | HCPCS | C9475, J9295 |
| | NDC | 00002-7716-01 |
| **Atezolizumab** | HCPCS | C9483, J9022 |
| | NDC | 50242-0917-01, 50242-0917-86, 50242-0918-01, 50242-0918-86 |
| **Ramucirumab** | HCPCS | C9025, J9308 |
| | NDC | 00002-7669-01, 00002-7678-01 |
| **Chemotherapies** | | |
| **Carboplatin** | HCPCS | J9045 |
| | NDC | 00015-3210, 00015-3211, 00015-3212, 00015-3213, 00015-3214, 00015-3215, 00015-3216, 61703-0360, 00703-3249, 47335-0150, 47335-0151, 50742-0447, 50742-0448, 66758-0047, 68083-0190, 68083-0191, 68083-0192, 68083-0193, 71288-0100, 00703-4239, 00703-4244, 00703-4246, 57277-0105, 57277-0106, 67457-0491, 67457-0492, 67457-0493, 67457-0494, 67457-0608, 55150-0386, 16729-0295, 69448-0005, 00703-4248, 61703-0339, 63323-0172, 25021-0202, 47335-0284, 47781-0603, 47781-0604, 47781-0605, 47781-0606, 50742-0445, 50742-0446, 57277-0107 |
| **Cisplatin** | HCPCS | C9418, J9060, J9062 |
| | NDC | 44567-0530, 00015-3070, 00015-3072, 00703-5747, 00703-5748, 16729-0288, 44567-0509, 44567-0510, 44567-0511, 63323-0103, 68001-0283, 68083-0162, 68083-0163, 67457-0424, 67457-0425, 70860-0206, 00069-0081, 00069-0084, 47781-0609, 47781-0610, 61126-0003, 61126-0004 |
| **Dacarbazine** | HCPCS | C9423, J9130, J9140 |
| | NDC | 00703-5075, 63323-0127, 63323-0128, 61703-0327 |
| **Docetaxel** | HCPCS | J9170, J9171 |
| | NDC | 70121-1221, 70121-1222, 70121-1223, 43066-0001, 43066-0006, 43066-0010, 00069-9141, 00069-9142, 00075-8001, 00075-8005, 00409-0366, 00409-0367, 00955-1020, 00955-1021, 00955-1022, |

| | | |
|---|---|---|
| | | 25021-0222, 43598-0258, 43598-0610, 43598-0611, 45963-0765, 63739-0932, 63739-0971, 66758-0050, 66758-0950, 25021-0245, 50742-0431, 50742-0463, 00143-9204, 00143-9205, 43598-0389, 47335-0323, 47335-0895, 47335-0939, 72485-0216, 72485-0215, 72485-0214, 71288-0143, 71288-0144, 71288-0150, 71288-0151, 00409-7870, 00409-0365, 00409-1732, 00409-4235, 00409-5068, 55150-0378, 55150-0379, 55150-0380, 68083-0401, 68083-0400, 68083-0399, 00409-0201, 00409-0368, 16729-0231, 16729-0267, 43598-0259, 45963-0734, 47335-0285, 67457-0533, 67457-0781, 69097-0372, 70700-0174, 70700-0175, 70700-0176, 00075-8003, 00075-8004, 00703-5720, 00703-5730, 16714-0465, 16714-0500, 16729-0120, 16729-0228, 39822-2120, 39822-2180, 39822-2200, 42367-0121, 45963-0781, 45963-0790, 50742-0428, 00069-9144, 00409-0369, 67457-0531, 67457-0532, 69097-0369, 69097-0371, 57884-3021 |
| **Etoposide** | HCPCS | C9414, C9425, J8560, J9181, J9182 |
| | NDC | 00378-3266, 00703-5653, 55390-0291, 55390-0292, 55390-0293, 55390-0491, 55390-0492, 55390-0493, 63323-0104, 00703-5657, 00703-5656, 16729-0114, 68001-0265, 00015-3404, 16729-0262 |
| **Gemcitabine** | HCPCS | J9201, J9198 |
| | NDC | 68001-0359, 68001-0350, 68001-0348, 68001-0342, 16714-0909, 16714-0930, 16729-0391, 16729-0419, 16729-0423, 00703-5775, 00703-5778, 00002-7502, 00781-3282, 00781-3283, 16729-0092, 16729-0117, 16729-0118, 55390-0391, 67457-0462, 67457-0463, 67457-0464, 68001-0282, 68083-0148, 68083-0149, 69097-0313, 69097-0314, 00409-0181, 00409-0182, 00409-0183, 00409-0185, 00409-0187, 25021-0209, 25021-0234, 25021-0235, 42236-0001, 42236-0002, 55111-0686, 55111-0687, 70860-0204, 70860-0205, 63323-0102, 63759-3028, 63759-3029, 71288-0117, 72485-0221, 72485-0222, 72485-0223, 00002-7501, 00143-9394, 00143-9395, 00409-0186, 23155-0213, 23155-0528, 25021-0208, 45963-0619, 63323-0125, 63323-0126, 67457-0616, 67457-0617, 67457-0618, 00069-3857, 00069-3858, 00069-3859, 00591-3562, 00591-3563, 23155-0214, 23155-0483, 23155-0484, 23155-0529, 25021-0239, 45963-0612, 45963-0620, 45963-0623, 45963-0624, 45963-0636, 47335-0153, 47335-0154, 50742-0496, 50742-0497, 50742-0498, 16729-0426, 62756-0008, 62756-0073, 62756-0102, 62756-0219, 62756-0321, 62756-0438, 62756-0533, 62756-0614, 62756-0746, 62756-0974, 71288-0113, 71288-0114, 60505-6113, 60505-6114, 60505-6115 |
| **Irinotecan** | HCPCS | C9474, J9206 |
| | NDC | 66758-0048, 68001-0284, 69171-0398, 68001-0480, 68083-0381, 70700-0169, 45963-0614, 55150-0352, 55150-0353, 55150-0354, 68001-0425, 68001-0426, 72485-0213, 68083-0382, 70700-0170, 00009-1111, 00009-7529, 00143-9583, 00143-9701, 00143-9702, 00703-4432, 00703-4434, 15054-0043, 16714-0027, 16714-0131, 25021-0214, 25021-0230, 47335-0937, 47335-0953, 50742-0401, 50742-0402, 59923-0702, 59923-0714, 59923-0715, 59923-0716, 61703-0349, 63323-0193, 72485-0211, 72485-0212, 16714-0725, 16714-0726, 23155-0179 |
| **Lomustine** | HCPCS | S0178 |
| | NDC | 00015-3030, 00015-3031, 00015-3032, 58181-3030, 58181-3031, 58181-3032, 58181-3040, 58181-3041, 58181-3042, 58181-3043 |
| **Lurbinectedin** | HCPCS | J9223 |
| | NDC | 68727-0712, |
| **Paclitaxel** | HCPCS | C9127, C9431, J9264, J9265, J9267 |
| | NDC | 47781-0595, 55390-0114, 55390-0304, 55390-0314, 66758-0043, 67457-0434, 67457-0449, 67457-0471, 68083-0178, 68083-0179, 68083-0180, 68817-0134, 70860-0200, 00703-3216, 00703-3217, 00703-3213, 00703-3218, 16714-0137, 69539-0158, 69539-0159, 69539-0157, 72205-0063, 72205-0062, 72205-0061, 00703-4764, 00703-4768, 44567-0504, 44567-0505, 44567-0506, 45963-0613, 61703-0342, 63323-0763, 00069-0076, 00069-0078, 00069-0079, 00703-4766, 00703-4767, 25021-0213, 68001-0516, 47781-0593, 47781-0594, 51991-0937, 51991-0938, 70860-0215 |
| **Pemetrexed** | HCPCS | C9213, J9304, J9305 |
| | NDC | 00002-7623, 00002-7640, 67184-0503 |
| **Temozolomide** | HCPCS | C1086, C9253, J8700, J9328 |
| | NDC | 54868-4142, 54868-5348, 54868-5350, 54868-5354, 54868-5980, 62175-0240, 62175-0241, 62175-0242, 62175-0243, 62175-0244, 62175-0245, 64144-0501, 64144-0502, 64144-0503, 64144-0504, 64144-0505, 64144-0506, 64980-0333, 64980-0334, 64980-0335, 64980-0336, 64980-0337, 64980-0338, 65162-0801, 65162-0802, 65162-0803, 65162-0804, 65162-0805, 65162-0806, 67877-0537, 67877-0538, 67877-0539, 67877-0540, 67877-0541, 67877-0542, 69189-7638, 00085-0381, 16729-0048, 16729-0050, 16729-0051, 16729-0129, 16729-0130, 40051-0604, 40051-0605, 40051-0606, 40051-0607, 40051-0608, 40051-0609, 47335-0893, 00054-0320, 00054-0321, 00054-0322, 00054-0323, 00054-0324, 00054-0325, 16729-0049, 50268-0761, 50268-0762, 47335-0890, 47335-0891, 47335-0892, 47335-0929, 47335-0930, 62559-0921, 62559-0920, 62559-0922, 62559-0923, 62559-0924, 62559-0925, 00085-3004, 00085-1366, 00085-1381, 00085-1417, 00085-1425, 00085-1430, 00085-1519, 00093-7599, 00093-7600, 00093-7601, 00093-7602, 00093-7638, 00093-7639, 00378-5260, 00378-5261, 00378-5262, 00378-5263, 00378-5264, 00378-5265, 00527-1777, 00527-1778, 00527-1779, 00527-1780, 00527-1781, 00527-1782, 00781-2691, 00781-2692, 00781-2693, 00781-2694, 00781-2695, 00781-2696, 42737-0101, 42737-0102, 42737-0103, 42737-0104, 42737-0105, 42737-0106, 43975-0252, 43975-0253, 43975-0254, 43975-0255, 43975-0257, 50268-0763, 51862-0083, 51862-0084, 51862-0085, 51862-0086, 51862-0087, 51862-0088, 75834-0132, 75834-0142, 75834-0143, 75834-0144, 75834-0145, 43975-0256, 59923-0703, 59923-0704, 59923-0705, 59923-0706, 59923-0707, 59923-0708, 59923-0709, 59923-0710, 59923-0711, 59923-0712, 59923-0713, |
| **Topotecan** | HCPCS | J8705, J9350, J9351 |
| | NDC | 66758-0051, 66435-0410, 67457-0474, 00007-4201, 00007-4205, 00007-4207, 00069-0075, 00078-0672, 00078-0673, 00078-0674, 00409-0302, 00703-4714, 16729-0151, 16729-0243, 45963-0615, 55390-0370, 62756-0023, 63323-0762, 67457-0662, 71288-0127, 25021-0206, 25021-0236, 25021-0824, 50742-0404 |
| **Vinorelbine** | HCPCS | C9440, J9390 |
| | NDC | 55390-0069, 55390-0070, 00008-0045, 64370-0532, 61703-0341, 00069-0099, 00069-0103, 00069-0205, 00703-4182, 00703-4183, 25021-0204, 45963-0607, 66758-0045, 67457-0431, 67457-0479, 67457-0481, 67457-0482, 50742-0420, 50742-0427 |
| **Betablockers** | | |

| | | |
|---|---|---|
| **Acebutolol** | NDC | 53746-670-01, 65162-669-50, 65162-670-03, 53746-670-30, 53746-669-05, 53746-670-05, 10135-630-01, 10135-631-01, 50268-050-15, 53746-669-01, 65162-669-10, 65162-670-10, 65162-670-50 |
| **Atenolol** | NDC | 0378-0757-01,0378-0757-10,60429-027-10,60429-027-90,43063-765-30,71205-096-90,64980-437-01,64980-439-10,71335-1616-1,71335-1616-2,71335-1616-3,71335-1616-4,71335-1616-5,71335-1616-6,71335-0912-2,50090-0442-1,50090-0442-4,42708-012-30,70934-142-30,61919-581-60,65841-023-02,72189-146-90,53002-2004-0,53002-2004-3,53002-2004-6,53002-2138-0,53002-2138-3,53002-2138-6,72189-135-30,67544-240-60,53002-4108-0,53002-4108-3,53002-4108-6,0378-0218-01,0378-0218-10,0378-0231-01,0378-0231-10,0615-8026-05,0615-8026-39,0615-8027-05,0615-8027-39,70518-0351-2,61919-257-30,65862-169-99,64980-438-10,67296-1589-3,63187-201-30,50090-3410-3,71335-0668-2,61919-257-90,70518-0379-0,70518-0379-2,55154-5455-0,63187-537-10,71335-1264-1,71335-1264-2,71335-1264-3,71335-1264-4,71335-1264-5,71335-1264-6,71335-1264-7,71335-1535-1,71335-1535-2,71335-1535-3,71335-1535-4,71335-1535-5,71335-1535-6,71335-1535-7,55154-5511-0,65841-022-01,65841-023-16,65841-023-24,65841-023-40,29300-410-01,29300-410-05,29300-411-01,29300-411-05,29300-412-01,29300-412-05,43063-765-90,60429-026-90,50090-5738-0,50090-5738-3,50090-5738-5,50090-5741-0,50090-2923-8,72189-152-90,67544-240-80,68071-1772-5,65841-022-24,65841-023-01,65841-024-40,70518-0882-0,65862-168-01,43063-952-30,43063-952-90,70518-2866-0,68071-2248-6,71335-0668-5,63187-201-90,68382-023-40,68382-024-01,68382-024-10,68382-024-16,50090-3410-5,68382-022-10,68382-022-40,68071-4527-1,68071-4527-3,68071-4527-6,70518-3036-1,65841-022-16,68071-4538-5,50090-3043-4,50090-3043-7,50090-3043-8,50090-3043-0,50090-3043-1,50090-3410-0,50090-3410-1,50090-3411-0,50090-3411-2,72789-144-30,72789-144-90,50090-5233-0,50090-5233-1,50090-5233-4,50090-5233-7,50090-5233-8,68071-4527-9,68071-4274-1,68071-4274-3,68071-4274-6,68071-4274-8,68071-4274-9,71205-096-30,71205-096-60,68382-022-16,68382-022-24,68382-023-01,50090-5266-3,50090-5266-5,50090-5363-7,50090-5363-8,50090-5367-8,50090-5408-2,50090-5408-5,42708-143-30,71335-0912-5,50090-5266-0,68071-4538-9,68071-4751-3,67296-1270-3,68071-4766-9,60760-164-90,63187-092-30,43353-834-45,43353-834-73,71335-0954-1,71335-0954-2,71335-0954-3,71335-0954-4,71335-0954-5,71335-0954-6,71335-0954-7,67544-332-60,0615-8372-05,0615-8372-30,0615-8372-39,0615-8373-05,0615-8373-30,0615-8373-39,55700-754-30,55700-754-90,55700-755-30,70518-0379-1,43353-834-80,67544-161-30,67544-161-80,67544-240-45,67296-1201-9,63187-537-90,68071-4526-3,68071-4526-5,68071-4526-6,68071-4526-8,68071-4526-9,68788-7876-1,68788-7876-3,68788-7876-6,68788-7876-9,63187-537-60,68071-4553-1,68071-5033-1,71335-0988-1,71335-0988-2,71335-0988-3,71335-0988-4,71335-0988-5,71335-0988-6,43063-805-30,68071-4538-8,71335-0668-1,65841-023-10,65841-024-01,65841-024-10,60760-256-90,60760-437-90,60760-643-30,60760-643-90,70518-3036-0,61919-279-30,61919-664-30,61919-854-30,61919-854-90,65862-168-99,65862-170-99,71335-0912-1,50090-0442-8,43353-986-60,60760-787-90,60429-025-10,50090-5741-1,50090-5741-4,63629-1785-5,63629-1785-6,50090-5741-7,50090-5741-8,63629-8029-1,63629-8029-2,50090-0442-7,71335-0668-3,71335-0668-4,60429-025-90,43353-834-30,50090-2923-7,63629-1785-1,63629-1785-2,63629-1785-3,63629-1785-4,68071-1771-1,70518-0399-0,63629-2626-1,63629-2626-2,63629-2626-3,63629-2626-4,63629-2626-5,70518-3007-0,70518-3007-1,50090-5408-0,67296-1726-3,67296-1732-3,63187-710-30,63187-710-60,70518-2939-2,68382-023-10,63187-201-60,51079-685-20,71335-0912-3,68382-024-40,65841-024-16,68071-5162-1,71335-0912-6,65862-169-01,60429-026-10,71335-0912-4,0904-7187-61,43063-805-01,43063-805-90,68645-585-54,60760-164-30,51079-759-20,68788-7528-1,68788-7528-3,68788-7528-6,68071-4538-3,68788-7528-9,68788-7535-1,68788-7535-3,68788-7535-6,68788-7535-9,68788-7616-1,68788-7616-3,68788-7616-6,68788-7616-9,70934-177-30,63187-537-30,50090-5363-0,50090-5367-0,68788-7784-1,68788-7784-3,68788-7784-6,68788-7784-9,68071-4538-6,72189-135-90,71205-096-10,67296-1201-3,67296-1288-3,68382-023-02,60687-605-01,51079-684-20,50090-2923-0,50090-3411-5,53002-1004-0,53002-1004-3,53002-1004-6,70518-3036-2,67544-161-45,67544-240-30,70518-2939-0,70518-2939-1,60760-642-90,53002-1108-0,53002-1108-1,53002-1108-3,53002-1108-6,53002-1138-0,53002-1138-3,64980-439-01,65862-170-01,70518-0399-2,43353-834-60,70518-0882-2,63187-092-90,70934-177-60,70934-177-90,70934-219-30,70934-219-90,70934-457-60,70934-458-30,70934-463-30,60760-163-90,70934-537-30,70934-537-90,67544-161-92,68382-022-01,68382-023-16,68382-023-24,63187-710-90,71205-229-30,71205-229-60,71205-229-90,67544-161-60,71205-496-30,71205-496-60,71205-496-90,71335-0540-1,71335-0540-2,71335-0540-3,71335-0540-4,71335-0540-5,50090-3411-4,71335-1108-1,71335-1108-2,71335-1108-3,71335-1108-4,71335-1108-5,71335-1108-6,50090-0442-0,0093-0752-01,0093-0752-10,0093-0753-01,0093-0753-05,0093-0787-01,0093-0787-10,65841-022-10,65841-022-40,61919-914-71,67296-1558-3,64980-437-10,64980-438-01,67296-1270-6,72189-146-30,72189-146-60,70954-390-10,70954-391-10,10135-733-01,10135-734-01,55289-993-90,16714-936-01,16714-937-01,68071-3065-3,68071-3065-6,68071-3065-9,29300-400-01,29300-400-05,29300-401-01,29300-401-05,0591-5782-01,0591-5783-01,55289-993-30,70518-2446-0,70710-1167-1,70710-1168-1,70771-1372-1,70771-1373-1,55289-993-60,52427-383-90,52427-382-90,52427-430-90,52427-431-90,52427-429-90 |
| **Betaxolol** | NDC | 60429-753-01,24658-700-01,24658-701-01,42806-038-01,17478-705-10,17478-705-11,17478-705-12,17478-705-25,42806-039-01,42806-039-10,42806-038-10,61314-245-01,61314-245-02,61314-245-03,10702-013-01,10702-014-01,0065-0246-10,0065-0246-15 |
| **Bisoprolol** | NDC | 71335-1678-1,71335-1678-2,71335-1678-3,71335-1678-4,71335-1678-5,52817-270-10,52817-270-30,52817-271-10,52817-271-30,68071-5097-3,63629-6907-1,63629-6907-2,63629-6907-3,63187-871-90,63187-871-60,68071-1582-3,65862-086-01,65862-086-30,51407-645-01,51407-645-30,51407-646-01,51407-646-30,61919-787-90,70954-455-10,62332-603-30,62332-603-31,62332-603-71,62332-604-30,62332-604-31,50268-127-15,63629-5173-1,63629-5173-2,63629-5173-3,63629-5173-4,63629-5173-5,70954-455-20,70954-456-10,70954-456-20,63187-871-30,29300-126-01,29300-126-05,29300-126-13,29300-127-01,29300-127-05,29300-127-13,65862-087-01,65862-087-30,62332-604-71,0093-3241-01,0093-3242-01,0093-3243-56,68462-878-01,68462-878-05,68462-878-30,68462-879-01,68462-879-05,68462-879-30,68462-880-01,68462-880-05,68462-880-30,70518-3333-0,42799-920-01,42799-920-02,42799-920-30,42799-921-01,42799-921-02,42799-921-30,42799-922-01,42799-922-02,42799-922-30,71335-1161-1,71335-1161-2,71335-1161-3,71335-0595-1,71335-0595-2,71335-0595-3,70954-412-10,70954-412-30,70954-412-40,70954-413-10,70954-413-30,70954-413-40,70954-414-10,70954-414-30,70954-414-40,50090-0796-0,50090-5905-0,0378-0501-01,0378-0503-01,0378-0505-01,71335-0240-1,71335-0240-2,70934-705-90,70518-1947-1,0378-0501-05,0378-0503-05,0378- |

| | | |
|---|---|---|
| | | 0505-05,29300-187-01,29300-187-05,29300-187-10,29300-187-13,29300-188-01,29300-188-05,29300-188-10,29300-188-13,29300-189-01,29300-189-05,29300-189-10,29300-189-13,50090-5849-0,50090-5849-2,67296-1268-3,70518-3207-0,71335-1769-1,71335-1769-2,71335-1769-3,71335-1931-1,71335-1931-2,71335-1931-3,50090-0789-0,50090-0789-1,50090-0789-2,50090-0799-0,67296-0947-1,70518-1947-0,51285-047-02,51285-050-02,51285-040-01 |
| **Carteolol** | NDC | 61314-238-05,61314-238-10,61314-238-15,58768-001-01,58768-001-02,58768-001-04 |
| **Carvedilol** | NDC | 68788-8151-1,68788-8151-3,68788-8151-6,68788-8151-9,51407-040-01,68788-9265-6,55111-252-05,55111-253-01,55111-253-10,55111-255-10,71335-1623-1,71335-1623-2,71335-1623-3,71335-1623-4,71335-1623-5,71335-1623-6,68788-9789-3,68382-092-01,50090-1856-0,50090-1856-1,50090-1856-2,50090-1856-3,65841-618-05,65841-619-01,55111-254-10,55111-254-30,71335-1813-7,61919-533-30,61919-533-90,63187-131-90,71335-0034-2,68788-8177-3,68788-8177-6,68788-8177-9,0615-8018-30,0615-8018-39,42708-072-60,65841-616-01,65841-616-05,65841-616-17,65841-617-17,70518-0426-1,65862-143-99,68462-162-01,68462-162-18,68462-162-60,68462-163-05,68462-163-60,68462-165-05,70518-0356-1,70934-910-60,70934-910-90,68382-092-05,68382-093-17,63187-409-60,63187-447-60,63187-424-60,63187-941-60,63187-941-90,51655-033-26,68788-7910-1,68788-7910-3,68788-7910-6,68788-7910-9,71335-0034-3,71335-0034-6,68001-152-00,68001-154-00,60760-235-60,43063-126-60,76385-112-50,65841-618-01,68071-2633-3,68071-2633-6,68071-2633-8,68071-2633-9,65862-142-01,65862-145-05,65862-145-99,68071-3078-3,68071-3078-6,68071-3078-9,68071-2230-8,68462-162-10,68462-165-10,63187-409-90,68071-3151-3,68071-3151-6,68071-3151-9,68071-5275-8,68071-3269-3,68071-3269-6,68071-3269-8,68071-3269-9,43063-129-60,70518-0426-0,70518-0426-3,65862-142-99,65862-144-05,71610-062-60,71610-063-53,50090-2119-0,50090-2119-3,50090-2119-4,65862-143-01,65862-143-05,51079-771-20,55154-5677-0,71335-1457-1,71335-1457-2,71335-1457-3,71335-1457-4,71335-1457-5,71335-1457-6,71335-1457-7,71335-1457-8,61919-984-30,68788-7883-1,68788-7883-3,68788-7883-6,68788-7883-8,68788-7883-9,50090-4171-0,50090-4171-1,50090-4171-4,68382-093-01,68382-093-05,68382-095-05,65841-617-01,65841-619-05,68645-350-59,50090-4898-0,68462-162-05,68462-164-01,68462-164-18,55111-252-30,55111-253-05,71335-2023-1,71610-062-80,71335-2023-2,71335-2023-3,71335-2023-4,71335-2023-5,71335-2023-6,71335-2023-7,71335-2026-1,71335-2026-2,71335-2026-3,71335-2026-4,71335-2026-5,71335-2026-6,71335-2026-7,71335-2033-1,71335-2033-2,71335-2033-3,71335-2033-4,71335-2033-5,71335-2033-6,71335-2033-7,71335-2033-8,71335-0368-1,71335-0368-2,71335-0368-3,71335-0368-4,71335-0368-5,71335-0368-6,71335-0368-7,76385-110-50,68382-094-01,50090-5115-1,50090-2066-0,50090-2066-1,50090-2066-2,50090-2066-3,60760-234-60,76385-113-01,51655-397-26,63187-946-30,51079-931-20,70518-1377-2,70518-1377-3,76385-113-50,0615-7944-30,0615-7944-39,0615-7945-30,0615-7945-39,0615-7946-30,0615-7946-39,71335-0561-1,71335-0561-2,71335-0561-3,71335-0561-4,71335-0561-5,71335-0561-6,71335-0561-7,71335-0561-8,71610-063-30,50090-5115-0,55111-252-01,51407-039-05,51407-041-01,68462-163-01,68462-164-10,68462-165-01,68462-165-60,68071-4495-3,68071-4495-6,68071-4495-8,68071-4495-9,68462-163-18,68462-164-05,68462-165-18,68001-151-03,68001-153-03,43353-837-80,43353-838-30,55154-7278-0,71335-0034-1,0904-6300-61,0904-6301-61,0904-6302-61,0904-6303-61,0378-3631-01,0378-3631-05,0378-3632-01,0378-3632-05,0378-3633-01,0378-3633-05,0378-3634-01,0378-3634-05,50090-2175-0,50090-2175-1,76385-111-01,55700-819-18,55700-819-90,43063-833-60,65862-142-05,65862-144-99,65862-145-01,68071-4956-6,68071-4974-3,63187-570-90,68071-4980-6,68788-9789-6,58118-0163-8,0781-5221-01,0781-5222-01,0781-5223-01,0781-5224-01,68645-351-59,72888-034-00,72888-034-01,68071-4600-3,68071-4600-6,68071-4600-8,68071-4600-9,50090-1064-0,50090-1064-1,50090-1064-4,60760-376-60,60760-494-60,60760-532-60,68645-496-59,72888-034-05,72888-034-30,72888-035-00,72888-035-01,72888-035-05,72888-035-30,72888-036-00,72888-036-01,72888-036-05,72888-036-30,72888-037-00,72888-037-01,72888-037-05,50090-1069-0,50090-1069-1,61919-219-30,61919-219-90,50090-1069-3,50090-1069-4,61919-728-30,61919-728-90,61919-730-30,61919-730-60,61919-730-90,71335-0034-4,71335-0034-5,65862-144-01,61919-984-90,63187-447-90,68788-9265-9,50090-2119-1,70786-0144-1,0615-8387-39,0615-8388-30,0615-8388-39,0615-8389-30,0615-8389-39,0615-8390-30,0615-8390-39,55154-6884-0,55154-6885-0,72888-037-30,63187-409-30,68788-9265-1,51407-039-01,51407-041-05,51407-042-05,55111-252-10,55111-253-30,55111-253-78,55111-254-01,55111-255-78,71610-062-53,71610-063-60,63187-570-30,50090-2175-2,50090-2175-3,50090-2175-4,50090-2555-0,50090-2555-1,50090-2555-3,63629-4060-1,63629-4060-2,63629-4060-3,63629-4060-4,63629-4060-5,63629-4060-6,63629-4070-1,63629-4070-2,63629-4070-3,63629-4070-4,63629-4070-5,63629-4070-6,43353-837-60,43063-125-93,43063-126-93,71610-066-80,71610-067-80,71610-063-80,68084-854-01,43353-838-53,43353-874-30,68001-151-00,68001-152-03,68001-154-03,68071-4470-3,68071-4470-6,68071-4470-9,70518-0356-0,43353-832-09,65841-617-05,65841-618-17,76385-110-01,76385-112-01,63187-946-90,63187-570-60,63187-946-60,68382-094-05,0615-8387-30,55154-6883-0,50090-2555-4,55111-252-78,55111-255-30,71610-067-60,71610-132-60,68788-7458-1,68788-7458-3,68788-7458-6,68788-7458-8,68788-7458-9,51079-930-20,68788-7539-1,68788-7539-3,68788-7539-6,68788-7539-9,43353-832-80,43353-838-60,43353-838-80,43353-874-60,43353-874-80,68788-7603-1,68788-7603-3,68788-7603-6,68788-7603-8,68788-7603-9,63187-941-30,68788-9789-1,68788-9789-9,70518-1377-0,55154-5675-0,71335-1813-1,76385-111-50,71335-1813-2,71335-1813-3,71335-1813-4,71335-1813-5,71335-1813-6,55111-254-05,55111-255-01,51407-042-01,70934-653-30,70934-653-90,65841-619-17,0093-0051-01,0093-0051-05,0093-0135-01,0093-0135-05,68382-092-17,68382-095-01,70518-0426-4,70518-0582-1,70518-1377-1,70518-1826-0,70518-1826-1,70518-1826-2,60760-233-60,63187-131-30,63187-131-60,55111-254-78,55111-255-05,70518-2389-0,68462-163-10,51079-932-20,55154-5676-0,55154-5678-0,43353-832-60,51407-040-05,70934-211-60,70934-406-30,68462-164-60,70934-515-30,70934-515-60,70934-515-90,70934-516-30,70934-516-60,70934-517-30,70934-517-60,70934-609-30,70934-609-60,70934-609-96,70934-686-30,70934-686-60,70934-686-90,70934-686-96,70934-707-60,70934-707-90,70934-707-96,70934-753-96,70934-759-60,70934-759-90,68382-094-17,68382-095-17,53002-1556-0,53002-1556-3,53002-1572-0,53002-1572-3,53002-1573-0,53002-1573-3,53002-1574-0,53002-1574-3,43063-129-30,43063-129-93,50090-4171-3,70934-707-30,71335-1937-1,71335-1937-2,71335-1937-3,71335-1937-4,71335-1937-5,71335-1937-6,71335-1937-7,63187-447-30,71335-1128-1,71335-1128-2,71335-1128-3,71335-1128-4,71335-1128-5,71335-1128-6,68001-153-00,71610-132-53,35356-526-90,0093-7295-01,0093-7295-05,0093-7296-01,0093-7296-05,68788-9265-3,71610-066-60,71610-132-80,71335-1407-1,71335-1407-2,71335-1407-3,71335-1407-4,71335-1407-5,71335-1407-6,43353-837- |

| | | |
|---|---|---|
| | | 53,71335-1463-1,71335-1463-2,71335-1463-3,71335-1463-4,71335-1463-5,71335-1463-6,71335-1533-1,71335-1533-2,71335-1533-3,71335-1533-4,71335-1533-5,71335-1533-6,71335-1545-1,71335-1545-2,71335-1545-3,71335-1545-4,71335-1545-5,71335-1545-6,51407-052-30,69784-713-13,69784-714-13,69784-715-13,69784-716-13,51407-053-30,57664-665-83,57664-663-83,57664-664-83,60505-3678-3,60505-3680-3,51407-050-30,51407-051-30,60505-4713-3,60505-4714-3,60505-4715-3,60505-4716-3,16714-227-01,16714-228-01,16714-229-01,16714-230-01,60505-3679-3,60505-3681-3,57664-666-83,63629-8794-1,0007-3383-13,0007-3385-13,0007-3387-13,0007-3388-13,0007-3370-13,0007-3371-13,0007-3372-13,0007-3373-13,0007-3373-61,0007-4139-20,0007-4140-20,0007-4141-20,0007-4142-20 |
| **Esmolol** | NDC | 10019-670-10,10019-666-10,10019-672-10,10019-055-61,10019-075-87,10019-115-01,10019-668-10,0404-9825-10,55150-420-10,55150-421-10,44567-812-10,0404-9858-10,25021-308-84,25021-309-82,25021-314-10,51662-1322-1,44567-811-10,50090-4512-0,50090-4699-0,55150-194-10,51662-1371-1,51662-1427-1,51662-1444-1,71872-7049-1,10019-120-01,68083-211-25,52584-194-10,63323-652-10,67457-182-10,71872-7136-1,67457-657-25,67457-658-10 |
| **Labetalol** | NDC | 71335-0696-4,71335-2063-1,71335-2063-2,71335-2063-3,70518-1546-0,70518-3411-0,50090-1161-1,0615-8131-39,63629-2211-1,63629-2212-1,63629-2213-1,63629-2214-1,63629-2215-1,63629-2216-1,71335-0696-5,0185-0010-01,0185-0010-05,0185-0117-01,0185-0117-05,0185-0118-01,0185-0118-05,49884-122-01,49884-122-05,60760-628-90,49884-123-05,0615-8130-39,71335-0696-2,50090-1161-0,60760-610-90,50090-1290-0,50090-1290-1,50090-1291-0,50090-1291-1,49884-123-01,70518-1546-1,71335-0696-1,71335-0696-3,49884-124-05,70934-913-60,49884-124-01,71335-0733-4,71335-0733-3,71335-1410-1,71335-1410-2,71335-1410-3,71335-1410-4,71335-1410-5,71335-0733-1,71335-0733-2,0143-9366-10,0143-9363-10,0143-9364-10,0143-9365-10,65145-124-01,65145-125-01,68382-798-06,68382-800-05,68382-800-16,63629-2053-1,63629-2054-1,63629-2055-1,47781-552-05,47781-552-01,70771-1163-1,70771-1165-1,0143-9622-01,0143-9623-01,68382-798-05,68382-799-01,71335-2057-1,71335-2057-2,71335-2057-3,47781-553-01,47781-554-01,55154-8192-0,23155-723-01,23155-723-05,23155-724-01,23155-724-05,23155-725-01,23155-725-05,51662-1228-1,71930-035-12,71930-035-52,71930-036-12,71930-036-52,71930-037-12,71930-037-52,70518-2728-0,68382-798-16,68382-799-05,68382-800-06,72266-102-01,72266-103-01,47781-553-05,70518-1163-1,70771-1163-3,70771-1163-9,70771-1164-5,70518-2903-0,72572-350-01,72611-734-01,72611-738-01,42806-327-01,42806-327-05,42806-327-10,42806-328-01,42806-328-05,42806-328-10,42806-329-01,42806-329-05,42806-329-10,70771-1164-1,36000-320-10,36000-322-02,36000-324-02,70771-1165-9,50090-3584-0,71247-126-01,71247-126-05,71247-127-01,71247-127-05,71247-130-01,71247-130-05,50090-4149-0,50090-4149-1,50090-4917-1,70518-1163-2,68083-111-01,0591-0605-01,0591-0605-05,0591-0605-10,0591-0606-01,0591-0606-05,0591-0606-10,0591-0607-01,0591-0607-05,0591-0607-10,51662-1228-9,70771-1164-9,70771-1165-3,70771-1165-5,68083-111-02,60760-231-90,70934-751-60,0904-5928-61,0904-5930-61,68382-799-06,68382-799-16,58657-602-01,58657-602-10,58657-602-50,58657-603-01,58657-603-10,58657-603-50,58657-604-01,58657-604-10,58657-604-50,51407-614-01,51407-614-05,51407-615-01,51407-615-05,51407-616-01,60687-439-01,60687-450-01,60687-461-01,55154-2339-0,0409-2267-20,0409-2267-54,0409-2339-34,72888-120-01,72888-120-05,72888-121-01,72888-121-05,63629-1161-1,63629-1162-1,63629-1163-1,63629-1164-1,63629-1165-1,63629-1166-1,72266-103-41,72888-122-01,72888-122-05,17478-420-20,17478-420-40,68001-381-00,68001-381-03,68001-382-00,68001-382-03,68001-383-00,68001-383-03,68382-800-01,63629-5268-1,63629-5268-2,63629-5268-3,63629-5311-1,63629-5311-2,63629-5311-3,63629-5311-4,63629-5311-5,63629-8898-1,63629-8899-1,63629-8900-1,63629-8901-1,63629-8902-1,63629-8903-1,70771-1163-5,70771-1164-3,0143-9320-01,68382-798-01,71205-095-60,71205-095-90,0904-7109-61,0904-7110-61,0904-7111-61,70377-060-11,70377-060-12,70377-060-13,70377-061-11,70377-061-12,70377-061-13,70377-062-11,70377-062-12,70377-062-13,70518-1163-0,0641-6252-10,71209-083-03,71209-083-11,71209-084-03,71209-084-11,71209-085-03,71209-085-11,71205-095-30,71335-1242-1,71335-1242-2,71335-1242-3,55154-4747-5 |
| **Metoprolol** | NDC | 59212-095-30,10631-011-30,10631-008-30,10631-009-30,10631-010-30,71872-7155-1,63323-660-05,55154-4451-5,55700-232-90,68001-120-00,70518-3164-1,55700-253-90,64679-734-01,64679-734-03,64679-736-03,64679-736-09,71335-0864-6,71335-0864-7,55700-192-30,71335-0864-2,71335-0864-4,68382-564-01,68382-565-05,68382-567-10,68001-356-03,0378-4595-10,0378-4595-77,0378-4596-10,0378-4596-77,0378-4597-10,0378-4597-77,0378-4598-05,0378-4598-77,71610-642-60,71335-1875-6,71335-1878-7,71335-1951-7,50090-2296-0,50090-2296-1,68788-7206-1,68788-7206-2,0615-7825-05,50742-615-05,50742-616-01,50742-616-10,50742-617-05,50742-618-90,68001-122-03,71610-509-45,71610-509-60,71610-512-45,71610-512-60,0904-6322-06,68071-1921-3,68071-1921-6,68071-1921-9,68071-2651-1,55700-192-90,50090-5950-0,50090-5950-1,64679-846-02,64679-846-03,64679-846-10,64679-847-02,64679-847-03,64679-847-10,64679-848-02,64679-848-03,64679-848-10,70771-1341-0,50268-541-15,75834-290-00,75834-290-01,75834-290-05,75834-291-00,75834-291-01,75834-291-05,75834-292-00,75834-292-01,75834-292-05,75834-293-00,75834-293-01,75834-293-05,50742-615-01,68788-7263-2,64679-849-02,64679-849-03,68001-119-03,68001-120-03,68001-121-00,68071-2191-3,68071-2191-6,60760-492-90,50090-5394-0,53002-2781-0,53002-2781-3,71335-0167-7,24979-037-01,24979-037-02,24979-037-03,24979-038-01,24979-038-02,24979-038-03,24979-039-01,24979-039-02,24979-039-03,24979-040-01,50090-3808-1,71610-161-60,53002-2612-0,53002-2612-3,71335-1875-1,71335-1875-2,71335-1875-3,71335-1875-4,71335-1875-5,68788-7263-6,62732-1046-1,71335-0864-1,72516-030-01,72516-030-10,72516-030-50,72516-031-01,72516-031-10,72516-031-50,72516-032-01,72516-032-10,72516-032-50,72516-033-01,71335-1878-1,71335-1878-2,71335-1878-3,71335-1878-4,71335-1878-5,71335-1878-6,68001-357-03,68001-358-03,68001-360-00,50090-5733-0,50090-5733-1,50090-5733-3,71335-1951-1,71335-1951-2,71335-1951-3,71335-1951-4,71335-1951-5,71335-1951-6,31722-589-05,31722-589-10,31722-590-05,31722-590-10,31722-591-05,16714-852-02,16714-854-01,16714-855-01,16714-852-03,16714-853-02,16714-854-03,16714-853-01,60760-493-90,50742-616-90,31722-591-10,31722-592-05,31722-592-10,71335-0786-1,71335-0786-2,71335-0786-3,71335-0786-4,71335-0786-5,71335-0786-6,71335-0786-7,71610-573-45,71610-573-60,70518-2770-0,43063-624-90,68071-2191-9,70518-2868-0,67877-590-01,67877-590-05,67877-590-10,67877-590-30,67877-590-60,67877-591-01,67877-591-05,67877-591-10,67877-591-30,67877-591-60,67877-592-01,67877-592-05,67877-592-10,67877-592-30,67877-592-60,67877-593-01,67877-593-05,67877-593-10,31722-589-01,31722-589-30,31722-590-01,31722-590-30,31722-591-01,31722-591-30,31722- |

| | | |
|---|---|---|
| | | 592-01,31722-592-30,50816-025-02,50816-025-03,50816-050-02,50816-050-03,50816-100-02,50816-100-03,50816-200-02,50816-200-03,68788-8098-1,68788-8098-3,68788-8098-6,68788-8098-9,50742-618-10,67877-593-30,67877-593-60,60760-634-90,45963-676-11,45963-676-96,45963-677-11,45963-677-96,45963-678-11,45963-709-11,45963-709-96,70518-2953-0,70518-0391-0,62732-1046-2,62732-1046-3,62732-1047-1,16714-852-01,16714-855-02,55111-467-78,70436-183-01,70436-183-02,70436-183-03,70518-2133-1,72789-010-30,50090-3808-0,50090-3847-0,50090-3847-1,50090-3847-2,50090-3896-0,50090-3746-1,68382-564-10,50090-4304-0,50090-4304-1,50090-3746-3,53002-1612-0,53002-1612-3,72789-107-90,53002-1781-0,53002-1781-3,72789-145-30,50090-5411-0,50090-5413-0,50090-5053-0,71610-161-45,50742-616-05,63304-009-30,63304-010-30,50090-4902-0,50090-4904-0,71335-0809-1,71335-0809-2,71335-0809-3,71335-0809-4,71335-0809-5,71335-0809-6,68788-7206-6,71335-0149-1,71335-0149-2,71335-0149-3,71335-0149-4,71335-0149-5,71335-0158-1,71335-0158-2,71335-0158-3,71335-0158-4,71335-0158-5,68071-2331-1,50090-5418-0,51079-171-03,68788-7263-3,68071-4351-3,68071-4351-6,68071-4351-9,68071-4363-3,68071-4363-6,68071-4363-9,68071-4377-3,68071-4377-6,68071-4377-9,0615-7823-05,0615-7823-30,0615-7823-39,0615-7824-05,0615-7824-39,0615-7825-39,64679-734-05,64679-735-09,64679-737-02,50090-5393-0,50090-5394-1,50090-5394-3,50090-5395-0,55111-467-60,55111-469-30,50090-5411-1,50090-5412-0,50090-5413-1,50090-5418-1,50090-5419-0,60687-413-01,64679-735-08,71205-368-30,71205-368-60,71205-368-90,64679-737-03,64679-734-08,68001-119-00,68001-121-03,63304-008-30,63304-011-30,55111-467-01,63629-9167-1,63629-9168-1,63629-9169-1,63629-9170-1,63629-9171-1,63629-9172-1,63629-9173-1,68001-469-00,68001-469-08,68001-470-00,68001-470-08,68001-471-00,50268-542-15,55154-4698-0,55154-7279-0,55154-7280-0,16714-853-03,0904-6322-61,0904-6323-61,0904-6324-61,50742-617-01,70771-1340-5,70771-1341-5,61919-334-30,61919-334-90,70771-1338-5,70771-1339-1,70771-1340-0,43063-841-30,68788-7997-1,68788-7997-3,68788-7997-6,68788-7997-9,70518-0391-1,71610-483-45,71610-483-60,71610-484-45,71610-484-60,63187-546-60,68382-565-01,24979-040-09,68645-477-54,55700-192-60,55111-466-78,55111-467-05,55111-468-05,55111-469-05,50742-615-90,50742-617-90,60050-001-01,60050-001-10,60050-002-01,60050-002-10,50090-3746-0,60050-003-01,60050-003-10,60050-004-01,60050-004-10,68788-7263-9,68071-1922-3,68071-1922-6,68071-1922-9,61919-754-90,63187-768-30,63187-768-60,16714-854-02,51407-403-01,51407-403-10,51407-404-01,60687-402-01,51407-404-10,51407-405-01,51407-405-10,51407-406-01,51407-406-10,64679-735-01,64679-735-02,64679-736-05,70771-1339-0,63187-768-90,62732-1047-2,62732-1047-3,63187-547-90,68788-7263-1,50090-5720-0,50090-5720-1,68001-358-00,68001-360-03,50268-540-15,50268-543-15,51655-203-52,63629-8100-1,63629-8863-1,63629-8864-1,63629-8865-1,63629-8866-1,63629-8237-1,63629-8237-2,63629-8237-3,63187-545-30,63187-545-60,63187-545-90,63187-546-30,60687-390-01,71335-0864-5,64679-737-05,68001-122-00,70771-1341-1,63187-546-90,43063-624-30,67296-1149-3,67296-1720-3,70518-3062-0,72789-107-30,64679-734-09,64679-735-05,64679-736-01,64679-736-02,71335-0167-1,71335-0167-2,71335-0167-3,68071-2522-1,68071-5282-1,55111-466-05,55111-466-30,55111-467-30,55111-468-30,50742-617-10,71335-0167-4,71335-0167-5,50742-618-05,71335-0167-6,64679-735-03,64679-737-08,60760-635-90,51079-169-20,71335-0864-3,51407-406-08,68001-500-00,68001-500-03,68788-7206-3,68001-500-08,68001-501-00,68001-501-03,68001-501-08,68001-502-00,68001-502-03,68001-502-08,68001-503-00,71610-496-45,71610-496-60,50742-615-99,50742-616-99,50742-617-99,50742-618-99,70771-1338-1,70771-1339-5,61919-761-30,61919-761-90,71335-1600-1,71335-1600-2,71335-1600-3,71335-1600-4,71335-1600-5,71335-1600-6,71335-1600-7,71335-1600-8,55289-382-90,52817-360-10,50090-2149-1,55154-1557-5,42708-041-60,71335-0743-2,50090-2149-0,62332-112-30,62332-112-91,62332-114-30,63187-532-30,63187-532-72,57237-100-99,71610-548-60,71610-548-80,71610-551-60,71610-551-80,0378-0018-01,0378-0018-05,0378-0032-01,0378-0032-10,0378-0047-01,0378-0047-10,63187-600-90,63187-609-72,55700-646-90,71610-147-70,71610-147-92,71610-147-94,52343-059-01,52343-059-99,0615-8016-05,0615-8016-30,0615-8016-39,0615-8017-30,0615-8017-39,0615-8322-05,0615-8322-30,0615-8322-39,42708-110-60,62332-113-31,62332-114-31,62332-114-91,62332-113-91,65862-062-99,62584-266-01,55289-382-14,55289-382-60,71610-443-60,71610-443-80,71610-443-92,63187-103-78,71610-556-45,63187-314-90,57237-102-01,70518-3291-0,63187-532-90,0143-9873-10,0143-9873-25,70882-133-60,70518-0381-0,71610-443-70,68071-4755-2,71335-0743-1,71335-0743-3,71610-517-53,71610-517-60,71610-517-80,52817-361-00,63187-600-30,68071-1288-3,68071-1288-6,68071-1288-8,68071-1288-9,71610-137-80,43063-821-93,70518-3293-1,43063-821-90,57237-101-99,43353-942-30,50090-1979-0,50090-1979-1,50090-1979-4,50090-1979-5,70882-132-60,55154-2126-0,52817-360-00,52817-362-00,68071-1794-1,68071-1795-1,50090-2149-4,52343-061-99,72266-122-25,57237-100-01,57237-101-01,62332-113-30,62584-265-01,65862-063-01,65862-064-99,43063-927-01,43063-927-14,43063-927-30,43063-927-60,43063-927-90,43063-927-93,43063-938-30,43063-938-60,43063-938-90,43063-938-93,57664-167-59,57664-506-52,57664-506-54,57664-506-59,50090-2149-5,63187-103-90,63187-314-30,57664-166-58,57664-167-52,57664-477-58,57664-506-58,68071-5236-1,63187-174-90,63187-252-30,63187-314-78,49999-575-90,55154-4752-0,62584-267-01,70518-0381-3,46708-290-10,46708-290-30,46708-290-31,46708-290-71,46708-290-91,46708-291-10,46708-291-30,46708-291-31,46708-291-91,46708-292-10,46708-292-30,46708-292-31,46708-292-91,51079-255-20,72611-740-10,72266-122-05,71610-556-30,71610-556-53,71610-556-60,71610-556-70,71610-556-80,71610-556-92,71610-556-98,71610-559-53,71610-559-60,71610-559-80,71610-565-30,71610-565-45,65862-064-60,71335-0743-5,43353-942-98,43353-944-80,67296-1424-3,68071-4014-3,68071-4014-6,68071-4014-8,68071-4014-9,68071-4084-1,50090-1974-0,50090-1974-1,50090-1974-3,50090-1974-4,50090-4966-0,57664-162-58,57664-166-52,50090-5060-0,50090-5070-0,63187-314-60,72888-004-00,72888-004-01,72888-005-00,72888-005-01,72888-006-00,72888-006-01,72888-023-01,72888-023-05,71205-003-78,71610-565-53,71610-565-60,71610-565-70,71610-565-73,71610-565-80,71610-565-92,71610-565-94,71610-565-98,43353-943-98,70934-105-30,68788-7832-1,68788-7832-3,68788-7832-6,68788-7832-9,68071-4261-1,68071-4269-1,68071-4307-9,68071-4309-3,71335-1146-1,71335-1146-2,71335-1146-3,71335-1146-4,71335-1146-5,71335-1146-6,71335-1146-7,71335-1146-8,71335-1157-1,71335-1157-2,71335-1157-3,71335-1157-4,71335-1157-5,68071-4356-3,68071-4356-6,68071-4356-8,68071-4356-9,51655-271-26,71872-7225-1,70518-3293-2,51662-1365-1,51662-1478-1,71610-080-80,71335-0607-1,71335-0607-2,71335-0607-3,71335-0607-4,71335-0607-5,71335-0607-6,71335-0607-7,71335-0607-8,71610-147-60,71610-158-80,50090-0488-0,50090-0488-1,50090-0488-4,50090-0488-5,52817-358-10,52817-358-50,52817-359-10,52817-359-50,43353-942-70,43353-943-92,68071-4747-1,68071-4755-9,53808-1117-1,68071-4759-9,68071-4769-1,68071-4771-9,43353- |

| | | |
|---|---|---|
| | | 942-53,71610-080-60,70518-3293-3,63187-174-30,0904-6340-61,0904-6340-80,0904-6341-80,0904-6342-61,63187-103-30,0904-6342-80,63187-252-78,63187-252-90,68071-4867-9,0378-4593-01,0378-4594-01,68071-4880-9,68071-4957-1,68071-4958-1,71610-443-94,71335-0241-4,58118-1062-8,63187-609-00,63187-609-30,63187-609-90,68645-191-59,63187-103-60,63187-174-60,63187-174-78,43063-821-30,43063-821-60,68071-4307-2,68071-4307-3,68071-4307-6,68071-4307-8,68071-4309-1,68071-4309-8,68071-4309-9,60760-243-60,60760-243-98,60760-295-90,68071-3337-1,68071-3337-3,68071-3337-6,68071-3337-9,60760-553-98,71335-0241-5,70934-736-30,70934-736-60,70934-736-90,70934-736-96,68071-1950-3,68071-1950-6,68071-1950-8,68071-1950-9,61919-052-30,12634-771-00,12634-771-01,12634-771-09,12634-771-12,12634-771-18,12634-771-40,12634-771-42,12634-771-45,61919-317-30,61919-317-60,12634-771-50,12634-771-52,12634-771-54,12634-771-57,12634-771-59,12634-771-60,12634-771-61,12634-771-63,12634-771-66,12634-771-67,12634-771-69,12634-771-71,12634-771-74,12634-771-78,12634-771-79,12634-771-80,12634-771-81,12634-771-82,61919-415-90,12634-771-84,12634-771-85,12634-771-90,12634-771-91,12634-771-92,12634-771-93,12634-771-94,12634-771-95,12634-771-96,12634-771-97,12634-771-98,12634-771-99,68645-190-59,71872-7226-1,67544-387-30,67544-387-53,67544-387-60,67544-387-70,67544-387-80,67544-387-92,67544-387-98,0409-1778-05,71335-0241-1,50090-0489-0,50090-0489-3,43353-942-80,43353-943-30,43353-943-70,61919-415-30,55154-5356-0,55154-5512-0,50090-2207-0,50090-2207-1,50090-2207-2,50090-2207-3,50090-2260-3,50090-2260-4,58118-0361-8,63629-1462-1,63629-1462-2,63629-1462-3,63629-1462-4,63629-1462-5,70518-1354-2,51407-109-01,51407-109-10,51407-110-01,51407-110-10,51407-111-01,51407-111-10,71335-0192-1,71335-0192-2,71335-0192-3,63629-8101-1,63629-8101-2,63629-8101-3,63629-8103-1,63629-8103-2,63629-8103-3,76420-054-20,76420-054-30,71335-0192-7,71335-0192-8,43353-942-92,43353-943-60,72888-101-01,72888-101-05,70518-3293-0,63187-600-60,63187-600-78,63187-609-78,63187-532-60,63187-532-78,50090-0489-1,50090-0489-4,67296-0701-1,67296-0729-1,67296-1667-3,63187-609-60,57664-162-52,71872-7283-1,55154-4745-5,52343-060-99,52817-361-10,52817-362-10,62332-112-31,35356-782-90,57664-162-59,57664-167-58,71610-138-60,71610-147-80,43353-943-73,43353-944-60,60760-553-60,71610-184-60,71610-186-60,65862-063-60,70934-105-96,68788-7338-1,68788-7338-3,68788-7338-6,68788-7338-9,57237-102-99,51079-801-20,68788-7548-1,68788-7548-3,68788-7548-6,68788-7548-9,55154-4790-0,70518-0381-2,71205-003-90,68788-7690-1,68788-7690-3,68788-7690-6,68788-7690-9,61919-428-90,0378-0424-01,0378-0434-01,0378-0445-01,62332-117-30,62332-116-71,62332-117-31,62756-368-08,62756-368-18,62756-368-83,62756-368-88,62756-369-08,62756-369-18,62756-369-83,62756-369-88,62756-370-08,62756-370-18,62756-370-83,62756-370-88,46708-116-30,62332-116-30,62332-116-31,46708-116-50,46708-116-10,46708-117-30,46708-117-50,62332-115-30,62332-115-91,62332-117-71,67544-555-51,67544-491-45,67544-491-80,67544-491-82,71610-373-45,71610-373-60,67544-491-99,70347-025-01,70347-025-02,70347-025-03,70347-050-01,70347-050-02,70347-050-03,70347-100-01,70347-100-02,70347-100-03,70347-200-01,70347-200-02,70347-200-03,71610-038-60,67544-282-51,71610-035-45,67544-491-53,67544-491-60,67544-555-45,67544-555-60,71610-035-60 |
| **Nadolol** | NDC | 27505-100-01, 27505-101-01, 27505-102-01, 78670-100-01, 78670-101-01, 67787-348-30, 67787-349-30, 68382-734-01, 0378-0028-01, 67787-347-10, 0904-7070-07, 0904-7070-61, 0904-7071-07, 0904-7071-61, 10135-686-01, 10135-687-01, 10135-688-01, 68382-732-01, 68382-732-10, 70771-1089-9, 70771-1090-0, 70771-1091-9, 68382-733-16, 70518-1755-0, 23155-730-01, 23155-731-01, 23155-732-01, 68001-319-00, 71335-0697-1, 71335-0697-2, 69238-1123-1, 0781-8005-01, 0781-8005-10, 0781-8006-10, 0781-8004-92, 0781-8006-92, 0781-8004-01, 0781-8005-92, 70771-1090-4, 68001-318-00, 70771-1090-9, 70771-1091-1, 69097-867-07, 69097-869-02, 51079-813-20, 69238-1125-1, 0378-1132-01, 0378-1132-10, 70771-1089-0, 68382-732-16, 68382-734-16, 23155-730-09, 23155-730-10, 23155-731-09, 23155-731-10, 23155-732-09, 23155-732-10, 70771-1091-4, 70771-1089-1, 67787-349-10, 60687-302-25, 70771-1090-1, 69097-869-15, 0378-1171-01, 0378-1171-10, 69097-868-15, 76282-347-01, 76282-348-01, 76282-349-01, 67787-348-10, 69097-868-07, 59651-251-01, 59762-0812-1, 76385-133-01, 76385-133-10, 76385-134-01, 76385-134-10, 76385-135-01, 76385-135-10, 71335-1972-1, 71335-1972-2, 67787-347-30, 71335-1972-3, 59651-252-01, 69097-869-07, 51079-812-20, 59762-0810-1, 69238-1124-1, 69097-868-02, 68382-732-77, 68382-734-77, 63629-6942-1, 72664-211-01, 72664-212-01, 72664-213-01, 70518-2139-0, 68001-317-00, 69097-867-02, 69097-867-15, 70771-1089-4, 70771-1091-0, 68382-733-10, 59762-0811-1, 0781-8004-10, 0781-8006-01, 71335-0697-3, 68382-733-01, 60687-313-25, 68382-733-77, 68382-734-10 |
| **Nebivolol** | NDC | 50090-3326-0, 50090-1127-0, 50090-1127-1, 50090-1307-0, 50090-1307-1, 0456-1402-01, 0456-1402-30, 0456-1402-63, 0456-1402-90, 0456-1405-01, 0456-1405-30, 0456-1405-63, 0456-1405-90, 0456-1410-01, 0456-1410-30, 0456-1410-63, 0456-1410-90, 0456-1420-01, 0456-1420-30, 0456-1420-63, 0456-1420-90, 55154-4623-2, 55154-4622-8, 55154-4621-2, 71209-058-01, 71209-058-04, 71209-058-05 ,71209-058-11, 71209-059-01, 71209-059-04, 71209-059-05, 71209-059-11, 71209-060-01, 71209-060-04, 71209-060-05, 71209-060-11, 71209-061-01, 71209-061-04, 27241-179-30, 27241-180-30, 27241-180-90, 27241-181-30, 27241-181-90, 27241-182-30, 27241-182-90, 51407-486-30, 51407-486-90, 0904-7189-04, 0904-7190-04, 13668-353-01, 62559-278-05, 43547-524-03, 43547-524-09, 43547-524-50, 43547-525-03, 43547-525-09, 43547-525-50, 43547-527-03, 43547-527-09, 43547-527-50, 71209-061-05, 71209-061-11, 0904-7225-04, 0904-7226-04, 59651-137-30, 59651-137-90, 59651-138-30, 72241-035-05, 72241-035-11, 50090-5708-0, 50090-5708-1, 50090-5761-0, 50090-5761-1, 50090-5761-2, 72241-033-11, 72241-033-22, 72241-034-04, 72241-034-05, 72241-034-11, 72241-034-22, 72241-035-04, 72241-035-22, 59651-138-90, 59651-139-30, 59651-139-90, 59651-140-30, 59651-140-90, 68462-615-01, 68462-615-11, 68462-615-30, 68462-616-01, 68462-616-11, 68462-616-30, 68462-617-01, 68462-617-11, 68462-617-30, 68462-618-01, 68462-618-11, 68462-618-30, 72241-032-04, 72241-032-05, 72241-032-11, 72241-032-22, 72241-033-04, 72241-033-05, 13668-353-05, 13668-353-30, 13668-353-90, 13668-354-01, 13668-354-05, 13668-354-30, 13668-355-01, 13668-355-05, 13668-355-30, 13668-356-01, 13668-356-05, 13668-356-30, 13668-354-90, 13668-355-90, 13668-356-90, 50090-5752-0, 50090-5752-1, 67877-390-01, 67877-390-30, 67877-390-90, 67877-391-01, 7877-391-30, 67877-391-90, 67877-392-01, 67877-392-30, 67877-392-90, 67877-393-01, 67877-393-30, 67877-393-90, 62559-275-30, 62559-275-83, 62559-276-30, 62559-276-71, 62559-276-90, 62559-277-05, 62559-277-30, 62559-277-90, 62559-278-30, 62559-278-90, 31722-585-01, 31722-585-30, 31722-585-32, 31722-585-34, 31722-586-01, 31722-586-30, 31722-586-32, 31722-586-34, 31722-586-90, 31722-587-01, 31722-587-30, 31722-587-32, |

| | | |
|---|---|---|
| | | 31722-587-34, 31722-587-90, 31722-588-01, 31722-588-30, 31722-588-32, 31722-588-34, 31722-588-90, 51407-483-30, 51407-484-30, 51407-484-90, 51407-485-30, 51407-485-90, 50090-5775-0, 50090-5775-1, 50090-5775-2, 29300-375-05, 29300-375-13, 29300-375-19, 29300-376-05, 29300-376-13, 29300-376-19, 29300-377-05, 29300-377-13, 29300-377-19, 29300-378-05, 29300-378-13, 29300-378-19 |
| **Penbutolol** | NDC | 0091-4500-15 |
| **Pindolol** | NDC | 0378-0052-01, 0378-0127-01, 70710-1063-1 70710-1064-1, 62559-561-01, 72789-099-01, 72789-100-01, 70771-1134-1, 70771-1135-1, 57664-656-88, 76385-131-01, 76385-132-01, 62559-560-01, 29033-028-01, 29033-029-01, 57664-655-88 |
| **Propanolol** | NDC | 64370-375-50, 64370-375-01, 62559-522-01, 62559-523-01, 62559-521-01, 62559-520-01, 62559-600-30, 62559-601-77, 62559-601-30, 62559-600-77, 62559-600-14, 62559-601-14, 62559-590-77, 62559-590-30, 62559-591-30, 62559-590-14, 62559-591-14, 62559-591-77, 71872-7195-1, 63323-604-01, 61919-794-30, 61919-488-30, 68382-163-05, 0115-1659-01, 0115-1659-03, 0115-1660-01, 0115-1660-03, 0115-1661-01, 0115-1661-03, 0115-1662-01, 0115-1662-02, 0115-1693-01, 55289-233-12, 70518-0022-0, 71335-1712-1, 71335-1712-2, 71335-1712-3, 71335-1712-4, 71335-1712-5, 71335-1712-6, 71335-1712-7, 42291-522-01, 42291-524-01, 68382-161-77, 68382-162-05, 71335-1765-1, 71335-1765-2, 71335-1765-3, 71335-1765-4, 65841-747-10, 0527-4116-37, 0527-4116-41, 0527-4117-37, 0527-4117-41, 0527-4118-37, 0527-4119-37, 0527-4119-41, 63629-2069-1, 63629-2070-1, 63629-2071-1, 63629-2072-1, 53002-4951-0, 53002-4951-3, 0378-0182-01, 0378-0182-10, 0378-0183-01, 0378-0183-10, 0378-0184-01, 0378-0184-10, 0378-0185-01, 0378-0185-05, 0378-0187-01, 0615-8134-39, 63187-839-30, 63187-839-60, 0904-6705-06, 51991-819-01, 71610-339-60, 71335-0907-2, 62559-530-01, 70518-0309-0, 61919-808-30, 68382-161-05, 68382-162-10, 68382-163-01, 68382-163-16, 68382-164-01, 70518-3377-0, 0615-8426-39, 60687-226-01, 70518-1041-0, 71335-0832-1, 71335-0832-2, 71335-0832-3, 71335-0832-4, 50090-3296-6, 50268-700-15, 50268-701-15, 0143-9872-10, 70518-3281-0, 55289-233-60, 68382-163-77, 65841-745-10, 65841-745-16, 65841-745-77, 65841-746-10, 65841-746-77, 65841-747-05, 65841-748-10, 71335-0234-5, , 1335-0234-6, 68788-7860-3, 51991-818-01, 51991-820-05, 60687-587-01, 60687-598-01, 60687-609-01, 71335-0082-1, 71335-0082-2, 71335-0082-3, 71335-0082-4, 71335-0082-5, 71335-0082-6, 71335-0082-7, 71335-0472-1, 71335-0472-2, 71335-0472-3, 71335-0472-4, 71335-0472-5, 71335-0472-6, 71335-0472-7, 71335-0554-1, 71335-0554-2, 71335-0554-3, 71335-0554-4, 42291-522-10, 50090-1501-0, 50268-702-15, 72189-018-30, 72189-074-30, 68382-161-10, 68382-162-77, 50090-2829-0, 65841-747-01, 51991-817-05, 51991-818-05, 51991-820-01, 71335-1712-8, 70518-0287-1, 62559-532-01, 0615-8413-39, 52584-872-10, 70518-0833-0, 70934-922-90, 55700-940-30, 0904-6705-61, 71335-0955-1, 62559-533-01, 0228-2778-11, 0228-2779-11, 0228-2780-11, 0228-2781-11, 65841-745-01, 65841-746-16, 65841-747-16, 65841-747-77, 70518-0164-0, 70518-1463-0, 55289-233-30, 50090-3296-1, 60687-215-01, 50090-3953-1, 50090-3953-6, 62559-531-01, 71335-1994-1, 71335-1994-2, 71335-1994-3, 71335-1994-4, 71335-1994-5, 71335-1994-6, 60429-128-05, 60429-129-05, 0603-5482-21, 0603-5482-32, 0603-5483-21, 0603-5483-32, 0603-5484-21, 0603-5484-32, 0603-5485-21, 0603-5486-21, 0603-5486-28, 51407-235-01, 51407-235-10, 51407-236-01, 51407-236-10, 51407-237-01, 51407-237-10, 51407-238-01, 51407-238-05, 51407-239-01, 71335-0234-4, 51655-348-26, 0615-8417-39, 72189-322-30, 51662-1338-1, 0591-5554-01, 0591-5554-10, 0591-5555-01, 0591-5555-10, 0591-5556-01, 0591-5556-10, 0591-5557-01, 0591-5557-05, 50090-0146-1, 50090-0146-5, 72189-322-90, 68071-4677-9, 68071-4794-3, 53808-1126-1, 0904-6550-61, 71335-0234-2, 68071-5071-1, 63629-1968-1, 68788-6855-3, 68071-4573-3, 68071-4573-6, 68071-4573-9, 60760-262-60, 60760-538-30, 53002-4561-0, 53002-4561-1, 53002-4561-3, 42291-525-01, 17856-3728-1, 70518-3053-0, 43353-084-60, 71335-0234-1, 71335-0661-1, 71335-0661-2, 71335-0661-3, 71335-0661-4, 71335-0661-5, 71335-0661-6, 71335-0955-3, 60429-126-05, 0615-8186-39, 63629-2276-1, 71335-0907-1, 0404-9944-01, 63629-3587-1, 63629-3587-2, 63629-3587-3, 63629-3587-4, 60429-127-05, 43063-647-30, 43063-647-90, 71205-590-30, 71205-590-60, 71205-590-90, 71335-0955-4, 69238-2077-1, 69238-2077-7, 69238-2078-1, 69238-2078-7, 69238-2079-1, 69238-2079-7, 69238-2080-1, 69238-2081-1, 69238-2081-5, 68071-2484-3, 68071-2484-9, 50090-0420-0, 66267-261-90, 63629-2274-1, 63629-2275-1, 63629-2277-1, 63629-2278-1, 63629-2279-1, 63629-2280-1, 63629-2281-1, 63629-2282-1, 63187-839-03, 71335-0731-1, 42291-523-10, 71335-0907-4, 0527-4118-41, 71335-0955-2, 71335-0907-3, 68382-162-01, 68382-163-10, 43353-961-60, 65841-746-01, 65841-748-01, 65841-748-16, 68788-7350-1, 68788-7350-3, 68788-7350-6, 68788-7350-9, 60687-306-01, 60760-545-60, 68788-7791-3, 0615-8187-39, 71610-339-45, 69292-530-01, 69292-530-10, 69292-532-01, 69292-532-10, 69292-534-01, 69292-534-10, 69292-536-01, 69292-536-10, 69292-538-01, 69292-538-10, 69292-538-50, 70518-3141-0, 63187-839-90, 51655-426-26, 51655-426-52, 55289-233-90, 63629-2882-1, 71872-7096-1, 70518-0164-1, 68071-2424-1, 70518-0833-2, 70518-1761-0, 68382-164-05, 68382-164-10, 68382-164-16, 70518-1831-0, 70518-2429-0, 70518-2464-0, 70518-3102-0, 70518-3377-1, 55289-233-01, 68382-161-01, 68382-161-16, 0121-0908-40, 71205-185-30, 71205-185-60, 71205-185-90, 71205-446-30, 71205-446-60, 71205-446-90, 71335-0731-2, 71205-499-30, 71205-499-60, 1205-499-90, 42291-523-01, 65841-745-05, 65841-746-05, 65841-748-05, 65841-748-77, 71335-0661-7, 71335-0907-5, 71335-0731-4, 71335-1120-1, 71335-1120-2, 71335-1120-3, 71335-1120-4, 71335-1120-5, 71335-1120-6, 71335-1120-7, 68382-162-16, 68382-164-77, 71335-1140-1, 71335-1140-2, 71335-1140-3, 71335-1140-4, 51991-817-01, 51991-819-05, 42291-524-10, 42291-525-10, 71335-0234-3, 71335-0731-3, 53002-3600-1, 53002-3600-3, 53002-4560-0, 53002-4560-3, 53002-4950-0, 53002-4950-3, 60687-295-01, 0054-3727-63, 0054-3730-63, 61919-489-30 |
| **Sotalol** | NDC | 70515-105-10, 70515-109-10, 13672-051-00, 13672-052-00, 13672-053-00, 70515-106-10, 70515-116-06, 70515-119-06, 13672-054-00, 13672-055-00, 13672-056-00, 70515-115-06, 0245-0012-01, 0245-0012-11, 0245-0013-01, 0245-0013-11, 0245-0014-01, 0245-0014-11, 0245-0015-01, 0245-0015-11, 71610-385-80, 76385-116-01, 10135-661-50, 10135-663-01, 10135-661-01, 10135-662-01, 76385-115-50, 76385-115-01, 76385-114-01, 76385-114-50, 76385-116-50, 71335-1189-1, 71335-1189-2, 71335-1189-3, 60505-0082-1, 71335-0260-3, 69724-112-10, 50090-1299-0, 71610-294-30, 71610-294-60, 10135-715-01, 10135-716-01, 10135-717-01, 71610-474-80, 60429-748-05, 50090-1299-1, 70518-3341-0, 60429-750-01, 71335-0260-2, 72789-135-01, 72789-136-01, 72789-137-01, 72789-138-01, 63187-426-60, , 3629-2421-1, 63629-2422-1, 63629-2423-1, 63187-804-30, 71610- |

| | | |
|---|---|---|
| | | 474-60, 55154-8179-0, 63187-804-90, 60505-0080-1, 60505-0224-1, 60505-0080-0, 71610-074-60, 71335-1917-1, 71335-1917-2, 71335-1917-3, 60429-748-01, 50268-724-15, 50268-725-15, 63187-426-30, 63187-426-90, 71335-0260-1, 42806-123-01, 76385-125-01, 76385-125-50, 76385-126-01, 76385-126-50, 76385-127-01, 76385-127-50, 60505-0081-1, 60505-0159-0, 60505-0224-2, 60429-749-01, 71335-0860-3, 42806-121-10, 42806-122-01, 42806-122-10, 42806-123-10, 43353-616-60, 60505-0159-1, 60505-0222-2, 60505-0223-1, 50090-1299-2, 63187-804-60, 71205-046-30, 0904-7143-61, 69584-841-10, 69584-841-50, 69584-842-10, 69584-842-30, 69584-843-10, 69584-844-10, 71335-0860-2, 68084-654-01, , 0505-0222-1, 60505-0223-2, 43353-616-30, 60429-751-01, 60505-0081-0, 60505-0082-0, 71335-0860-1, 42806-121-01, 71205-046-60, 71205-046-90, 71335-1125-1, 71335-1125-2, 0093-1060-01, 0093-1061-01, 0093-1062-01, 0093-1063-01, 71610-074-30, 24338-530-25, 24338-530-48 |
| **Timolol** | NDC | 76478-001-05, 76478-002-10, 76478-002-15, 76478-002-05, 76478-002-12, 0781-7186-70, 0781-7186-75, 0781-7186-85, 60505-6225-2, 60505-6225-0, 60505-6225-1, 0023-9211-03, 0023-9211-05, 0023-9211-10, 0023-9211-15, 17478-605-10, 17478-604-30, 69315-305-05, 69315-305-10, 17478-514-11, 42571-382-27, 42571-382-73, 50383-233-10, 60429-115-10, 50383-261-61, 50383-261-91, 50383-233-05, 65862-946-01, 70069-051-12, 61314-030-01, 61314-030-02, 50090-1247-0, 24208-486-05, 24208-486-10, 42571-147-26, 62332-553-10, 65862-947-18, 65862-947-60, 70377-082-11, 14445-405-10, 0527-1763-73, 24208-004-01, 24208-004-02, 24208-004-03, 61314-224-05, 61314-225-05, 61314-225-25, 61314-224-25, 67877-229-11, 0378-0715-01, 61314-226-05, 61314-227-05, 73152-029-01, 64980-513-05, 64980-514-01, 0378-0055-01, 0378-0221-01, 73152-030-01, 73152-031-01, 17478-189-24, 67877-229-15, 45865-121-01, 61314-226-15, 24208-818-25, 24208-819-05, 50090-3441-0, 68682-045-50, 50090-5091-0, 24208-496-05, 50090-0558-0, 82260-496-05, 60505-1005-1, 50383-021-10, 67877-229-55, 60758-801-10, 60758-802-05, 60758-802-10, 68682-045-25, 50383-021-05, 50383-021-15, 60505-1005-4, 63629-7167-4, 17478-288-10, 17478-288-11, 17478-288-12, 17478-288-25, 17478-289-10, 17478-289-11, 17478-289-12, 17478-289-25, 17478-365-05, 17478-366-05, 17478-366-10, 17478-366-15, 68682-812-05, 68682-813-05, 68682-813-10, 63629-7167-1, 63629-7167-2, 63629-7167-3, 61314-227-10, 61314-227-15, 61314-226-10, 64980-513-01, 64980-513-15, 64980-514-15, 60758-801-05, 64980-514-05, 50090-5769-0, 62332-545-05, 62332-546-05, 24208-812-05, 24208-813-05, 24208-813-10 ,69918-601-60, 69918-602-60, 0187-1496-05 ,0187-1496-99, 0187-1498-25, 24208-498-34, 24208-499-00, 24208-499-68, 24208-814-25, 24208-816-05 |
